# Large-scale imputation models for multi-ancestry proteome-wide association analysis

**DOI:** 10.1101/2023.10.05.561120

**Authors:** Chong Wu, Zichen Zhang, Xiaochen Yang, Bingxin Zhao

## Abstract

Proteome-wide association studies (PWAS) decode the intricate proteomic landscape of biological mechanisms for complex diseases. Traditional PWAS model training relies heavily on individual-level reference proteomes, restricting its capacity to harness the emerging summary-level protein quantitative trait loci (pQTL) data in the public domain. Here we introduced BLISS, a novel framework to train protein imputation models using only pQTL summary statistics. By leveraging extensive pQTL data from the UK Biobank, deCODE, and ARIC studies, we applied BLISS to develop large-scale European PWAS models covering 5,779 unique proteins. We further extended BLISS to integrate with small-scale non-European individual-level datasets, enabling the development of models tailored to Asian and African ancestries. We validated the performance of BLISS models through a systematic multi-ancestry analysis of over 2,500 phenotypes across five major genetic data resources. The newly identified protein-phenotype associations offer valuable insights into their cross-ancestry transferability, the contributions of different proteomic platforms, and the complementary perspectives provided by distinct genomic mapping approaches relevant to drug discovery. The developed models and data resources are freely available at https://www.gcbhub.org/.

## Introduction

Human proteome provides unique insights into human biology and disease^1-3^. Large-scale data resources and computational approaches have been developed to link genomics and proteomics, identifying protein quantitative trait loci (pQTL) where genetic variations affect protein levels^4-14^. To identify potential therapeutic targets and empower drug discovery, it is of great interest to integrate the genomic profile of proteins with the genetic architecture of complex traits depicted in genome-wide association studies (GWAS)^15,16^. Proteome-wide association studies (PWAS) have emerged as a promising approach to identify proteome-phenotype connections using genetically imputed proteomic models and GWAS summary statistics^17-19^, quickly led to the widespread usage to study complex traits, including stroke^20^, Alzheimer’s disease^19,21^, depression^22,23^, post-traumatic stress disorder (PTSD)^24,25^, neuropsychiatric and substance use disorders^26,27^, and multiple sclerosis^28^, aiding in the mapping of genetic causal pathways and identifying druggable targets. Nevertheless, most current PWAS have focused on a limited set of disease outcomes. While these studies have uncovered some disease-specific findings, comprehensive data resources and insights into the multi-ancestry landscape of protein-phenotype associations remain lacking.

Due to data privacy restrictions and logistical challenges, the genetic community has endorsed the practice of sharing summary-level pQTL data rather than individual-level proteomic and genetic data. This poses a challenge for traditional PWAS models, which require access to individual-level reference proteomes for training. Consequently, existing PWAS models are typically trained on a relatively small reference proteomic dataset, often encompassing just a few hundred participants^17-19^. Furthermore, the constraints of small sample sizes (and consequently reduced statistical power) in PWAS model training have restricted the potential of large-scale PWAS applications, particularly when scrutinizing massive phenotypes across curated GWAS databases, given the heavy multiple testing burden. As a result, many important questions remain difficult to answer. For example: Do PWAS findings differ across ancestries? Do different proteomic platforms yield consistent protein-phenotype associations? How do PWAS results compare to those from other approaches, such as transcriptome-wide association studies (TWAS), Mendelian randomization (MR), or Bayesian colocalization analyses? And do these PWAS findings have the potential to identify viable drug targets?

To address these questions, we introduced BLISS, a statistical framework for protein imputation model training using only pQTL summary statistics and linkage disequilibrium (LD) reference panel. BLISS directly address the practical challenges posed by the rapidly evolving and heterogeneous landscape of proteomic platforms, as well as the often-restricted access to matched individual-level data required by other methods such as SUMMIT^29^ for hyperparameter tuning (**Fig. 1A**). We applied this pipeline to summary-level pQTL data from the UK Biobank^4^ (UKB, *n* = 49,341 participants), deCODE genetics^7^ (*n* = 35,892), and Atherosclerosis Risk in Communities study^8^ (ARIC, *n* = 7,213 for European and 1,871 for African ancestries) (**Fig. 1B**). Additionally, we leveraged large-scale European summary-level pQTL data to improve prediction accuracy in non-European populations, utilizing their publicly available individual-level data from the UKB study (*n* = 914 for Asian and 1,171 for African ancestries). We applied BLISS models to curated GWAS datasets and conducted a systematic multi-ancestry analysis to address the questions outlined above, resulting in a comprehensive atlas of multi-ancestry protein-phenotype associations (**Figs. 1C-1D**). The developed models and web tool are freely accessible at GCB Hub (https://www.gcbhub.org/), covering over 2,500 phenotypes by using GWAS summary statistics from the Million Veteran Program^30^ (MVP), Global Biobank Meta-analysis Initiative^31^ (GBMI), FinnGen^32^, IEU OpenGWAS^33^, and Biobank Japan^34^ (BBJ) projects.

**Fig. 1.**
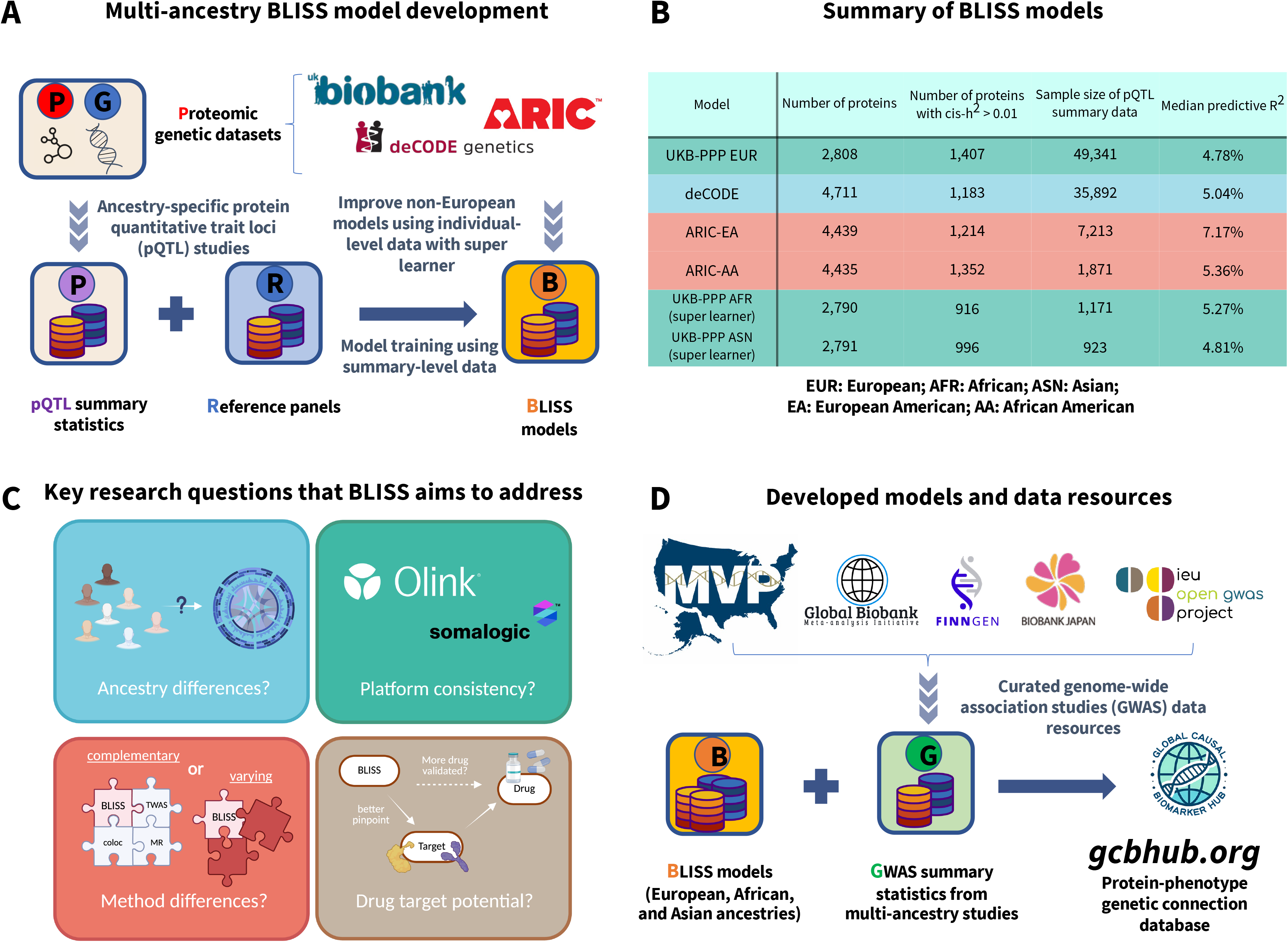
Study design and model performance. (A) Schematic of the BLISS model training pipeline combining proteomic and genetic datasets from biobanks with ancestry-specific pQTL summary statistics and reference panels. (B) Performance characteristics of BLISS models, including protein counts, sample sizes, and median predictive. (C) Key research questions addressed by BLISS regarding ancestry differences, platform consistency, methodological comparisons, and drug target identification. (D) Application workflow combining BLISS models with GWAS summary statistics from five major databases to generate protein-phenotype associations on the GCBHub platform.

## Results

### Development of BLISS models

We introduced Biomarker expression Level Imputation using Summary-level Statistics (BLISS), a novel method designed for developing PWAS-type protein imputation models using summary-level pQTL data. BLISS used *cis*-single nucleotide polymorphisms (*cis*-SNPs, within 1 megabase of the transcription start and end sites) as its predictors. Using a penalized regression framework, protein imputation models were trained on summary-level pQTL data alongside a LD reference panel. For model parameter tuning, BLISS selected parameters by maximizing the predictive R-squared (*R*^2^) in a pseudo-tuning setup^35^ using only summary-level data. BLISS used a robust pseudo-ensemble learning approach to combine prediction models generated with varying LD clumping parameters (*r*^2^ thresholds of 0.01, 0.05, 0.1, 0.2, 0.5, and 0.9). This was achieved by resampling pseudo-training, tuning, and ensemble learning datasets conditional on summary-level pQTL data (Methods). The fully summary statistics-based approach distinguishes BLISS from previous methods such as SUMMIT^29^. While SUMMIT accepts summary-level training data, it still relies on individual-level tuning data for hyperparameter selection. This requirement becomes increasingly problematic due to the rapid advancement of proteomics technologies and growing concerns over data privacy.

We applied BLISS to plasma proteins from two platforms: (i) SomaScan v4, an aptamer-based multiplex protein assay; and (ii) Olink Explore 3072, an antibody-based proximity extension assay. Specifically, for European ancestry, we trained models using summary-level pQTL data of 35,892 Icelanders from the deCODE project^36^ measured via the SomaScan platform, 7,213 European American (EA) individuals from the ARIC^8^ project also measured via the SomaScan platform, and 49,341 White British individuals from the UK Biobank Pharma Proteomics Project^4^ (UKB-PPP) measured by the Olink platform. For African ancestry, our models were trained using data from 1,871 African American (AA) ARIC individuals measured via the SomaScan platform. We evaluated the prediction accuracy of proteins with an estimated *cis*-heritability greater than 0.01^8,19^, including 1,407 proteins for UKB-PPP (European), 1,183 for deCODE, 1,214 for ARIC-EA, and 1,352 for ARIC-AA. Of these models, the median estimated predictive *R*^2^ values were 4.78%, 5.04%, 7.17%, and 5.36% for UKB-PPP (European), deCODE, ARIC-EA, and ARIC-AA, respectively (**Fig. 1B** and **Table S1**). Notably, the BLISS-trained ARIC-EA and ARIC-AA models had predictive *R*^2^ values consistent to those of the original ARIC-EA and ARIC-AA PWAS models trained on individual-level data (**Fig. S1**).

To evaluate the reliability of the predictive *R*^2^ derived from BLISS, we compared it with the individual-level predictive *R*^2^ estimated from hold-out UKB-PPP data. The individual-level predictive *R*^2^ was calculated in an independent UKB testing dataset consisting of 2,932 White non-British individuals. We found that the summary-level predictive *R*^2^ values from the BLISS closely aligned with those computed from the individual-level testing data (**Fig. 2A**, correlation = 0.99), indicating that BLISS-derived *R*^2^ values were reliable estimates for predictive performance. This was further supported by the high concordance (**Fig. S2**, correlation = 0.97) between the predictive *R*^2^ values and previously reported *cis*-heritability estimates4. As expected, predictive *R*^2^ showed moderate correlations across distinct cohorts and platforms (**Fig. S3**), likely reflecting a combination of shared underlying genetic components and known differences in platform technologies and cohort-level features such as sample size.

**Fig. 2.**
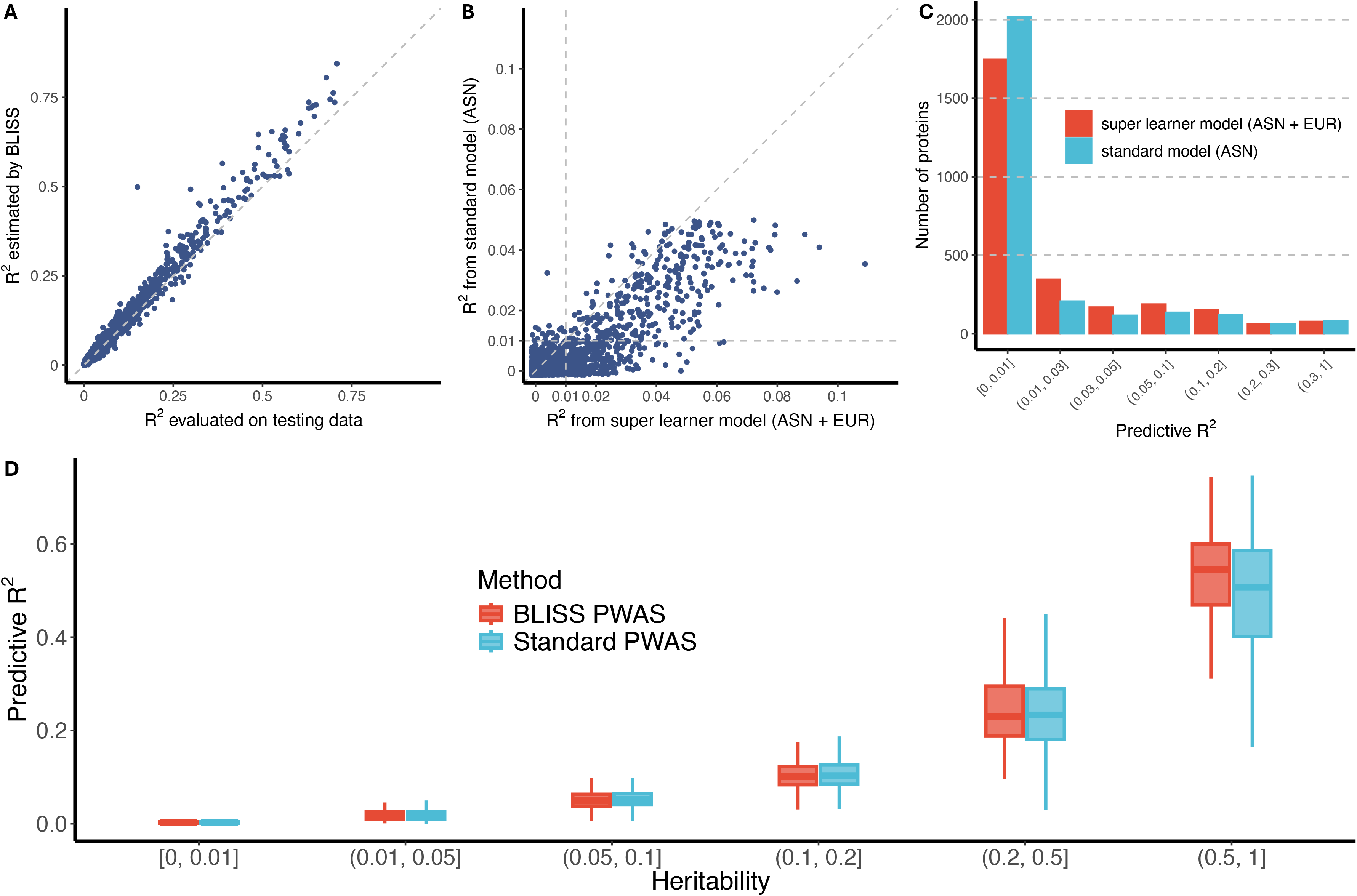
BLISS model validation and performance comparison in real data. (A) Correlation between BLISS internal *R*^2^ estimates using summary-level pseudo-testing data and *R*^2^ evaluated on independent individual-level testing data for UKB-PPP European (EUR) models. (B) Improvement in predictive *R*^2^ achieved by super learner enhancement over standard UKB-PPP Asian (ASN) ancestry models. Each point represents one protein; dashed lines indicate *R*^2^ = 1%. (C) Distribution of proteins across predictive *R*^2^ bins for standard (ASN) and super learner (ASN + EUR) models. (D) Performance comparison across methods stratified by heritability bins. Standard PWAS models used Elastic-net on individual-level data.

We found that the predictive *R*^2^ is robust to the choice of LD reference panel in the BLISS framework. To understand the role of reference panel sample size, we conducted experiments varying the size of the LD reference panel of 503 (matching the sample size of 1000 Genomes reference data) or 49,341 UKB individuals. Larger reference panels (*n* = 49,341) outperformed smaller ones (*n* = 503) and achieved 10% average higher predictive *R*^2^ (0.065 vs. 0.059, **Fig. S4**). When the LD sample size is small, the models derived from UKB LD reference panel with randomly selected 503 subjects and 1000 Genomes LD reference panel were highly similar: the correlation between predictive *R*^2^ is 0.97 (**Fig. S5**) and the number of proteins with predictive *R*^2^ > 0.01 were 1,209 and 1,214, respectively. These results suggest that BLISS does not require direct alignment between the LD reference panel and pQTL samples. Moreover, both deCODE and ARIC models demonstrated consistent performance across different LD reference panels (**Fig. S6**). Correlation between *R*^2^ was over 0.97, and the number of proteins with predictive *R*^2^ > 0.01 was also similar (deCODE (UKB LD): 1,145; deCODE (1000 Genomes LD): 1,153; ARIC (UKB LD):1,129; ARIC (1000 Genomes LD): 1,120). As expected, when these models were applied to GWAS datasets in later section, we observed high concordance in association results that were identified by at least one approach (*Z*-score correlations around 0.99 for all associations, **Fig. S7**). Furthermore, we demonstrated that combining models from multiple LD clumping *r*^2^ thresholds enhanced predictive accuracy compared to models utilizing a single LD clumping threshold (for example, 0.2, **Fig. S8**).

The UKB-PPP also includes protein data for 1,171 individuals of African ancestry and 923 individuals of Asian ancestry. Leveraging these publicly available individual-level data, we investigated strategies to improve prediction accuracy in non-European populations by combining European-trained BLISS models with limited target population data. Notably, directly applying European models to non-European individuals resulted in substantially reduced predictive performance compared to models trained on ancestry-matched pQTL data, despite the much larger European training sample sizes (**Fig. S9**). To address this limitation, we developed a super learner framework that integrates ancestry-specific imputation models with European BLISS models (Methods). We found that the predictive performance of these super learner-enhanced models was generally higher compared to using standard PWAS models that trained on individual-level pQTL alone in both Asian and African ancestries in UKB-PPP, especially for proteins that had low prediction accuracy in standard models (**Figs. 2B-2C** and **S10**). For Asian models with a predictive *R*^2^ below 5%, the median predictive *R*^2^ increased 3.6-fold from 0.31% to 1.32% when enhanced with super learner (one-sided Wilcoxon rank test, *P* < 2.2 × 10^−16^). The super learner models increased the number of proteins with an *R*^2^ exceeding 1% from 726 to 996. For African models, a similar improvement was observed, albeit more modest, potentially attributed to the closer genetic relationship between Europeans and Africans as compared to between Europeans and Asians. Specifically, among the African models with a predictive *R*^2^ under 5%, the median predictive *R*^2^ was 0.84% for super learner models, whereas it was 0.28% for standard models (*P* < 2.2 × 10^−16^). Additionally, to rigorously evaluate the estimates from our nested cross-validation procedure accurately reflect true out-of-sample prediction accuracy, we conducted an independent validation analysis by randomly partitioning ancestry-specific individual-level data into independent training (80%) and testing (20%) sets. We found that the predictive *R*^2^ estimates from this hold-out testing set closely matched those inside the super learner (**Fig. S11**). Furthermore, we investigated the factors that driving the efficacy of super learner integration. Proteins exhibiting high transferability gain (>10% improvement in *R*^2^ by super learner) were characterized by two features: (i) smaller inter-ancestry differences in local LD patterns, and (ii) greater overlap of lead pQTLs between ancestries (**Fig. S12**).

In UKB-PPP, we also observed overall consistency between BLISS models and standard PWAS models trained on individual-level data across proteins spanning all heritability bins (**Fig. 2D**). Notably, BLISS models showed better predictive accuracy compared with standard PWAS models when the heritability was high. These results suggest that BLISS can train models using only summary-level pQTL data without compromising prediction accuracy, providing substantial practical advantages and convenience. In summary, our multi-ancestry model development, leveraging up-to-date pQTL data resources, marks a substantial advance in scope and potential analytical power over previously available PWAS models, which were typically more restricted in ancestral diversity and data scale (Supplementary Note).

### The performance of BLISS in simulations

We conducted extensive simulations to evaluate the performance of BLISS, varying protein expression heritability, pQTL training sample sizes, and phenotypic heritability (Methods). Using a sample size (*n* = 45,000) close to our real-data sample sizes (*n* = 49,341) and 5% of genetic variants to be causal, we assessed the predictive *R*^2^of BLISS models constructed using summary-level pQTL data relative to standard PWAS models constructed using individual-level data (**Fig. 3A**). BLISS achieved comparable predictive *R*^2^ to standard PWAS models at lower protein expression heritability values 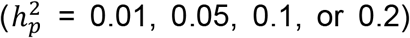 and achieved higher predictive *R*^2^ at higher heritability values (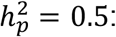 median *R*^2^, 0.467 vs. 0.446, *P* < 2.2 × 10^−16^ by one-sided Wilcoxon rank test; 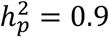 : median *R*^2^, 0.855 vs. 0.780, *P* < 2.2 × 10^−16^). We subsequently evaluated the statistical power to detect protein-phenotype associations using GWAS summary data for simulated complex traits, comparing BLISS with standard PWAS models. BLISS demonstrated comparable or better statistical power to standard PWAS across varying protein expression heritability levels. For example, at intermediate protein expression heritability 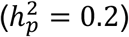, BLISS achieved power of 0.80 versus 0.72 for standard PWAS, approaching the theoretical upper bound of 0.85 using the true protein models without genetic imputation (**Fig. 3B**). Similar patterns were observed in further simulations across different parameter settings (**Figs. S13-S14**).

**Fig. 3.**
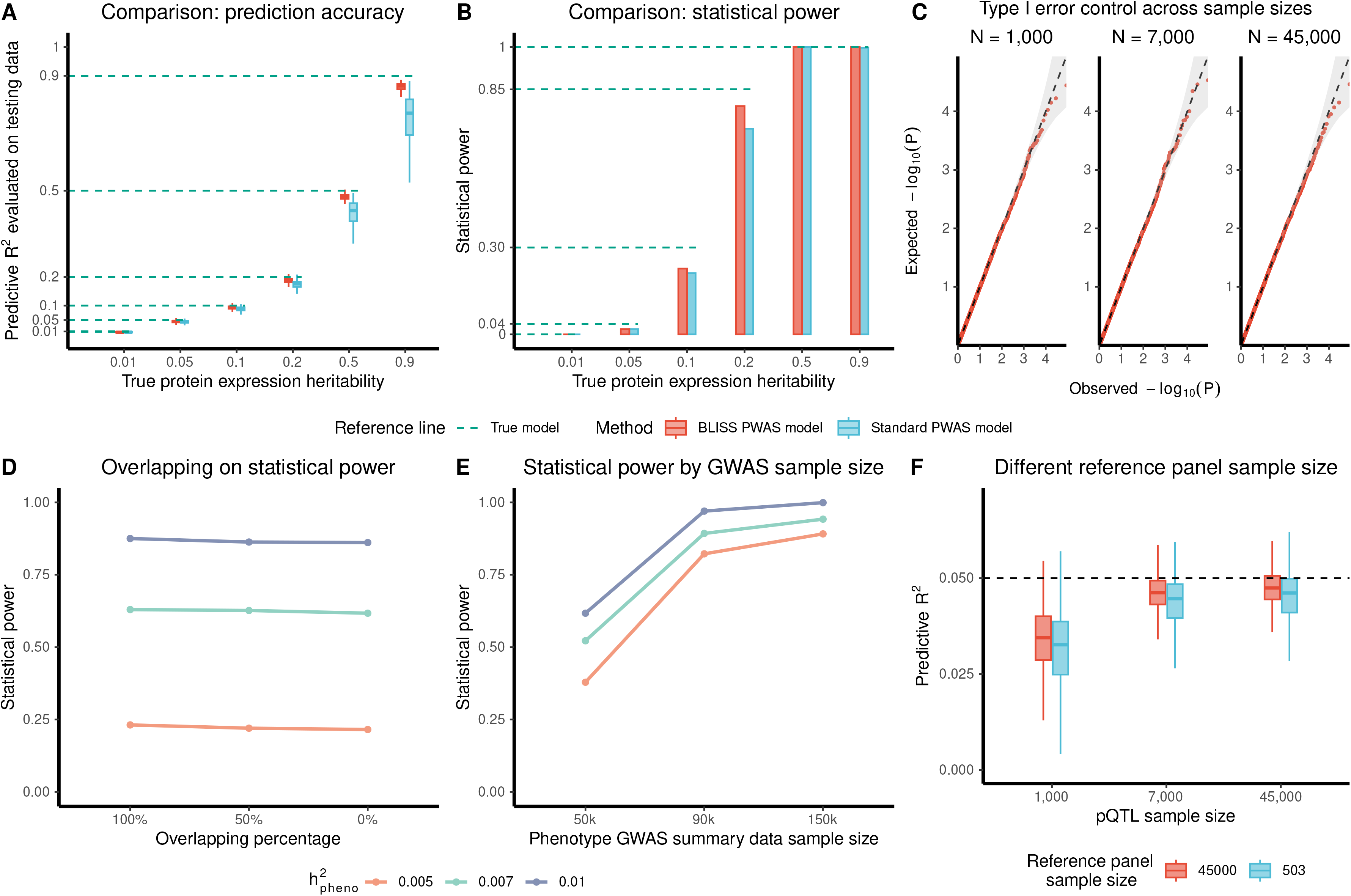
Simulation study evaluation of BLISS model performance. (A) Predictive accuracy comparison between BLISS and standard PWAS across varying protein expression heritability levels. Box plots show median, quartiles, and whiskers extending to 1.5 × IQR (interquartile range). (B) Statistical power comparison for association studies. Dashed lines indicate theoretical maximum power of the true model. (C) Quantile-quantile plots of p-values under null hypothesis across sample sizes (*n* = 1,000, 7,000, or 45,000). Gray bands represent 95% confidence intervals. (D) Statistical power versus data overlap percentage (proportion of BLISS training data also present in phenotype GWAS). (E) Statistical power as a function of phenotype GWAS sample size, stratified by phenotype heritability 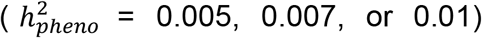. (F) Predictive *R*^2^ comparison for models using different reference panel sample sizes. Dashed line indicates preset protein expression heritability of 0.05. Box plots show median, quartiles, and whiskers extending to 1.5 × IQR.

Additionally, under the null hypothesis where there was no association between protein expression and phenotype, BLISS models yielded well-controlled empirical type I error rates across the training pQTL sample size (**Fig. 3C**), protein heritability levels (**Fig. S15**), and different LD reference panels (**Fig. S16**). These results indicate that the uncertainty inherent in BLISS models did not compromise type I error control under standard causal mediation assumptions^37^ (Supplementary Note). However, we found that BLISS models had reduced statistical power compared to true protein models where effect sizes are known and no imputation is needed. As expected, this power gap narrowed with increasing pQTL sample sizes, as larger samples resulted in more stable BLISS model weights (**Figs. 3B** and **S14**). These findings highlight that although model imputation uncertainty can reduce association power, BLISS preserves the validity of statistical tests under basic assumptions, a conclusion consistent with previous studies^38-40^.

We further investigated the impact of sample overlap between the datasets used for BLISS model training and those used for generating GWAS summary statistics. Specifically, we tested scenarios with 0%, 50%, and 100% sample overlap between the BLISS model training data and GWAS. Our analyses revealed that the degree of sample overlap had a negligible impact on the results, both in terms of statistical power and type I error rate (**Figs. 3D** and **S17**). These findings indicate the robustness of our approach regardless of the sample overlap between imputation model building and GWAS datasets. Additionally, we conducted analyses to assess the impact of both GWAS sample size and LD reference panel sample size. We found that increasing the GWAS sample size substantially improved power. Specifically, when phenotypic heritability equals 0.01, increasing the GWAS sample size from 50,000 to 90,000 and 150,000 led to power increases from 0.62 to 0.97 and 1, respectively (**Fig. 3E**). In contrast, increasing the LD reference panel size had only a minimal effect on performance. For example, expanding the LD reference panel from 503 to 45,000 resulted in only slight improvements in both predictive accuracy (**Fig. 3F**) and association test power (**Fig. S18**).

### Multi-ancestry protein-phenotype associations identified by BLISS

We applied BLISS models to GWAS summary statistics from the multi-ancestry MVP database^30^, which includes large cohorts of both European (*n* = 449,042) and African (*n* = 121,177) ancestry participants. The European cohort comprised 1,470 binary and 199 quantitative traits, and the African cohort included 1,187 binary and 181 quantitative traits. In this section, we present findings based on the Olink models, highlighting the similarities and differences in protein-phenotype associations across ancestries. To satisfy the relevance assumption, our analysis focused on proteins with *cis*-heritability greater than 0.01. A detailed comparison of Olink and SomaScan results across a broad spectrum of phenotypes will follow in a later section to provide further insights into platform-shared and specific effects.

In the European cohort, BLISS identified 32,072 significant protein-phenotype associations across 1,532 phenotypes when controlling the false discovery rate (FDR) at a 5% level across all proteins and phenotypes (*P* < 6.9 × 10^−4^, **Fig. 4A**). Comparing BLISS results to TWAS^41^, we found that 64.3% (9,601 out of 14,929) of these BLISS associations were not identified by PrediXcan^42^. For example, BLISS revealed a significant positive association between genetically predicted levels of IRE1α (encoded by *ERN1*) expression and type 2 diabetes (*β* = 0.11, *P* = 1.1 × 10^−5^), an association missed by TWAS after multiple testing adjustment (*P* = 2.9 × 10^−3^). IRE1α serves as a key sensor in the unfolded protein response pathway^43^, whose dysregulation is recognized as key contributors to type 2 diabetes pathogenesis^44^. In addition, ECM1 demonstrates significant osteological relevance through its multifaceted roles in skeletal homeostasis and exhibits pronounced functionality in osteogenesis and matrix mineralization. BLISS identified significant associations between ECM1 and several bone-related phenotypes, for example, localized osteoarthritis (*β* = 0.02, *P* = 3.7 × 10^−6^), substantiating its functional involvement in skeletal pathophysiology. These associations were not detected by TWAS (*P* > 0.5), highlighting the importance of post-transcriptional regulation and the ability of proteomic data to uncover associations missed by transcript-based approaches.

**Fig. 4.**
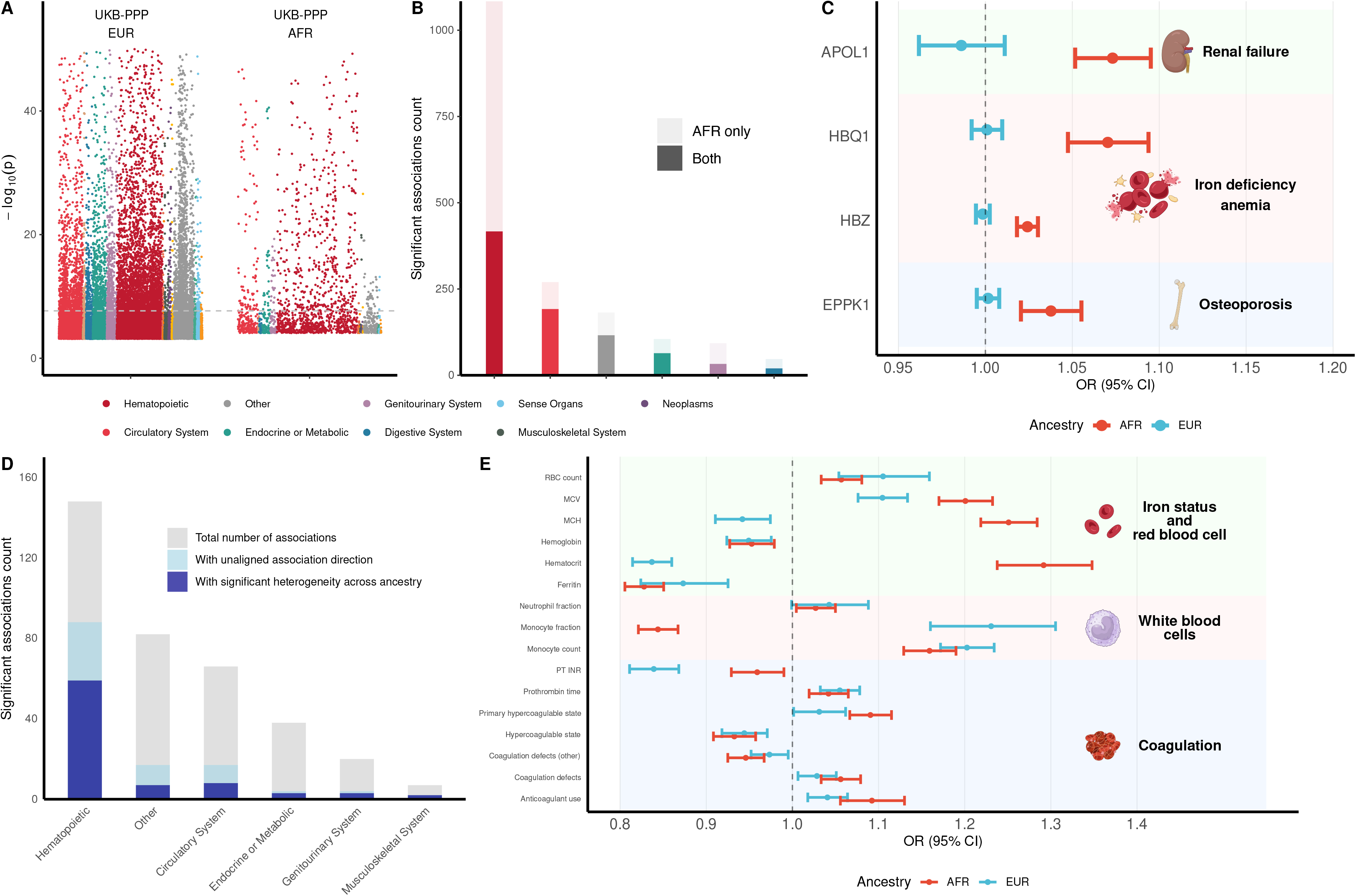
BLISS findings across ancestries. (A) Manhattan plots showing significant protein-phenotype associations in UKB-PPP European (EUR) and African (AFR) models. Each dot represents a significant association (controlling the false discovery rate at a 5% level), with colors indicating phenotype categories. The horizontal dotted line denotes the Bonferroni-corrected significance threshold. (B) Comparison of association counts across phenotype categories between EUR and AFR ancestries, stratified by AFR-ancestry specificity (AFR-only vs. both ancestries). (C) AFR-specific associations for three proteins with established biological relevance: APOL1 in renal failure, HBQ1 and HBZ in iron deficiency anemia, and EPPK1 in osteoporosis. We show odds ratios (OR) with 95% confidence interval (CI). (D) Distribution of proteins exhibiting significant heterogeneity between ancestries (Cochran’s Q test) across phenotype categories. Gray bars show total associations; blue bars indicate associations with significant heterogeneity. (E) OR with 95% CI for protein ABO’s associations with hematopoietic traits, comparing EUR (blue) and AFR (red) models.

In the African cohort, we identified 2,063 significant protein-trait associations across 444 phenotypes (**Fig. 4A**). Similar to European analysis, 961 (67%) of these BLISS associations were not identified by an African TWAS model^45^. Notably, when comparing BLISS findings across the two ancestry cohorts, 1,187 of the 2,063 significant associations (58%) were unique to the African cohort (**Fig. 4B**). While a small subset of these (109 associations involving 22 proteins) resulted from the absence of corresponding European-specific BLISS models, the majority (1,078 associations) arose because the protein-phenotype pairs did not reach significance in the European-specific analysis. These African-specific significant findings provide compelling biological insights (**Fig. 4C**). For example, a higher genetically predicted level of apolipoprotein L1 (APOL1) was strongly associated with an increased risk of renal failure (*β* = 0.07, *P* = 8.8 × 10^−12^), an association not detected in European analysis (*P* > 0.1). This finding provides molecular evidence linking APOL1 protein to the known higher prevalence of APOL1-driven kidney disease in African populations^46^. Another example is ancestry-specific associations in anemia-related pathways. In African analysis, we identified two key proteins involved in erythropoiesis that showed significant associations with iron deficiency anemia: Hemoglobin subunit theta-1 (HBQ1, *P* = 7.2 × 10^−10^) and Hemoglobin subunit zeta (HBZ, *P* = 1.9 × 10^−15^). These associations were exclusively observed in African cohort, with no significant associations observed in European analysis (*P* > 0.2). HBQ1 and HBZ, members of the alpha-globin gene cluster, are crucial for embryonic hemoglobin synthesis^47^. Their specific associations with iron deficiency anemia in African individuals may indicate a distinct regulatory mechanism or interaction with environmental factors more prevalent in this population, such as sickle cell disease or alpha-thalassemia^48,49^, potentially exacerbating hemoglobin deficits under iron-limited conditions. A third example involves Epiplakin (EPPK1) and osteoporosis, with significant associations observed in African analysis (*P* = 1.5 × 10^−5^) but not in European analysis (*P* > 0.2). While EPPK1 is primarily known for maintaining keratin network integrity and regulating keratinocyte behavior^50^, this novel association with osteoporosis suggests a potential link between keratin biology and bone health, possibly reflecting shared systemic processes influencing both nail keratin and bone protein integrity^51,52^.

To systematically compare effects across populations, we conducted heterogeneity analyses comparing protein-phenotype association coefficients between African and European cohorts for 1,367 shared phenotypes. Using Cochran’s Q test with Bonferroni correction, we identified 310 proteins exhibiting significant effect size heterogeneity (*P* < 5.7 × 10^−5^, **Table S2**). These heterogeneous proteins showed distinctive patterns in both lead variant alignment (defined as the ratio of shared lead variants to total lead variants across populations) and local LD structure (measured by average Frobenius norm differences between LD matrices) (**Fig. S19**). An example is FCGR2A’s association with leukocyte count: a lower genetically predicted level of FCGR2A was associated with higher leukocyte count (*β* = −0.16, *P* = 5.3 × 10^−114^), whereas the association was much weaker but significant in the opposite direction in the European cohort (*β* = 0.009, *P* = 3.5 × 10^−5^). This directional and strength difference suggests that the regulatory role of FCGR2A in leukocyte homeostasis^53^ may be ancestry-specific, potentially modulated by ancestry-specific immune exposures. Notably, this pattern of ancestry-opposite effects extends beyond individual pairs and is particularly enriched within specific phenotype categories. Specifically, among the protein-phenotype associations that reached statistical significance in both ancestries, 85 exhibited opposite directional effects, with a significant majority (59 of 85, 67%, *P* < 1 × 10^−20^ by a hypergeometric enrichment test) specifically linked to hematopoietic phenotypes (**Fig. 4D**). For example, associations between ABO and several hematopoietic phenotypes (hematocrit, monocyte fraction, and mean corpuscular hemoglobin) had opposite directions between European and African populations (**Fig. 4E**). This enrichment of hematopoietic-associated proteins may reflect the critical role of blood cell formation and immune function as targets of evolutionary selection, as populations encountered different pathogenic environments during human migration^54,55^.

Despite the 310 proteins showing evidence of heterogeneity, the effects for the majority protein were consistent across the two populations. After removing these 310 proteins with significant heterogeneity, we evaluated concordance across shared phenotypes. A weak positive correlation of *Z*-scores between African and European cohorts was observed across all protein-phenotype pairs, regardless of statistical significance (*ρ* = 0.11). This weak correlation is likely caused by the large number of null associations. When restricting the analysis to associations that were significant (FDR < 0.05) in at least one population, concordance strengthened considerably: 66% had consistent effect directions and the *Z*-score correlation increased (*ρ* = 0.44). Among the 879 protein-trait associations significant in both populations, the agreement was strongest, with 92% showing concordant effect directions and *Z*-score correlation being 0.71 (**Fig. S20**). We found that certain associations were consistently identified across both ancestries, even when the African models had substantially lower prediction accuracy than the European models. For example, both European and African models detected significant associations between lipoprotein(a) (LPA) and ischemic heart disease (*P* = 3.5 × 10^−222^ for European and *P* = 1.4 × 10^−5^ for African), despite a substantial difference in model performance (*R*^2^ = 0.37 for European and 0.03 for African). There is a well-established causal relationship between LPA and coronary artery disease, one of the strongest known genetic risk factors^56,57^. In addition, a systematic evaluation of discovery gains from multi-ancestry meta-analysis, including the benefits of incorporating super learner-enhanced African models, is provided in Supplementary Note (**Table S3**). Overall, our application of BLISS models to MVP data resources underscores the value of multi-ancestry imputation analysis in elucidating both ancestry-specific and shared protein-phenotype associations.

### Complementary drug target insights from BLISS and existing approaches

To evaluate the novel contribution of BLISS models, we systemically compared their results with those from TWAS, MR, and Bayesian colocalization analyses. Our analysis focused on a shared set of 594 genes/proteins that were analyzable by all four methods, using European ancestry data due to the larger sample sizes in both pQTL and GWAS summary statistics. At a 5% FDR level, BLISS identified 14,879 significant associations (*P* < 7.5 × 10^−4^), which was higher than that of TWAS (11,449 associations, *P* < 5.8 × 10^−4^) and MR (3,820 associations, *P* < 1.9 × 10^−4^). In addition, Bayesian colocalization identified 1,778 associations with high posterior probability (PPH4 > 0.8) of shared causal variants between protein and phenotype. While there was notable overlap: 5,315 (36%) of the BLISS associations were also identified by TWAS, 3,182 (21%) by MR, and 1,579 (11%) by colocalization, each method also contributed a distinct set of associations not captured by the others (**Figs. 5A-5B**).

**Fig. 5.**
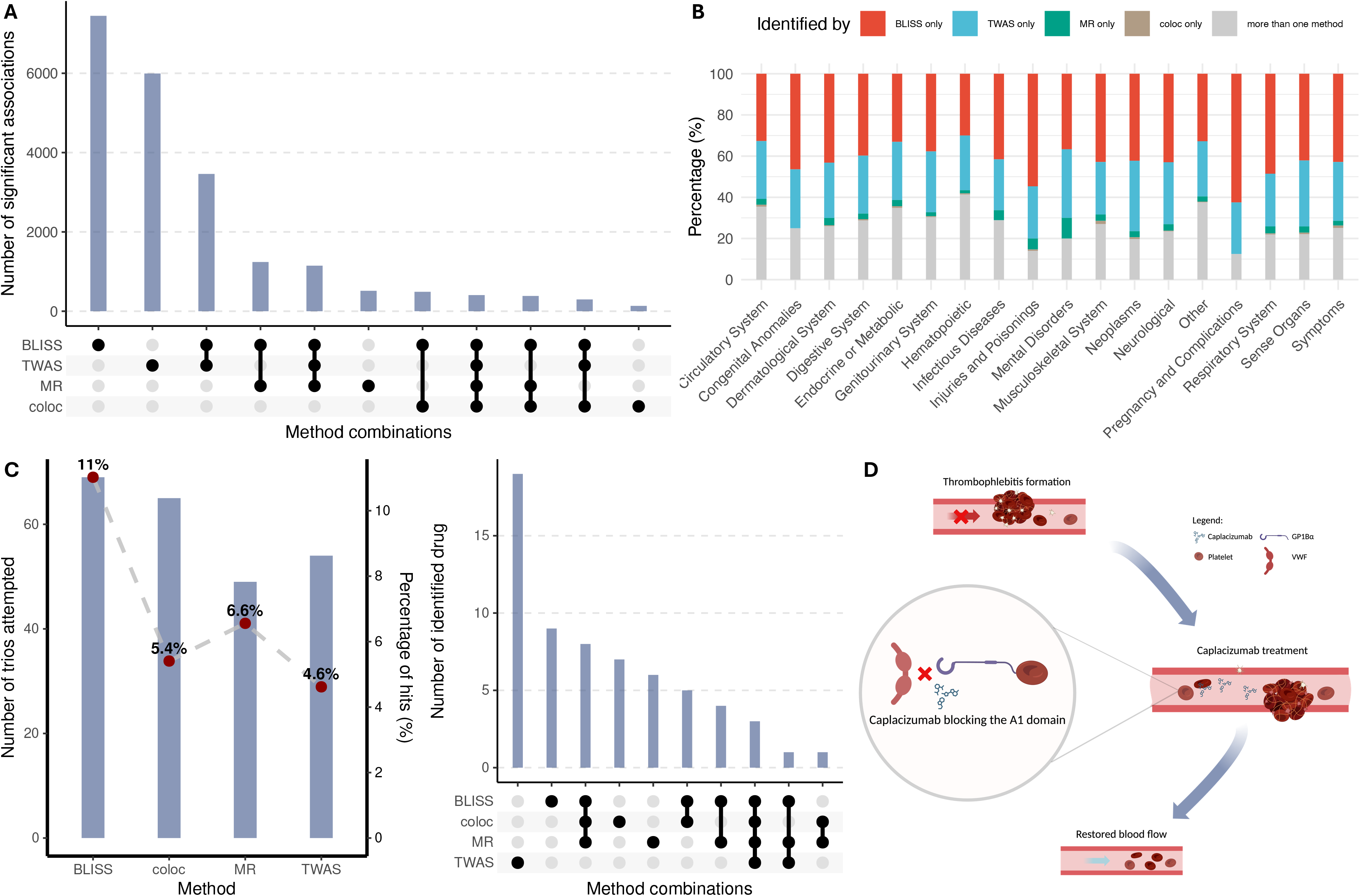
Comparison of BLISS with existing methods. (A) UpSet plot showing overlap of significant associations identified by different methods (TWAS, MR, and coloc). Bar heights indicate the number of associations found by each method combination, with filled circles below indicating which methods contribute to each intersection. (B) Proportion of associations identified exclusively by each method across phenotype categories. Colors indicate the identifying method: BLISS only (red), TWAS only (blue), MR only (green), coloc only (brown), and multiple methods (gray). (C) Performance comparison of all four methods for drug target identification. Left panel displays the percentage of successfully identified trios (red dots) with corresponding absolute numbers. UpSet plot on the right panel showing the overlap of trios identified by different methods. (D) Mechanistic illustration of the von Willebrand factor (VWF)-venous thrombosis-caplacizumab trio, exclusively identified by BLISS. The diagram shows the pathway from thrombophlebitis formation through caplacizumab’s mechanism of action (blocking the A1 domain) to restored blood flow.

More importantly, we found that BLISS models identified a larger number of therapeutically validated drug targets, with a higher level of enrichment (Methods). Specifically, when analyzing a recently curated dataset of 3,081 drug-target-phenotype trios^58^ (**Table S4**), BLISS achieved both the highest overlap rate and the largest absolute number of overlapping trios, 69 out of 626, compared to TWAS (54/1,169), MR (49/747), and colocalization (65/1,137) (**Fig. 5C**). Notably, each method could only analyze a distinct subset of these trios due to differences in coverage, resulting in the identification of partially non-overlapping drug targets. For example, 21 of the 69 (33.3%) trios identified by BLISS did not meet the PPH4 > 0.8 threshold for colocalization yet represent validated drug targets. One such example was the trio involving Von Willebrand factor (VWF), the target of Caplacizumab for treating venous thrombosis. The VWF-venous thrombosis association was uniquely identified by BLISS (*P* = 1.5 × 10^−3^), while it was missed by TWAS (VWF not included in models), MR (*P* > 0.1), and colocalization (PPH4 = 0) (**Fig. 5D**). Caplacizumab is a humanized nanobody that inhibits the interaction between VFW and platelets by targeting the VWF A1 domain^59^. This mechanism, which prevents pathological aggregation in thrombotic thrombocytopenic purpura, validates the clinical relevance of the positive association between plasma VWF with venous thrombosis incidence identified by BLISS^60^. Furthermore, BLISS demonstrated superior performance in African analysis, validating 9 out of 313 trios, compared to just 1 out of 1,580 for TWAS, 2 out of 451 for MR, and none out of 690 for colocalization. These results underscore the biological relevance of integrating complementary analytic methods and demonstrate the external validation of BLISS findings through independent drug target datasets.

### Cross-platform comparison in protein-phenotype associations

In this section, we compared protein-phenotype associations identified using two proteomic platforms: Olink (used in the UKB-PPP) and SomaScan (used in deCODE and ARIC). The Olink-based UKB-PPP dataset included pQTLs for 2,808 proteins in the European cohort and 2,797 proteins in the African cohort. The SomaScan-based datasets encompassed 4,710 proteins from deCODE, 4,435 from ARIC-EA, and 4,431 from ARIC-AA. There were 1,774 proteins common to both UKB-PPP and deCODE, 1,679 proteins shared across all three European ancestry datasets (UKB-PPP, deCODE, and ARIC-EA), and 1,684 proteins overlapping between the UKB-PPP AFR and ARIC-AA models (**Fig. S21**).

First, we compared our BLISS-trained ARIC models with the original ARIC models, which were trained on individual-level pQTL data, to assess which provided higher statistical power for downstream comparison. For ARIC-EA, BLISS-trained models identified 27,452 significant protein-phenotype associations (*P* < 6.8 × 10^−4^) at a 5% FDR, representing a 16.8% increase over the 23,505 associations (*P* < 5.6 × 10^−4^) detected by the original ARIC-EA models. Similar improvements were observed in ARIC-AA. BLISS-trained models identified 6,941 significant associations (*P* < 1.9 × 10^−4^), more than doubling the 3,248 associations (*P* < 8.6 × 10^−5^) discovered by the original ARIC-AA models. Despite the increased number of discoveries, the *Z*-scores for significant protein-phenotype pairs remained highly correlated between the BLISS-trained and original ARIC models, with correlation coefficients of 0.98 and 0.81 for the EA and AA cohorts, respectively (**Fig. S22**). Based on this enhanced power and overall consistency, we used the BLISS-trained ARIC models for all subsequent cross-platform comparisons.

We found that proteins unique to each platform contributed substantially to the complementary discoveries. In the European ancestry analysis, SomaScan-based deCODE models identified 25,200 significant protein-phenotype associations (*P* < 6.4 × 10^−4^) at a 5% FDR level. Of these, 18,016 associations were not detected by the Olink-based UKB-PPP models, including 12,640 associations involving 586 proteins exclusively measured by SomaScan (**Fig. 6A**). A notable example is the association between periostin (POSTN) and prostate cancer, which is a biologically plausible link given POSTN’s established role in promoting prostate cancer aggressiveness^61^. This association was detected by SomaScan-based models (*P* = 5.7 × 10^−4^) but was not captured by Olink due to lack of coverage. Conversely, Olink-based UKB-PPP models identified 22,098 significant associations that were not detected by the SomaScan-based models, including 18,492 associations involving 788 proteins uniquely measured by Olink. For example, the epidermal growth factor receptor (EGFR), an Olink-specific protein, showed a significant association with congenital foot deformities (*P* = 3.0 × 10^−4^), consistent with EGFR’s critical role in embryonic development, particularly in limb formation^62^.

**Fig. 6.**
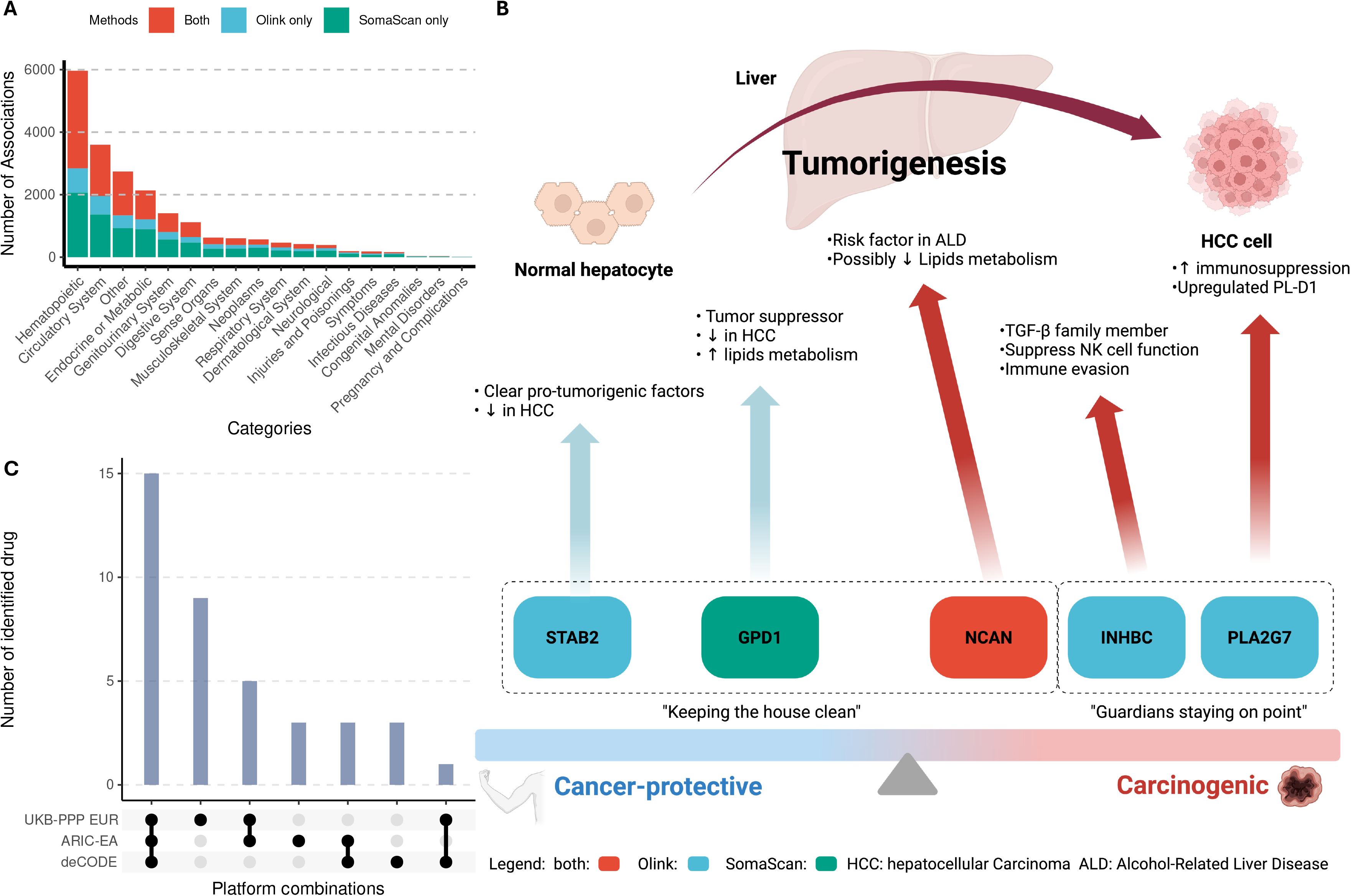
Comparison of BLISS findings between different proteome platforms. (A) Number of significant associations identified across phenotype categories, stratified by detection platform. Colors indicate method overlap: both platforms (red), Olink only (blue), and SomaScan only (green). (B) Schematic illustration demonstrating how proteins detected by the two platforms provide complementary biological insights into hepatocellular carcinoma etiology. (C) Overlap of protein-drug-phenotype trios identified between Olink-based (UKB-PPP) and SomaScan-based models (deCODE and ARIC-EA). Bar heights show the number of trios identified by each platform combination, with filled circles indicating contributing platforms.

To further assess cross-platform differences and similarities, we focused on proteins shared across all three European ancestry models (UKB-PPP, deCODE, and ARIC-EA). Within this common protein set, the UKB-PPP models identified 22,656 significant protein-phenotype associations (*P* < 6.8 × 10^−4^), while deCODE and ARIC-EA models identified 12,574 (*P* < 6.4 × 10^−4^) and 13,370 (*P* < 6.7 × 10^−4^) associations, respectively, all at a 5% FDR threshold. UKB-PPP models revealed 12,762 significant associations not detected by either deCODE or ARIC-EA. Conversely, deCODE models identified 3,952 associations absent in UKB-PPP and ARIC-EA. Despite these platform-specific discoveries, we observed strong concordance in association statistics across datasets. Among protein-phenotype pairs significant in at least one model, the cross-platform *Z*-score correlations were robust: *ρ* = 0.83 between UKB-PPP and deCODE, *ρ* = 0.81 between UKB-PPP and ARIC-EA, and *ρ* = 0.89 between ARIC-EA and deCODE (**Fig. S23**). To rigorously evaluate the consistency of effect size, we conducted Cochran’s Q tests with Bonferroni correction. Between the SomaScan-based ARIC-EA and deCODE models, both using the same platform, only 113 out of 4,398 proteins (3%) exhibited significant heterogeneity. Cross-platform comparisons showed slightly higher but still small heterogeneity: 113 out of 1,685 proteins (7%) between ARIC-EA and UKB-PPP, and 132 out of 1,774 proteins (7%) between deCODE and UKB-PPP. Excluding these heterogeneous proteins further improved *Z*-score concordance: the UKB-PPP vs. deCODE correlation increased from 0.83 to 0.87, and ARIC-EA vs. deCODE from 0.89 to 0.90. Furthermore, across all 31,327 significant associations (*P* < 6.8 × 10^−4^; **Table S5**) identified by any of the three European ancestry models, 36% (11,335 associations) were significant in at least two models, and 19% (5,940 associations) were shared across all three. These 5,940 overlapping associations exhibited high *Z*-score consistency across datasets, with correlation coefficients ranging from 0.887 to 0.946. Notably, over 95% of these shared associations demonstrated consistent directions of effect (**Fig. S24**). We conducted a similar comparison for the African ancestry-based models (UKB-PPP AFR and ARIC-AA) and observed comparable patterns (Supplementary Note and **Figs. S25-S26**).

The complementary discoveries from the Olink and SomaScan platforms together provide a more comprehensive view of the biological mechanisms underlying disease, as each platform may illuminate different component of disease pathways. One example is about complement and coagulation cascades. Both platforms concordantly identified associations for factor XI (F11), factor II (F2/prothrombin), and factor VII (F7), establishing a shared foundation of pathway dysregulation.

However, each platform also captured complementary regulatory elements. Olink uniquely identified associations for factor X (F10; *P* = 8.5 × 10^−9^) and protein C (PROC, *P* = 1.5 × 10^−6^), capturing key components of the common coagulation pathway and its regulation. Notably, the PROC system plays a vital role as a natural anticoagulant pathway, regulating blood coagulation and preventing excessive clot formation^63^. On the other hand, SomaScan exclusively identified factor V (F5, *P* = 4.9 × 10^−32^), complement factor H (CFH, *P* = 6.1 × 10^−4^), SERPINA5 (protein C inhibitor, *P* = 4.3 × 10^−4^), and SERPINF2 (α2-antiplasmin, *P* = 1.1 × 10^−4^), revealing both the complement regulatory arm and critical serpin inhibitors of the cascade. These SomaScan-specific discoveries have important biological implications. SERPINF2 serves as the primary plasmin inhibitor in fibrinolysis, while SERPINA5 has dual coagulant and anticoagulant functions through its interactions with multiple proteases^64^. In addition, the identification of CFH reveals a critical regulatory hub that the Olink platform missed. CFH mutations are associated with diverse pathologies including age-related macular degeneration, atypical hemolytic uremic syndrome, and C3 glomerulopathy^65^. Furthermore, the joint detection of F11 and CFH aligns with recent evidence of cross-talk between the complement and coagulation systems^66^, as activated F11 can cleave and neutralize CFH, enhancing complement activation. This interaction is evident only when both F11 and CFH are identified.

Complementary insights also emerge in the interleukin signaling pathway implicated in asthma pathogenesis. While both platforms detected AGER and IL1RL1, suggesting robust associations with innate immunity and type 2 inflammation, each platform revealed distinct immunological components. Olink exclusively identified IL7R (*P* = 5.8 × 10^−6^), IL1R1 (*P* = 7.1 × 10^−6^), and ITGAM (*P* = 4.5 × 10^−4^), capturing adaptive immune and myeloid cell components. Notably, IL7R encodes the alpha chain of the IL-7 receptor, which plays crucial roles in T cell homeostasis and has emerged as an important regulator of innate lymphoid cells that contribute to type 2 inflammation in asthma through TSLP signaling^67^, a pathway component that would remain undetected using SomaScan platform alone. Conversely, SomaScan uniquely detected CBL (*P* = 2.0 × 10^−15^) and IL2RB (*P* = 3.8 × 10^−4^), revealing signal transduction and T cell regulatory elements.

The value of this multi-platform approach is further demonstrated in hepatocellular carcinoma (HCC), where we identify a dual mechanistic framework disease progression (**Fig. 6B**). The first component involves disruption of cellular maintenance and metabolic homeostasis, characterized by three key proteins: stabilin-2 (STAB2), glycerol-3-phosphate dehydrogenase 1 (GPD1), and neurocan (NCAN). STAB2 is a scavenger receptor responsible for clearing waste products from the bloodstream, including modified lipoproteins, apoptotic cells, and other ligands. Through these clearance functions, STAB2 plays a vital role in maintaining liver homeostasis^68^. GPD1 regulates lipid metabolism by converting dihydroxyacetone phosphate to glycerol-3-phosphate, a key step in triglyceride synthesis, and is increasingly recognized as a potential contributor to liver disease^69^. NCAN, an extracellular matrix proteoglycan, is associated with increased risk for HCC in patients with alcoholic liver disease^70^, possibly by disrupting lipid metabolism homeostasis. In our analysis, NCAN was consistently identified by all models (*P* < 4.1 × 10^−9^). In contrast, STAB2 was uniquely detected by the Olink-based UKB-PPP model (*β* = 0.009, *P* = 2.0 × 10^−4^), while GPD1 was exclusively identified by the SomaScan-based ARIC-EA model (*β* = 0.04, *P* = 2.1 × 10^−4^). The second component involves failures in immune surveillance and tumor suppression, highlighted by two key proteins: inhibin subunit beta C (INHBC) and phospholipase A2 group VII (PLA2G7). INHBC, a member of the TGF-β superfamily, contributes to immune evasion by suppressing natural killer (NK) cell function, facilitating cancer cell escape from immune detection^71^. Complementing this mechanism, PLA2G7 acts as a potent immunosuppressive factor within the HCC tumor microenvironment. High PLA2G7 expression in tumor-associated macrophages is associated with poor prognosis and resistance to anti–PD-1 immunotherapy, primarily by impairing CD8^+^ T cell activation^72^. Mechanistically, PLA2G7 promotes immunosuppression via the STAT1/PD-L1 axis, directly upregulating PD-L1 expression and fostering an environment conducive to tumor progression^73^. In our analysis, both INHBC and PLA2G7 were uniquely identified by Olink-based models (*P* < 4.3 × 10^−4^). In summary, Olink captures critical cellular maintenance disruption (STAB2) and immune evasion mechanisms (INHBC and PLA2G7), while SomaScan reveals essential metabolic dysregulation (GPD1). Both platforms converge to identify NCAN’s multifaceted role in genetic susceptibility and metabolic disruption. These findings underscore the value of combining data from multiple proteomic platforms to uncover the complex interplay of metabolic reprogramming, immune evasion, and genetic predisposition that underlies HCC development. This pattern of complementary insights extends throughout our analysis. We provided a summary table of 100 distinct biological pathways across 108 phenotypes in which at least 25% of constituent proteins were captured by BLISS. Most of these pathways contain both platform-specific findings and proteins measured by both platforms (**Table S6**).

In addition, we evaluated the performance of each platform in prioritizing known therapeutically relevant drug targets^58^. For European ancestry models, both platforms demonstrated comparable overlap rates and numbers of overlapping targets. Specifically, the Olink-based UKB-PPP models identified 69 out of 626 trios (11.0%), while the SomaScan-based deCODE and ARIC-EA models identified 49 out of 391 (12.5%) and 61 out of 435 (14.0%) trios, respectively (**Fig. S27**). Fifteen trios involving 9 unique proteins were consistently identified across all three sets of models (**Fig. 6C**). Each platform also uniquely identified additional validated targets, 9 by UKB-PPP Olink and 3 by SomaScan (deCODE or ARIC-EA). Similar patterns were observed in the African cohort. UKB-PPP AFR models identified 9 out of 313 (2.9%) trios, while ARIC-AA models identified 13 out of 365 (3.6%) trios at a 5% FDR level.

### Atlas of multi-ancestry protein-phenotype associations

In addition to analyzing MVP GWAS datasets, we conducted a systematic multi-ancestry analysis and expanded the catalog of protein-phenotype associations using four additional genetic data resources from GBMI^31^, FinnGen^32^, IEU OpenGWAS^33^, and BBJ^34^ (**Table S7**). For GBMI, focusing on seven diseases with multi-ancestry GWAS summary statistics, we identified 171 significant protein-disease associations (*P* < 7.3 × 10^−4^) at a 5% FDR level, 56 of which achieved significance in more than one ancestry. FinnGen data analysis yielded 3,812 significant associations (*P* < 7.4 × 10^−4^) across 299 diseases. For IEU OpenGWAS, we identified 5,435 significant protein-phenotype associations (*P* < 7.4 × 10^−4^) spanning 310 distinct phenotypes. For BBJ dataset, we identified 1,339 significant associations (*P* < 3.1 × 10^−4^) across 215 phenotypes.

We developed an interactive web tool (https://www.gcbhub.org/) to enable quick and in-depth exploration of our protein-phenotype association results. This resource integrates major biobank-scale GWAS datasets with the latest pQTL data, covering over 2,500 phenotypes and their corresponding protein signals. By providing these resources, we aim to accelerate the discovery of protein-mediated disease mechanisms and support the prioritization of novel therapeutic targets across diverse populations.

## Discussion

In this study, we developed a statistical framework for training protein imputation models using the latest multi-ancestry pQTL data from both Olink and SomaScan platforms. We applied these models to recent multi-ancestry GWAS summary statistics to address key questions in the genetic mapping of protein-phenotype associations (**Fig. 1C**). Our analysis revealed novel biological insights into the proteomic architecture of complex traits and diseases, highlighted ancestry-specific differences in protein-phenotype associations, and underscored both the converging and distinct contributions of the Olink and SomaScan platforms. Additionally, we demonstrated how our new models provide complementary strengths in prioritizing potential drug targets. All developed models and the full set of our analysis results have been made freely available at https://www.gcbhub.org/.

We demonstrated the complementary associations and biological insights revealed by the Olink and SomaScan platforms. For proteins measured by both platforms, the observed association differences align with the recent report to compare the proteomics data from Olink UKB-PPP and SomaScan deCODE^36^, which may be related to the fact that SomaScan and Olink platforms detect different proteoforms for some proteins. These proteoforms can participate in distinct biological processes and consequently show different associations with phenotypes. Additionally, some platform-specific associations may reflect epitope effects rather than actual protein level differences, particularly when associated with coding variants without corresponding expression changes. In addition, while our super learner integration substantially improved within-platform model performance, applying the same framework to combine pQTL data from distinct platforms did not confer similar benefits, instead resulting in reduced predictive accuracy compared to using the single-platform (Supplementary Note and **Fig. S28**). This observation may be consistent with the intrinsic heterogeneity of the data sources. We recommend interpreting results for heterogeneous proteins with caution, taking into account platform-specific context.

In training BLISS models, we found that model performance is largely robust to the choice of LD reference panel, although using large, cohort-matched LD panels can slightly increase predictive *R*^2^ (**Fig. S4**). All LD reference panels used in our study have been made publicly available, and we recommend using the largest ancestry-matched panel when cohort-specific LD data are not accessible. Due to limited sample sizes, we grouped all Asian ancestry samples from UKB-PPP together, which may not fully capture the genetic heterogeneity across Asian subpopulations. As proteomic data resources expand, particularly with the anticipated release of data from 500,000 UKB-PPP participants in the next few years, we plan to continuously improve our BLISS models. Future updates will incorporate larger pQTL sample sizes and more fine-grained, population-specific models, such as those for Eastern and Southern Asian populations, to ensure our online resources remain up to date.

We conclude our manuscript by discussing several limitations and outlining future directions. First, our current super learner framework, designed to improve model performance in non-European ancestries by leveraging large-scale European data, requires access to small-scale individual-level pQTL data from the target non-European ancestry. This represents a deviation from the purely summary-statistic-based nature of our foundational BLISS methodology. Although this approach yields significant improvements in prediction accuracy (**Figs. 2B-2C** and **S10**), the development of sophisticated super learning techniques capable of achieving comparable enhancements using only summary statistics constitutes a critical next step. Such an advance, particularly one adept at handling the often small non-European pQTL datasets, would further democratize the application of powerful cross-ancestry predictive modeling.

Second, although the imputation framework underlying PWAS effectively identifies protein-phenotype associations and helps prioritize proteins for downstream validation, careful interpretation is crucial, particularly concerning causality. The inherent complexities of genomic architecture, such as the common co-regulation of neighboring genes within intricate loci and the potential for LD to confound signals, means that a PWAS association for a specific protein does not necessarily establish its causality^74^. Furthermore, the current incomplete coverage of the proteome by existing platforms implies that the true causal protein might not always be measured. Consequently, PWAS methods including BLISS is most effectively utilized as a powerful hypothesis-generating tool that prioritizes candidate proteins for further investigation. We have conducted a comprehensive comparison with other analytical approaches, including TWAS, MR, and statistical colocalization, highlighting their complementary insights into drug target validation, as well as the unique contributions of each method. Integrating PWAS findings with evidence from these approaches can help generate a more comprehensive list of protein candidates for subsequent experimental validation^75^. A more detailed discussion on these interpretative aspects and suggestive practices is provided in Supplementary Note.

## METHODS

Methods are available in the **Methods** section.

*Note: One supplementary pdf file and one supplementary table file are available*.

## ACKNOWLEDGEMENTS

Research reported in this work was supported by the National Cancer Institute under Award Numbers R01CA263494, U01CA293883, and P30CA016672; National Institute of Mental Health under Award Number R01MH136055; and National Institute on Aging under Award Numbers RF1AG082938 and R01AG085581. The content is solely the responsibility of the authors and does not necessarily represent the official views of the National Institutes of Health. We thank the individuals represented in the genetic and proteomic studies for their participation and the research teams for their work in collecting, processing, and disseminating these datasets for analysis. We gratefully acknowledge all the studies and databases that made pQTL and GWAS summary data available. This research has been conducted using the UK Biobank resource (application number 76139), subject to a data transfer agreement. We would like to thank the research computing groups at the University of Texas MD Anderson Cancer Center, Purdue University, and the Wharton School of the University of Pennsylvania for providing computational resources and support that have contributed to these research results.

## AUTHOR CONTRIBUTIONS

B.Z. and C.W. designed the study. C.W., Z.Z., X.Y., and B.Z. developed the models, analyzed the data, and interpreted the results. Z.Z. and C.W. performed the simulation study. Z.Z. developed the online data resources. B.Z. and C.W. wrote the manuscript with feedback from all authors.

## METHODS

### pQTL data resources

We used pQTL summary statistics from the UK Biobank (UKB), deCODE, and ARIC studies. Detailed descriptions of the deCODE and ARIC pQTL summary statistics can be found in Eldjarn, et al. ^36^ and Zhang, et al. ^8^, respectively. Briefly, in the deCODE cohort, the average participant age was 55 years and 57% were women. The deCODE pQTL study included 35,892 Icelanders who had both genetic and proteomic data available. The plasma proteome data were measured using the SomaScan version 4 assay (SomaLogic) and SMP normalization was applied for improved data quality^36^. Imputed variants with minor allele frequency (MAF) > 1% and imputation information > 0.9 were used in the deCODE pQTL study. The rank-inverse normal transformed levels were used in the linear mixed model implemented in BOLT-LMM^76^, while adjusting for age and sex effects. The ARIC pQTL study, also using the SomaScan version 4 assay, comprised data from 7,213 European American participants and 1,871 African American participants. Genotypic data were collected by the Affmetrix 6.0 DNA microarray and subsequently imputed to the TOPMed reference panel^77,78^ (Freeze 5b). For pQTL analysis, the log-transformed relative abundance values were residualized to adjust for probabilistic estimation of expression residual factors^79^, sex, age, study site, and ten genetic principal components (PCs). The residuals were rank-inverse normalized and pQTL analysis was performed using QTLtools^80^. For both deCODE and ARIC pQTL data, we followed the pipeline outlined in Zhang, et al. ^8^, focusing on 4,483 unique proteins or protein complexes encoded by 4,435 autosomal genes passed quality controls (QCs), including excluding aptamers that mapped to multiple gene targets and without any SNPs in the *cis*-region. To prevent potential interference from the Major Histocompatibility Complex region, we excluded all single nucleotide polymorphisms (SNPs) located within chr6:28,510,120-33,480,577 (GRCh38), and removed genes with substantial overlap in this region from our sequential model building.

We derived the UKB pQTL summary statistics from 49,341 white British participants in the UKB-PPP project, focusing on 2,815 proteins (**Table S1**). We used the imputed genotyping data^81^ and any imputed variants with a QC *r*^*2*^-value below 0.8 were excluded. Subsequent to this, we applied standard QC procedures^82^, which included i) omitting subjects with over 10% missing genotypes; ii) discarding variants with an MAF under 0.01; iii) removing variants with a missing genotype rate exceeding 10%; and iv) excluding variants that did not pass the Hardy-Weinberg equilibrium test at a threshold of 1×10^−7^ level. For every protein, outliers were excluded, which were defined as values exceeding five times the median absolute deviation from the median. We used linear mixed-effect models with fastGWA^83^, accounting for effects of covariates including age, sex, genotype array, visiting center ID, age squared, the interaction between age and sex, the interaction between age squared and sex, and the top 40 genetic PCs^81^. We considered *cis*-SNPs within each gene region and extended our consideration to a 1-megabase window on either side. Additionally, we used three smaller UKB proteomic datasets for the purpose of assessing prediction accuracy or generating non-European super learner models. These datasets included 2,923 white non-British subjects, 1,171 African ancestry subjects, and 914 Asian ancestry subjects.

To avoid potential biases, we randomly eliminated one from any pair of related individuals within each of the three small datasets and excluded any relations of the white British subjects.

### Biomarker expression Levels Imputation using Summary Statistics (BLISS)

To train the protein expression imputation model only using pQTL summary statistics and a linkage disequilibrium (LD) reference panel, we introduced a new method called BLISS. Consider the following linear regression model for protein expression:

where *X* is the protein expression levels, *G* = (*G*_1_, ⋯, *G*_*p*_)′ is the standardized genotype matrix of *p cis*-SNPs around the gene, *w* = (*w*_1_, ⋯, *w*_*p*_)′ is the effect size to be estimated, and *ε* is random noise with a mean of zero. The objective function *f*(*w*) for a penalized regression to estimate *w* is:

where *J*_*λ*_(*w*) is a penalty term such as Elastic-net, Lasso, or MCP, and *N* is the sample size of the pQTL study. We generated solutions for a candidate set of tuning parameters. Specifically, *G*^′^*X*/*N* represented the correlation between cis-SNPs and protein expression levels, which can be estimated by 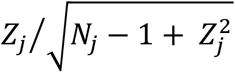, where *Z*_*j*_ and *N*_*j*_ are the *Z* score (estimated effect size divided by its standard error) and sample size for SNP *j*, respectively. The term *G*^′^*G*/*N* was the LD correlation matrix, which can be estimated using LD reference panel such as the 1000 Genomes Project^84^ or UKB of matched ancestry. We used the shrinkage estimator and shrunk the off-diagonal entries toward zero based on their genetic distance^29,85,86^. To ensure a unique solution upon optimization, a small *L*_2_ penalty term *θw*^′^*w* was added to the objective function. Then the objective function was optimized through a coordinate descent algorithm.

Notably, a key advantage in BLISS is its ability to generate independent pseudo-training, pseudo-tuning, and pseudo-ensemble learning datasets from summary-level pQTL data, avoiding the needs of matched individual-level pQTL data to select tuning parameters such as in SUMMIT^29^. We achieved this by sampling marginal association statistics of pQTL data for subsets of individuals conditional on the complete summary-level pQTL data. Specifically, we generated the pseudo-training data 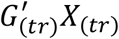 with sample size *n*_1_ by a conditional distribution: 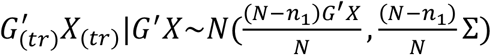. Here · | · denotes the conditional distribution and Σ is the observed covariance matrix, which can be estimated by using minor allele, estimated effect sizes, and standard errors provided by pQTL summary data^35^. We then calculated the estimated effect sizes and their standard errors for pseudo-training data and obtained the remaining independent data 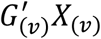 by 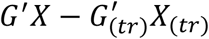, following the same approach used in meta-analysis tools such as METAL^87^. Next, we generated the pseudo-tuning data sample size *n*_2_ by a conditional distribution: 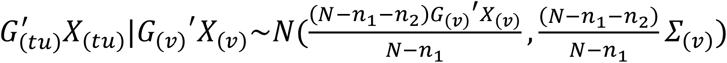.

Using the same methodology, we generated the pseudo-ensemble learning data 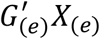 with sample size *n*_3_. Finally, the pseudo-testing data 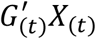 were obtained by subtracting all previously generated datasets from 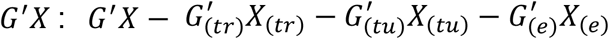. By default, we set *n*_1_ = 0.7 × *N, n*_2_ = 0.1 × *N*, and *n*_3_ = 0.1 × *N*.

We trained the prediction models for different set of SNPs selected by varying LD clumping thresholds (*R*^2^ = 0.01, 0.05, 0.1, 0.2, 0.5, and 0.9). Specifically, SNPs were clumped if they were *r*^*2*^ greater than clumping thresholds with the index variant to improve the signal-to-noise ratio (window size = 10 megabases, which performed whole-genome clumping). Under each clumping threshold, we calculated the summary-level predictive *R*^*2*^, which was the squared Pearson correlation coefficient between genetically predicted and directly measured protein expression levels, using the pseudo-tuning data^35^. We selected the tuning parameters that maximized the summary-level predictive *R*^*2*^. We repeated the above procedure five times and selected the parameters that maximized the average predictive *R*^*2*^ across these five replicates. Recognizing that different models may capture varying but complementary predictive genetic architecture, we combined models from different clumping thresholds using the pseudo-ensemble learning dataset. Specifically, we produced weighted sum of predictions from different models, requiring each model to contribute positively (i.e., having a corresponding coefficient greater than zero). Finally, we calculated the summary-level predictive *R*^*2*^ using the pseudo-testing data 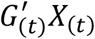. We repeated the above procedure five times and selected the ensemble weighting parameters that maximized the average predictive *R*^*2*^ across these five replicates. We then trained protein prediction models using all data and the selected tuning parameters and ensemble weights. These models were used in the subsequent analysis.

### Super learner integration for non-European ancestry

Recognizing that the BLISS models for Europeans were built on much larger sample sizes and could potentially improve the prediction accuracy of non-European models, we integrated the BLISS-based European models with non-European models using the pseudo-ensemble learning dataset. Specifically, for each non-European ancestry group (e.g., Asian and African), we added European BLISS-trained models as a contributing model in ensemble learning step in our BLISS framework. Notably, we required each contributing model positively influenced the ensemble prediction, thus this method allows us to capitalize on the strengths of models trained on large, European-centric datasets while incorporating ancestry-specific information to improve performance in diverse populations.

### BLISS model training and evaluation

We applied BLISS to train models using above-mentioned pQTL data resources, including summary-level pQTL data from deCODE, UKB-PPP (European, African, and Asian separately), and ARIC (European and African separately). We focused on models where the number of SNPs with non-zero weights was greater than 15, since having a sufficiently large number of these SNPs was crucial to ensure reliable asymptotic statistical properties during the second stage of association tests. Additionally, for non-European populations, we integrated the super learner framework to enhance model training for African and Asian ancestries by leveraging BLISS-trained European models. For genotype data, we excluded SNPs with MAF < 1%, those that were non-biallelic or had ambiguous strand mapping (i.e., no A/T or C/G SNPs), and those did not present in the LD reference panel. We summarized the models and related information in **Table S1**.

The performance of the imputation models was evaluated by predictive *R*^*2*^, the squared Pearson correlation coefficient between genetically predicted and directly measured protein expression levels, which was provided by both BLISS and super learner. To assess the reliability of the summary-level predictive *R*^*2*^ values in BLISS, we calculated the individual-level predictive *R*^*2*^ using an independent testing dataset comprising 2,923 white non-British individuals from the UKB study. Specifically, we used BLISS weights derived from our UKB pQTL analysis from white British individuals to predict protein expression levels in the 2,923 white non-British UKB individuals. The individual-level predictive *R*^*2*^ values were compared with those calculated from BLISS. The performance of these non-European super learner models was also evaluated. As a benchmark, we first trained baseline models for the same non-European cohorts (UKB-PPP Asian and African) using a standard Elastic-net pipeline applied directly to the individual-level data, and these baseline models were also evaluated using the procedures described below. Our primary evaluation used a nested 5-fold cross-validation procedure (https://github.com/hakyimlab/PredictDB_Pipeline_GTEx_v7/blob/master/README.md) within the respective non-European datasets. The inner loop of this nested cross-validation was used for model tuning and training the super learner weights, while the outer loop provided an unbiased estimate of out-of-sample predictive performance. To further corroborate the reliability of this nested cross-validation, we also conducted an independent 80/20 split validation for both Asian and African ancestries within UKB-PPP, where super learner were applied on 80% of the data and predictive *R*^*2*^ was assessed on the held-out 20%.

### Protein-phenotype association testing

When only summary-level GWAS data were available, standard PWAS applied a burden-type *Z* test to examine the association between the genetically predicted protein expression levels and phenotype^8,19^. Under the null hypothesis, the test statistics was given by 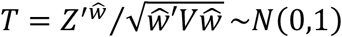, where *Z* is the vector of z-scores from GWAS, 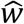 is the estimated weights from BLISS model, and *V* is the LD matrix of corresponding SNPs estimated from a population reference panel. It was worth mentioning that one assumption underlying this *Z* test was that the LD matrix estimated from the reference panel precisely matched that from the GWAS data. This alignment may be crucial as any discrepancy could potentially lead to an inflated type 1 error rate^37^. To reduce the potential negative effects of such LD covariance matrix discrepancy, in our association test, we estimated the effect size 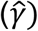 and its variance between the protein and phenotype by^37^

where *n*_*s*_ is the sample size of the GWAS data, 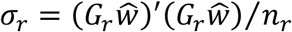, and 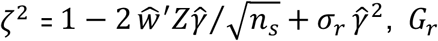, *G*_*r*_ is the standardized (mean 0 and variance 1) genotype matrix of the population reference panel, and *n*_*r*_ is its sample size. Notably, while uncertainty in the protein prediction model may affect statistical power, it does not impact the type I error rate under valid instrumental variable assumptions (Supplementary Note).

### Simulations

We used the UKB imputed genotype data^81^ to conduct simulations, focusing on evaluating the protein prediction accuracy, testing power, and type 1 error rate of different methods. Specifically, we selected three separate sets of unrelated White British individuals: the first 45,000 individuals as a training dataset for imputation model building, the following 10,000 individuals as an independent testing dataset for evaluating predictive *R*^*2*^, and the following 50,000 individuals for generating GWAS summary statistics. To ensure the generalizability of our results, we included all proteins in UKB-PPP to generate simulated data and used the real-data SNP list of each protein as our starting point. Additionally, to reduce computational burden, we restricted our simulations to HapMap SNPs.

Protein expression levels (*X*) and phenotype values (*Y*) were simulated by *X* = *Gw* + *ε*_*x*_ and *Y* = *γX* + *ε*_*y*_, respectively. Here *G* is the genotype matrix, *w* is the vector of effect sizes in the imputation model, the scalar *γ* is the association coefficient between protein expression and phenotype, and *ε*_*x*_ and *ε*_*y*_ are vectors of normally distributed noise to achieve pre-specified protein expression heritability and phenotypic heritability. We generated pQTL summary data by running a marginal association scan between protein expression and each SNP in the imputation training data. Additionally, GWAS summary statistics were generated by regressing the phenotype on each SNP in the GWAS data. We considered three different imputation training sample sizes, 1,000, 7,000, and 45,000, to investigate the impact of sample size on model prediction accuracy. We investigated the impact of uncertainty in protein prediction model accuracy on the performance of BLISS by comparing BLISS-trained models, which incorporate estimation error, with true protein prediction models that do not have any estimation error. Furthermore, we examined the impact of the choice of LD reference panel on the predictive accuracy of BLISS models. We compared the performance of BLISS using different LD reference panels, including the 1000 Genomes Project and the UKB datasets. Given that biobank data have been widely meta-analyzed in modern GWAS, we considered three settings where 0%, 50%, and 100% of individuals in the training dataset were also part of the GWAS dataset.

We considered two scenarios with different proportions of causal SNPs: 5% and 10% of SNPs within each pQTL region were assigned non-zero effects sampled from a standard normal distribution. We also examined different levels of protein expression heritability (0.01, 0.05, 0.1, 0.2, 0.5, and 0.9) and phenotypic heritability (0.001, 0.003, 0.005, 0.007, and 0.01). For each scenario, we generated training and testing datasets for each protein to evaluate the predictive *R*^*2*^. For each training dataset, we produced two replicated GWAS datasets for evaluating empirical statistical power. The empirical statistical power was calculated as the proportion of over 5,000 repeated simulations with a *P*-value less than 1 × 10^−5^. Furthermore, we produced two replicated GWAS datasets for each training dataset under the null (where *γ* = 0), resulting in over 90,000 replicates across all settings considered. The type 1 error rates were evaluated using the quantile-quantile plot with a 95% confidence interval.

### Protein-phenotype analysis with five GWAS databases

We applied our BLISS models to over 2,500 complex traits and diseases from five major genetic data resources, including the MVP^30^, GBMI^31^, FinnGen^32^, IEU OpenGWAS^33^, and BBJ^34^ studies. **Table S7** summarized the information of these GWAS summary statistics. The primary data resource was from the MVP Genomics Release 4^30^. The MVP cohort included 8.8% female, with an average age of 61.9 years. Following quality control, phenotypes were extracted from Veterans Affairs electronic health records, including diagnosis codes, lab measures, vital signs, and health survey responses. The dataset we used comprised 1,470 binary and 199 quantitative traits for European ancestry, and 1,187 binary and 181 quantitative traits for African ancestry. Additionally, we analyzed MVP GWAS summary statistics obtained from dbGaP study accession phs001672.v3.p1, which had data from individual studies targeting specific phenotypes for both European and African ancestries. Furthermore, we included Alzheimer’s disease, for which we also had GWAS summary statistics for both European and African ancestries^88,89^. The GBMI GWAS summary statistics were downloaded from https://www.globalbiobankmeta.org/resources and we focused on seven diseases with available GWAS summary statistics across European, African, and Asian ancestries. For stroke, we additionally incorporated data of its various subtypes, obtained from a recent multi-ancestry GWAS^20^. FinnGen GWAS summary statistics were accessed from https://www.finngen.fi/en/access_results (version R9). We focused on 278 clinical outcomes defined by FinnGen, most of which had more than 10,000 cases. We also incorporated three diseases with fewer than 10,000 cases, namely Alzheimer’s disease (*n* cases = 9,301), aortic aneurysm (*n* cases = 7,395), and chronic kidney disease (*n* cases = 9,073). For the IEU OpenGWAS database (https://gwas.mrcieu.ac.uk/datasets), we first downloaded all European phenotypes in the ‘ebi-’ and ‘ieu-’ categories that had more than 5×10^6^ genetic variants and a GWAS sample size exceeding 10,000. Furthermore, we included cancer-related traits with a GWAS sample size greater than 3,000. After manually reviewing each downloaded GWAS dataset, we excluded 31 phenotypes that posed potential issues (e.g., missing an autosome), leading to a finalized list of 309 traits. For the BBJ database, we downloaded GWAS statistics for 220 Asian phenotypes^34^ (https://www.ebi.ac.uk/gwas/publications/34594039). We set a threshold of *n* cases > 2,000 for binary traits, resulting in a selection of 107 phenotypes. In all analyses, we only considered protein-phenotype pairs where the number of SNPs in the test exceeded 10.

When interpreted our findings in our main text, we focused on the MVP Genomics Release 4^30^ data resources and considered 1,183 SomaScan proteins measured in deCODE, 1,214 SomaScan proteins in ARIC-EA, 1,352 SomaScan proteins in ARIC-AA, and 1,407 Olink proteins measured in UKB-PPP. These proteins were selected based on a *cis*-heritability threshold greater than 0.01^4,8,19^ to satisfy the relevance assumption. The association results of all phenotypes using the full set of BLISS models were available through our web tool (https://www.gcbhub.org/).

### Method comparison

Using our primary data resource from the MVP^30^, we systematically compared the results of BLISS models with those from transcriptome-wide association studies (TWAS), Mendelian randomization (MR), and Bayesian colocalization analyses. We conducted TWAS using gene expression imputation models of the whole blood tissue. For the European ancestry analysis, we used models from PrediXcan^42^ trained on GTEx v8, and for the African ancestry analysis, we used models from the GALA study^45^. Next, we conducted an MR analysis. We first restricted our analysis to the common set of variants that were shared by pQTLs and GWAS. To meet the relevance assumption, we required that pQTLs must contain at least one SNP with *P* < 5 × 10^−8^. To avoid the potential issue of collinearity, we applied LD clumping (threshold *r*^2^ = 0.001) to obtain a set of independent SNPs as the candidate instrumental variables. We applied either the Wald ratio (when only one instrumental variable was available) or inverse variance weighting^90^ to test the association between protein and phenotype. We also performed Bayesian colocalization analysis to estimate the posterior probability that the protein and phenotype shared the same causal variant, using summary-level *cis*-pQTLs and GWAS data. Specifically, we used the coloc^91^ package (v5.2.2), with its default setups in the coloc function, to estimate the posterior probability of both protein and phenotype being influenced by the same causal variant (i.e., the PPH4). We chose PPH4 > 0.8^91,92^ as the threshold. Proteins satisfying this threshold would suggest a shared causal variant for the *cis*-pQTLs and GWAS associations.

### Biological validation using known drug targets

We used a reference dataset of known drug target-indication relationships. This dataset was based on a curated list of 2,832 FDA- or EMA-approved therapies compiled by Minikel, et al. ^58^, which integrates drug information from sources including OpenTargets, DrugBank, and FDA. Following the filtering strategy employed by Minikel, et al. ^58^, we excluded several drug categories not typically pursued through common disease GWAS-driven target discovery, including antineoplastic drugs (*n* = 277), anti-infectives and antiparasitics (*n* = 392), hormonal preparations (*n* = 154), and vitamins and analogues (*n* = 90). We also excluded drugs for which the primary protein target was listed as unknown (*n* = 902). This filtering process resulted in a final set of 1,030 drugs, which collectively target 598 unique protein-coding genes.

To prepare for our analysis, we mapped these drug targets to their corresponding protein products and linked them to disease indications relevant to the phenotypes available in our MVP GWAS data resources. This process yielded a total of 3,081 unique drug-protein-phenotype trios where an approved drug is known to modulate a specific protein for the treatment of a particular disease/phenotype (**Table S4**).

For each of the four analytical methods (BLISS, TWAS, MR, and colocalization), we determined the subset of these 3,081 trios that were analyzable (i.e., the proteins/genes that were included in the respective method’s model set). A known drug-protein-phenotype trio was considered ‘validated’ or ‘overlapped’ by a given method if the association met a pre-defined significance threshold: false discovery rate < 5% for BLISS, TWAS, and MR, or PPH4 > 0.8 for colocalization analysis. The validation rate for each method was calculated as the number of overlapped trios divided by the total number of analyzable trios for that method.

## Code availability

Our data analysis pipeline can be accessed at https://github.com/gcb-hub/BLISS. All software and tools utilized in this study are publicly accessible. The final version of code for model development will be available on Zenodo.

## Data availability

The web tool data resources and trained models are available at https://www.gcbhub.org/. GWAS summary statistics used in our analysis can be downloaded or requested via their individual project websites. The individual-level data used in this study can be accessed from the UK Biobank study at https://www.ukbiobank.ac.uk/.

