## Supplementary material for "Large-scale imputation models for multi-ancestry proteome-wide association analysis": supp_information

### Supplementary Note

August 15, 2025

#### CONTENTS


1 **Supplementary Figures** 2

2 **Comparative advancement of PWAS models derived by BLISS** 30

3 **Impact of PWAS model uncertainty on type I error rate and power** 31

4 **Enhanced discovery via multi-ancestry meta-analysis** 33

5 **Cross-platform comparison in protein–phenotype associations for African ancestry** 34

6 **Platform consistency is critical in super learner-based PWAS model building** 35

7 **A practical guide to interpreting PWAS findings and integrating complementary evidence** 35

### 1 Supplementary Figures

#### 1.1 Supplementary Figure 1

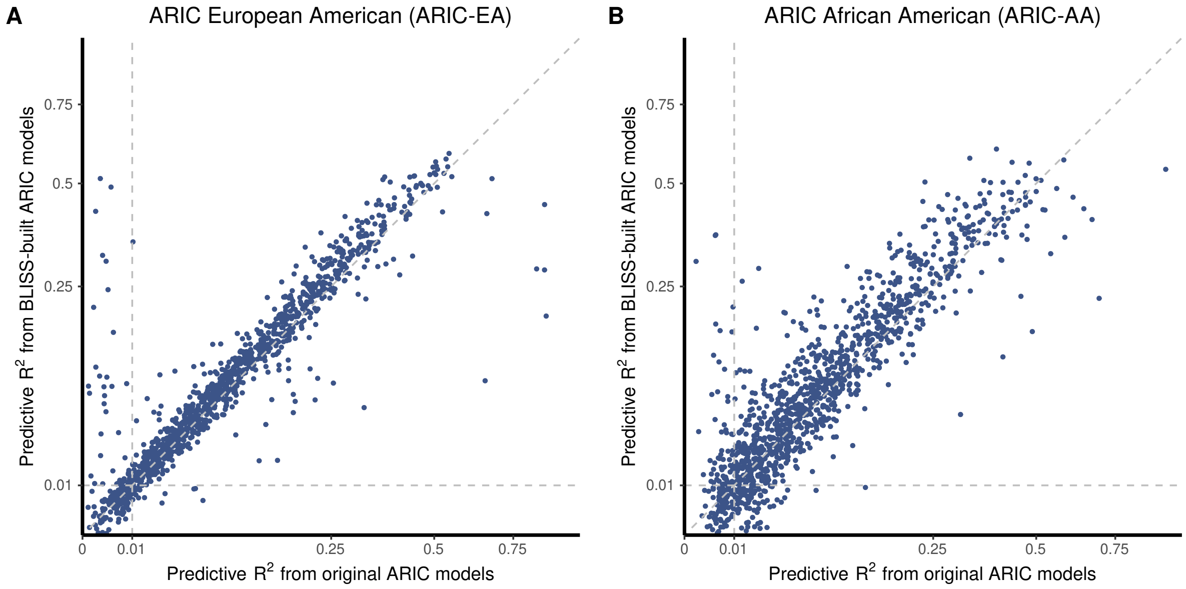


**Supplementary Figure 1: BLISS training improves predictive performance of ARIC protein models across ancestries.** (A) Compares the predictive performance between original ARIC models (x-axis) and BLISS-trained models in ARIC European American study (ARIC-EA). (B) Compares the predictive performance between original ARIC models (x-axis) and BLISS-trained models in ARIC African American study (ARIC-AA). Each point represents a protein; dashed line represents $y = x$.

#### 1.2 Supplementary Figure 2

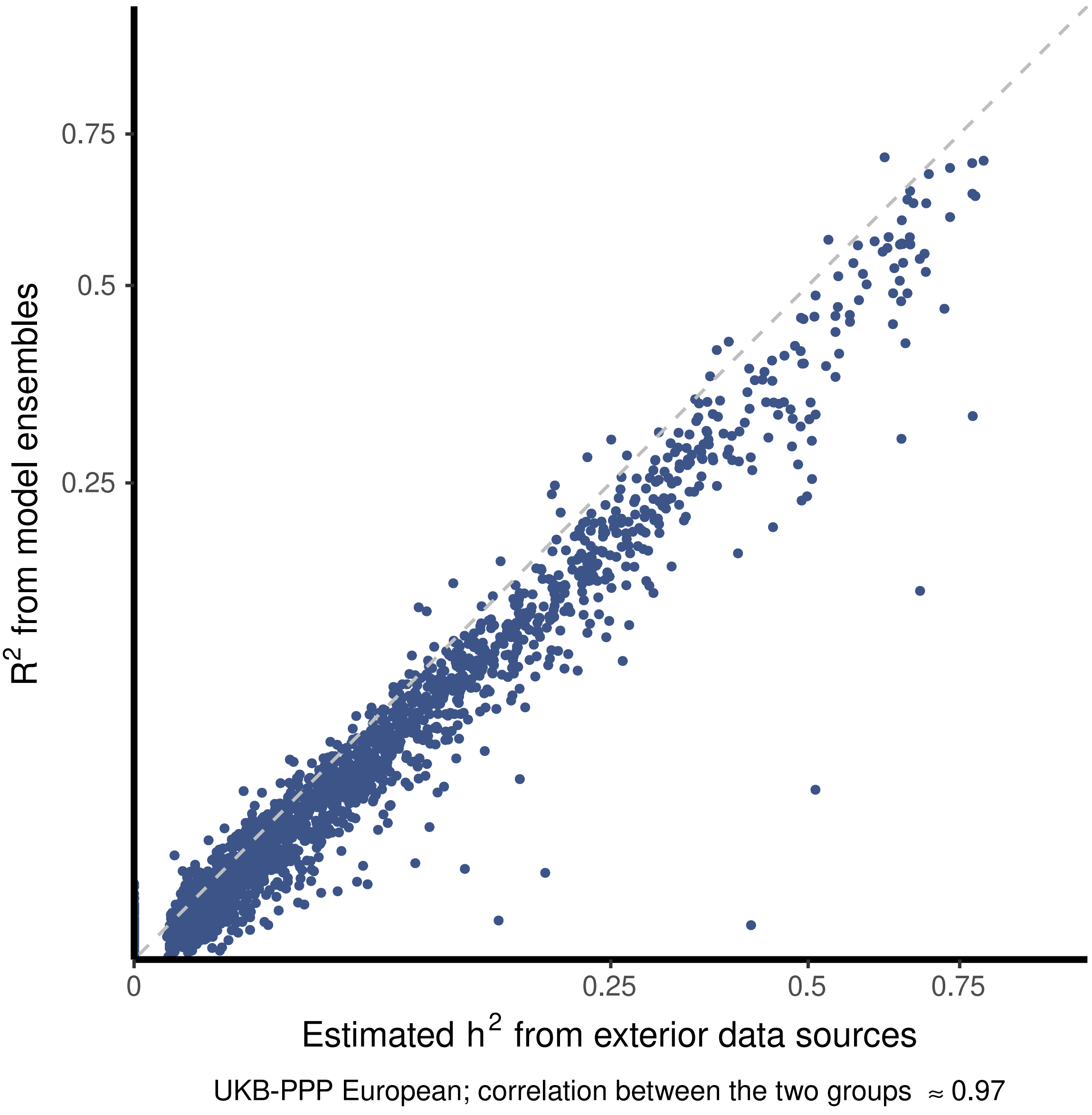


**Supplementary Figure 2: Comparison of predictive performance of BLISS-trained UKB-PPP European models and corresponding estimated *cis*-heritability.** Compares the estimated *cis*-heritability ($h^2$) from [7] (x-axis) and the performance $R^2$ from BLISS-trained UKB-PPP European models (y-axis). Each point represents a protein; dashed line represents $y = x$.

#### 1.3 Supplementary Figure 3

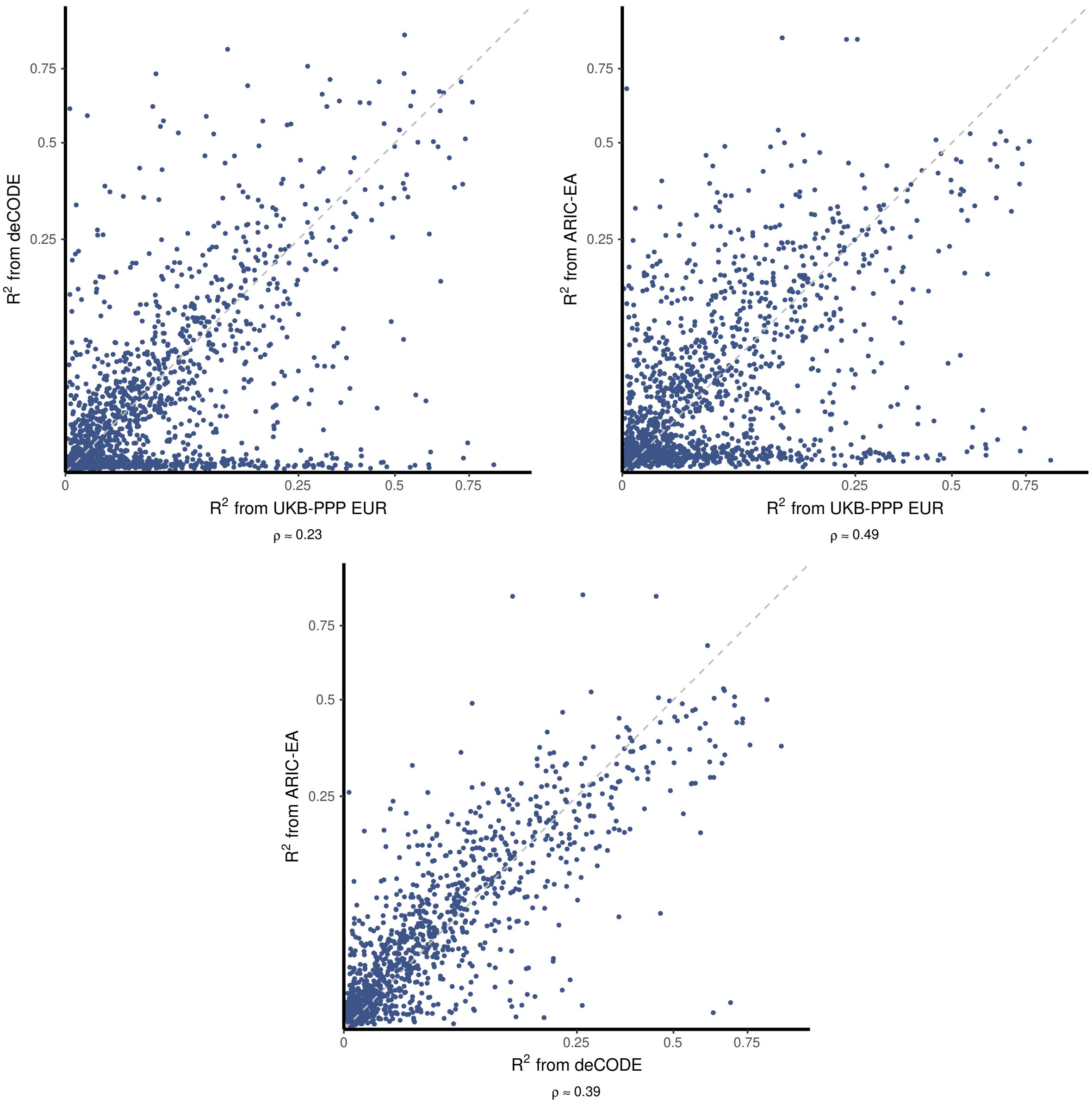


**Supplementary Figure 3: Consistency of predictive performance by BLISS among studies.** Compares the the predictive $R^2$ of BLISS-trained models from different studies: UKB-PPP European (UKB-PPP EUR), deCODE, ARIC European American (ARIC-EA). Each point represents a protein; gray dashed line represents $y = x$.

#### 1.4 Supplementary Figure 4

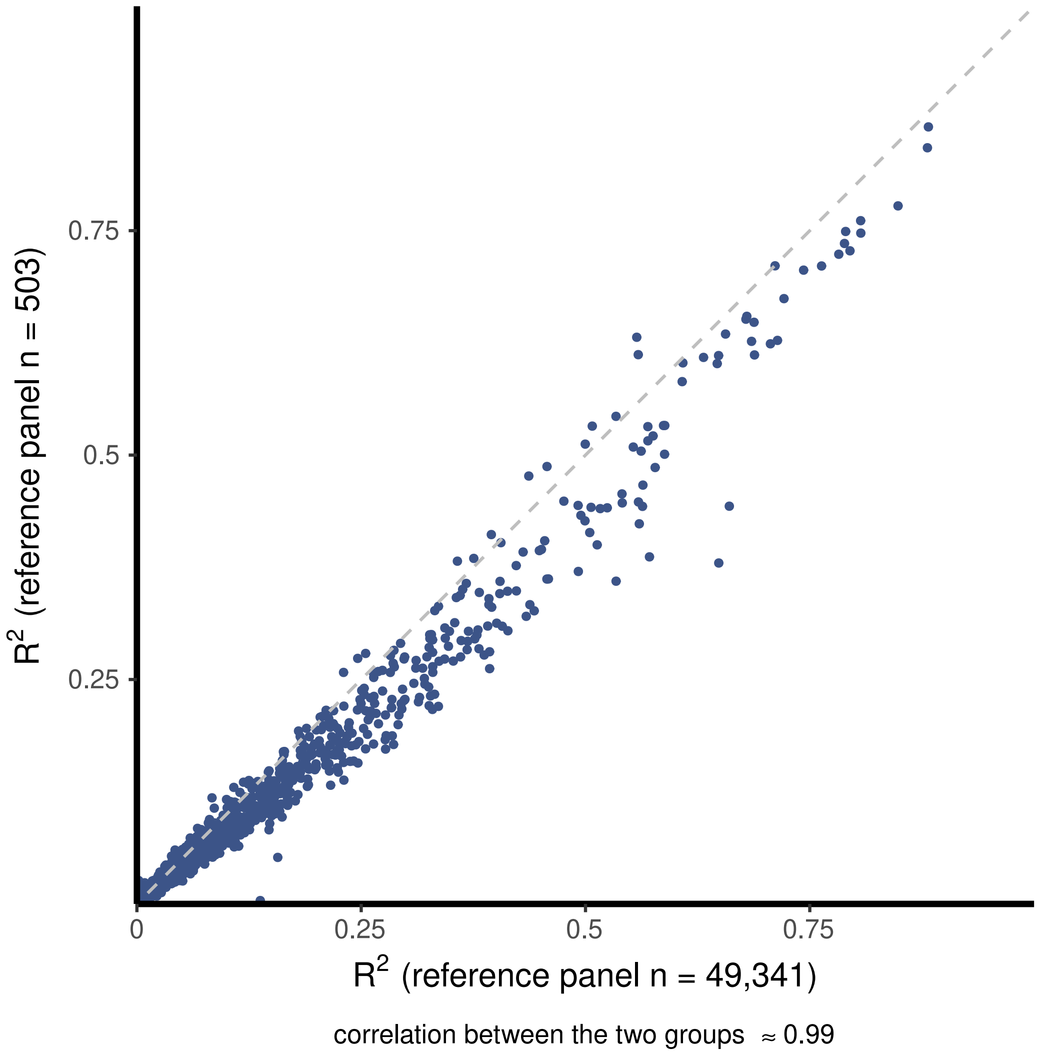


**Supplementary Figure 4: Comparison of predictive performance between models built using larger and smaller reference panels.** Compares the predictive $R^2$ of UKB-PPP European models trained by BLISS leveraging larger (N = 49,341) reference panel (x-axis) and smaller (N = 503) reference panel (y-axis). Both reference panels use genetics data from UKB-PPP European study. Each point represents a protein; gray dashed line represents $y = x$.

#### 1.5 Supplementary Figure 5

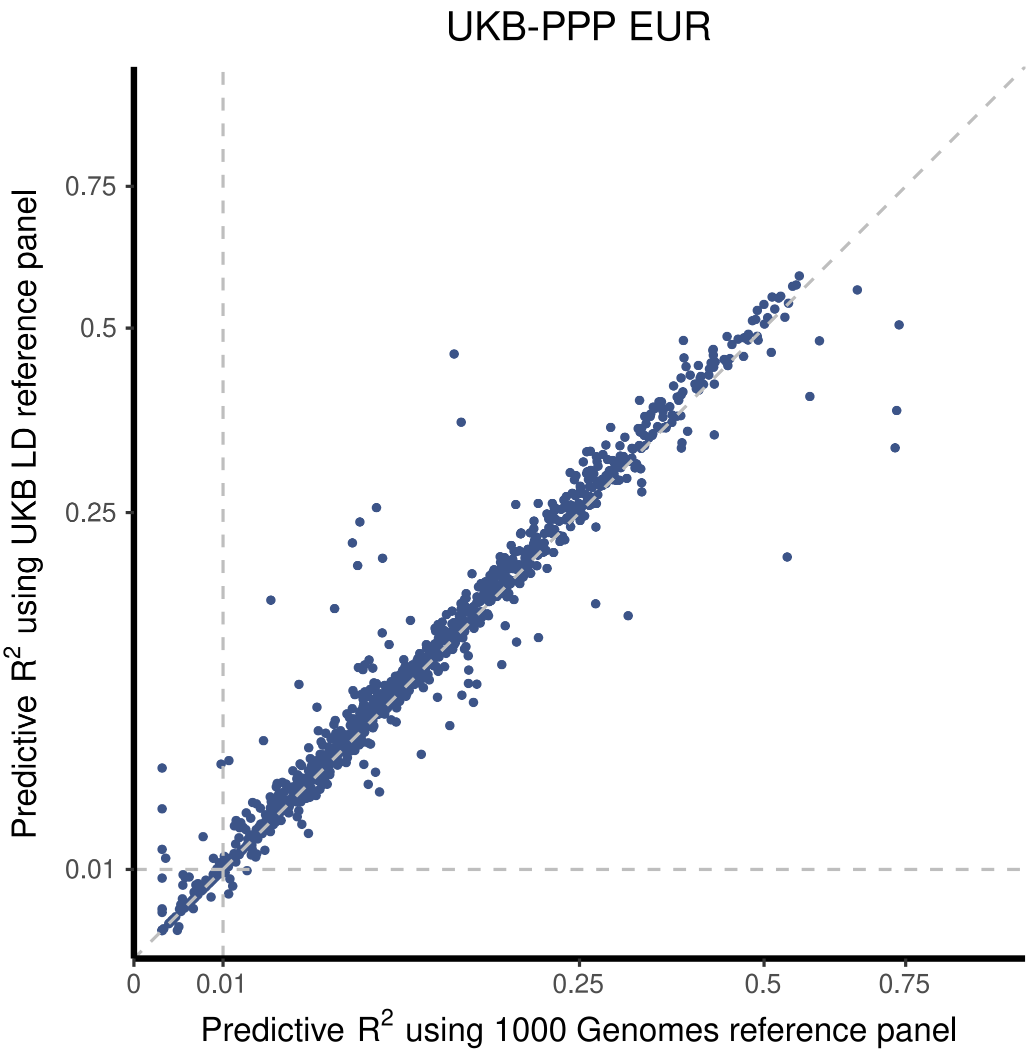


**Supplementary Figure 5: Comparison of predictive performance between models built using reference panels from varying sources.** Compares the predictive performance of UKB-PPP European (UKB-PPP EUR) models trained by BLISS leveraging reference panel from 1000 Genomes Project (N = 503; x-axis) and UKB-PPP EUR (N = 503; y-axis). Each point represents a protein; dashed line represents $y = x$, $x = 0.01$, and $y = 0.01$.

#### 1.6 Supplementary Figure 6

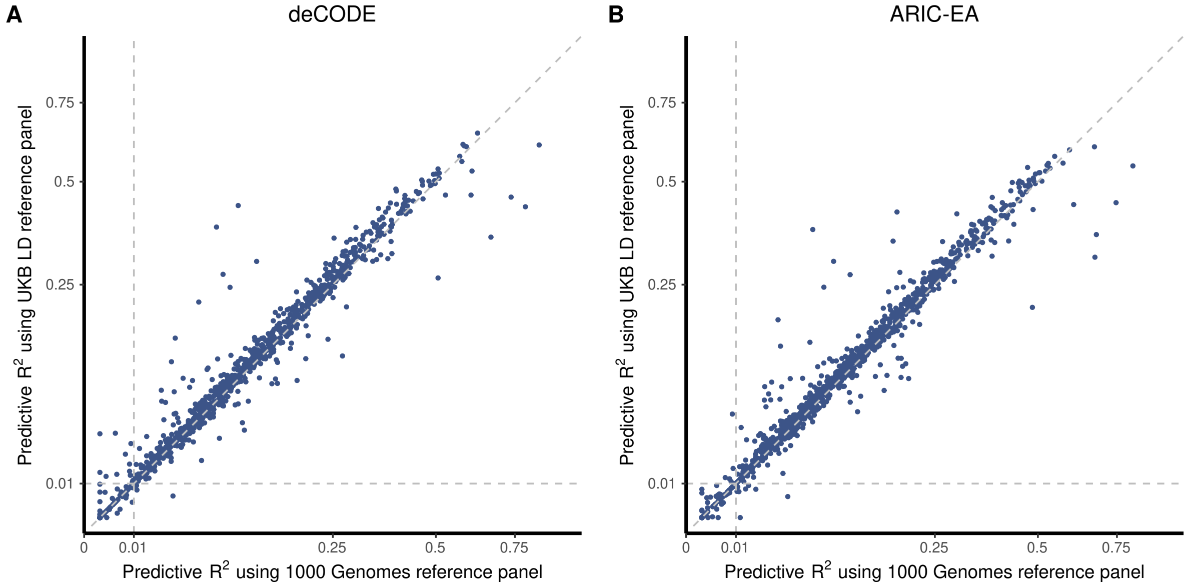


**Supplementary Figure 6: Comparison of predictive performance between models built using reference panels from varying sources in deCODE and ARIC studies.** (A) Compares of predictive performance by deCODE models constructed by BLISS using reference panels from 1000 Genomes Project (N = 503; x-axis) and UKB-PPP European (N = 503; y-axis). (B) Same comparison in ARIC European American (ARIC-EA). Each point represents a protein; dashed line represents $y = x$, $x = 0.01$, and $y = 0.01$.

#### 1.7 Supplementary Figure 7

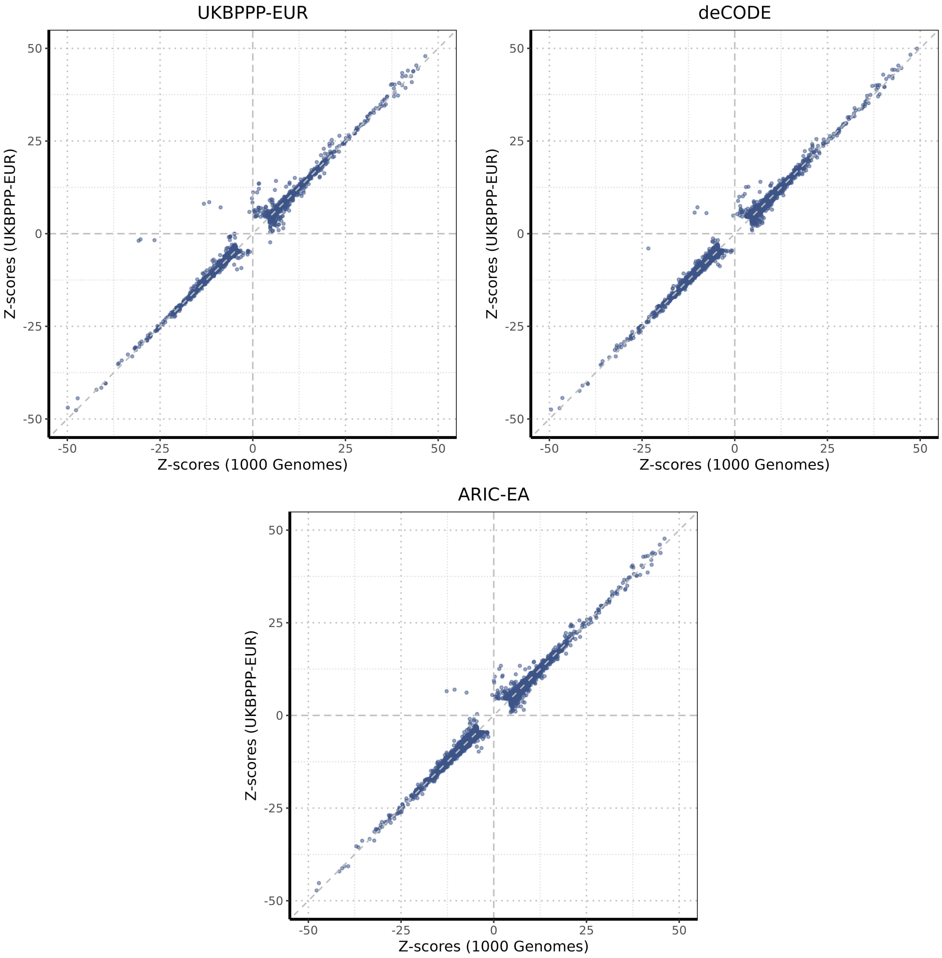


**Supplementary Figure 7: Comparison of association study $\mathbb{Z}$-scores between BLISS models using different reference panels.** Each panel compares $Z$-scores from models using 1000 Genomes Project data (x-axis) versus models using UKB-PPP European (UKB-PPP EUR) data (y-axis) for three cohorts: UKB-PPP EUR (top left), deCODE (top right), and ARIC European American (ARIC-EA) (bottom). Each point represents a protein, with only associations identified by at least one approach included; dashed line represents $y = x$, $x = 0$, and $y = 0$.

#### 1.8 Supplementary Figure 8

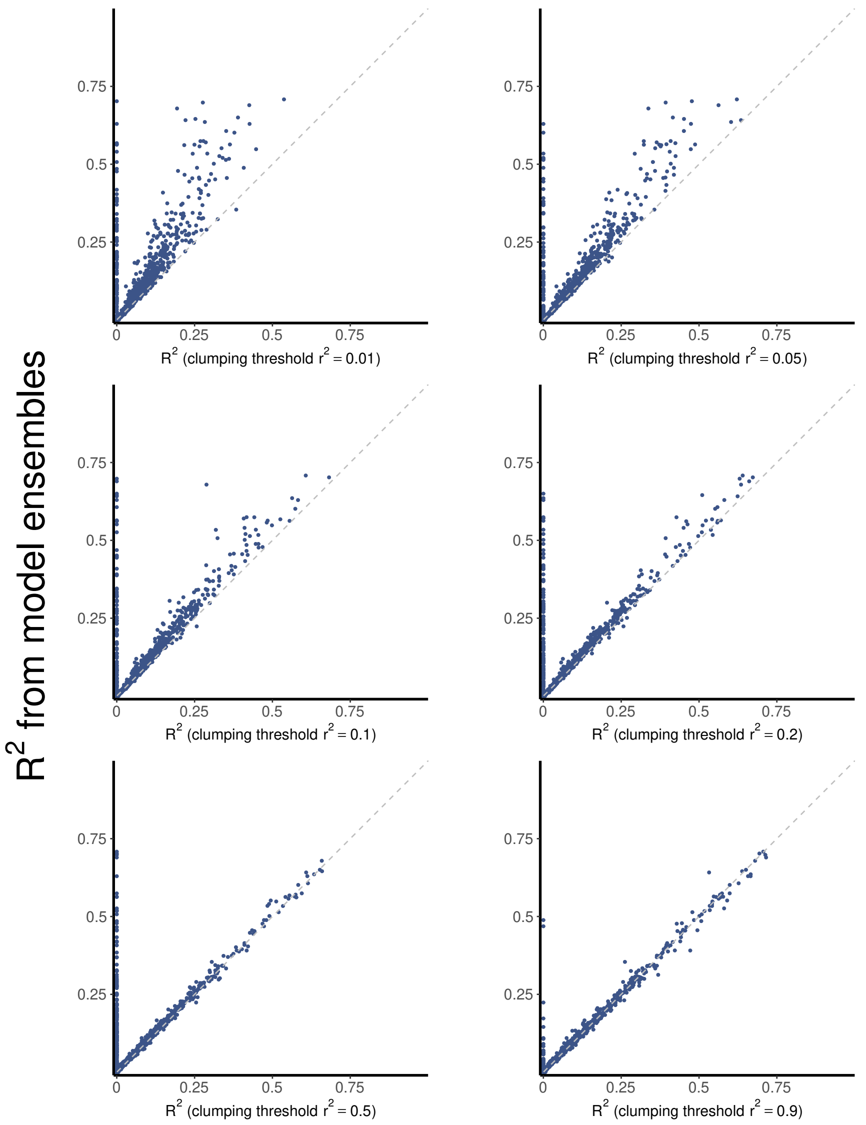


**Supplementary Figure 8: Comparison of predictive performance between model ensembles and models using single LD clumping thresholds.** Each panel compares the predictive performance of models using a single LD clumping threshold (x-axis) versus model ensembles that integrate multiple thresholds (y-axis). The six panels show results for different clumping thresholds: $r^2$ = 0.01, 0.05, 0.1, 0.2, 0.5, and 0.9. Each point represents a protein; dashed line represents $y = x$.

#### 1.9 Supplementary Figure 9

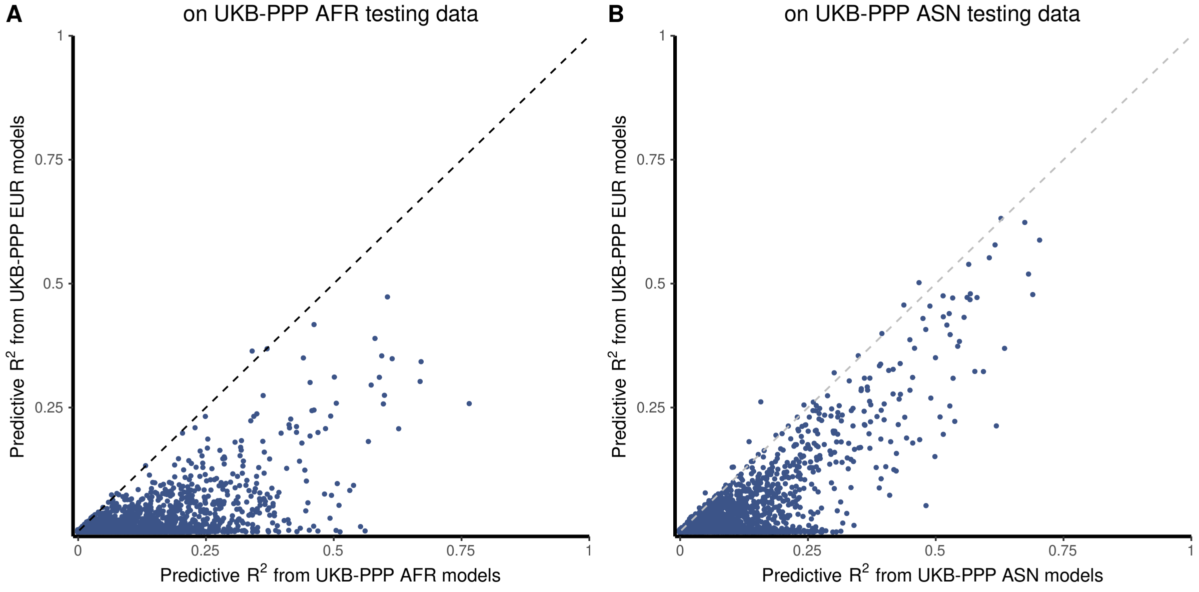


**Supplementary Figure 9: Cross-ancestry evaluation reveals reduced predictive performance with ancestry-mismatched models.** (A) Compares the predictive $R^2$ of UKB-PPP African (UKB-PPP AFR) models (x-axis) and UKB-PPP European (UKB-PPP EUR) models (y-axis) evaluated on standalone UKB-PPP AFR testing data. (B) Same comparison on UKB-PPP Asian (UKB-PPP ASN). Each point represents a protein; dashed line represents $y = x$.

#### 1.10 Supplementary Figure 10

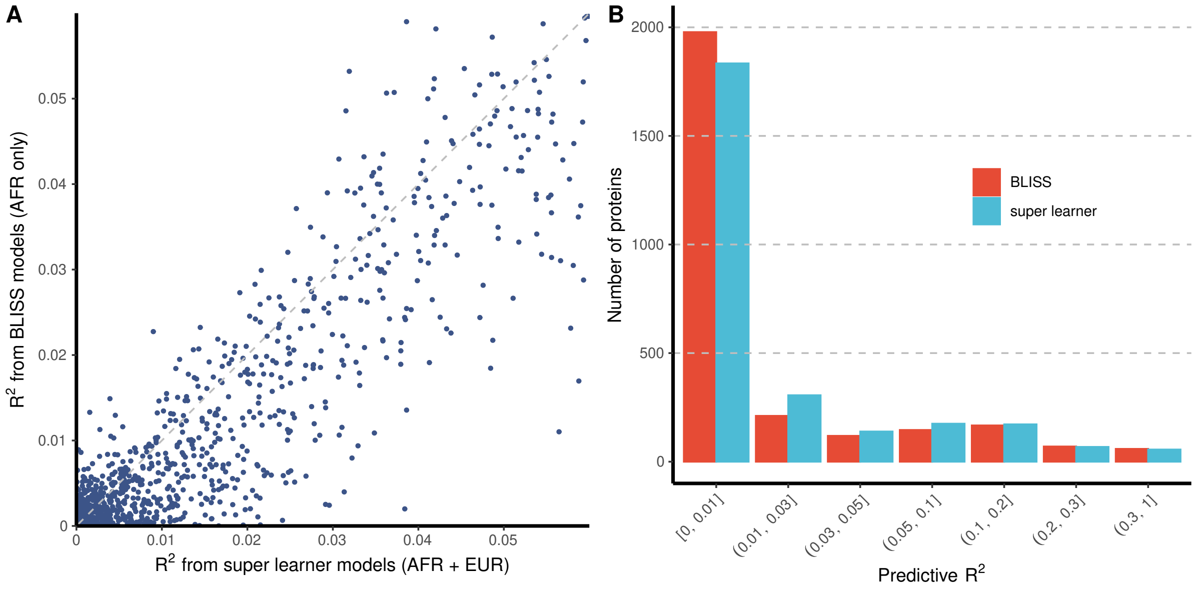


**Supplementary Figure 10: Super learner models combining African and European data outperform regular BLISS models.** (A) Compares the predictive $R^2$ between super learner models (AFR + EUR) (x-axis) and BLISS models trained on UKB-PPP African (AFR) data only (y-axis), with each point representing a protein and dashed line representing $y = x$. (B) Demonstrates the distribution of predictive $R^2$ values for both approaches across binned ranges, with BLISS (red bars) representing the BLISS models and super learner (blue bars) representing super learner models (AFR +EUR).

#### 1.11 Supplementary Figure 11

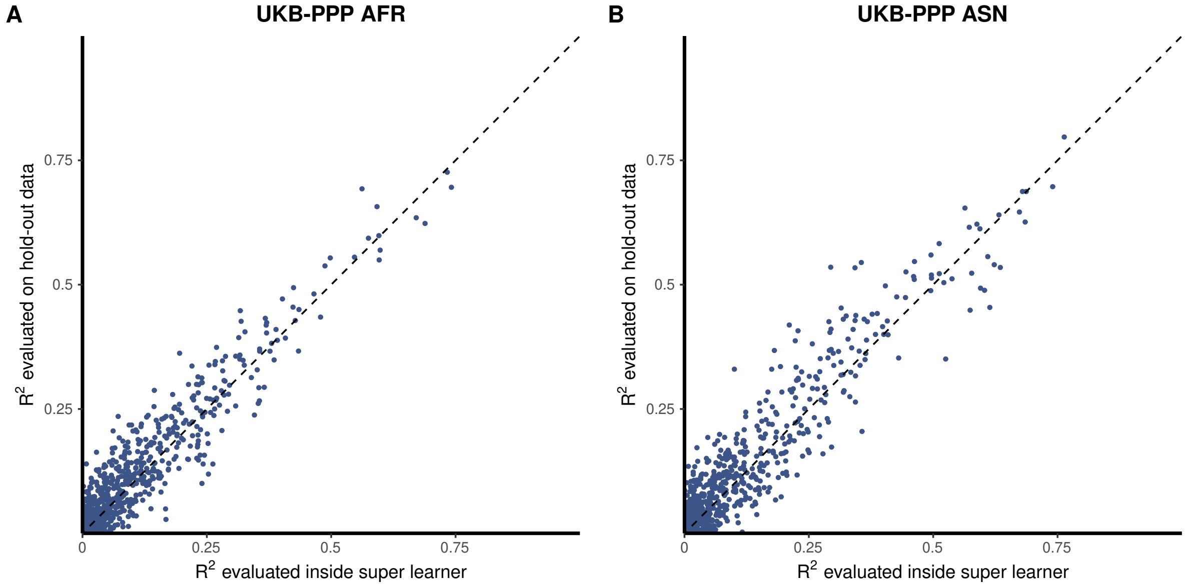


**Supplementary Figure 11: Comparison of predictive performance of super learner models evaluated on hold-out data and internally estimated.** (A) Compares the predictive $R^2$ internally estimated by super learner (x-axis) and the predictive $R^2$ of the UKB-PPP African (UKB-PPP AFR) super learner models evaluated on hold-out data. (B) Same comparison in UKB-PPP Asian (UKB-PPP ASN). The correlation between these two sets of $R^2$ is $0.96$ for UKB-PPP AFR and $0.95$ for UKB-PPP ASN. Each point represents a protein; dashed line represents $y = x$.

#### 1.12 Supplementary Figure 12

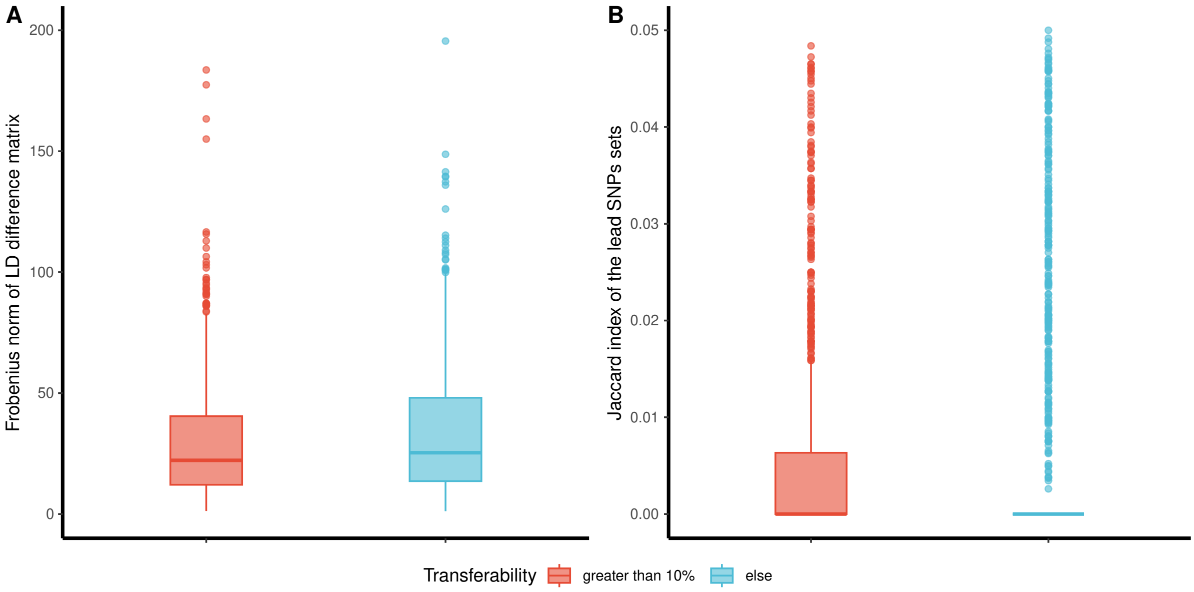


**Supplementary Figure 12: Comparison of quantitative characteristics of proteins with different transferability.** (A) Compares the Linkage Disequilibrium (LD) structures difference (quantified by Frobenius norm of LD difference between UKB-PPP African and European) between proteins with transferability greater than 10% (red) and others (blue). (B) Compares the genetic data alignment (quantified by Jaccard index of SNPs with $p \leq 2.5 \times 10^{-6}$ in the corresponding pQTL summary-level data) between proteins with transferability greater than 10% and others. Transferability is measured as percentage predictive $R^2$ gain in African ancestry models when enhanced by super learner. Box plots show median, quartiles, and whiskers extending to $1.5\times$IQR (interquartile range).

#### 1.13 Supplementary Figure 13

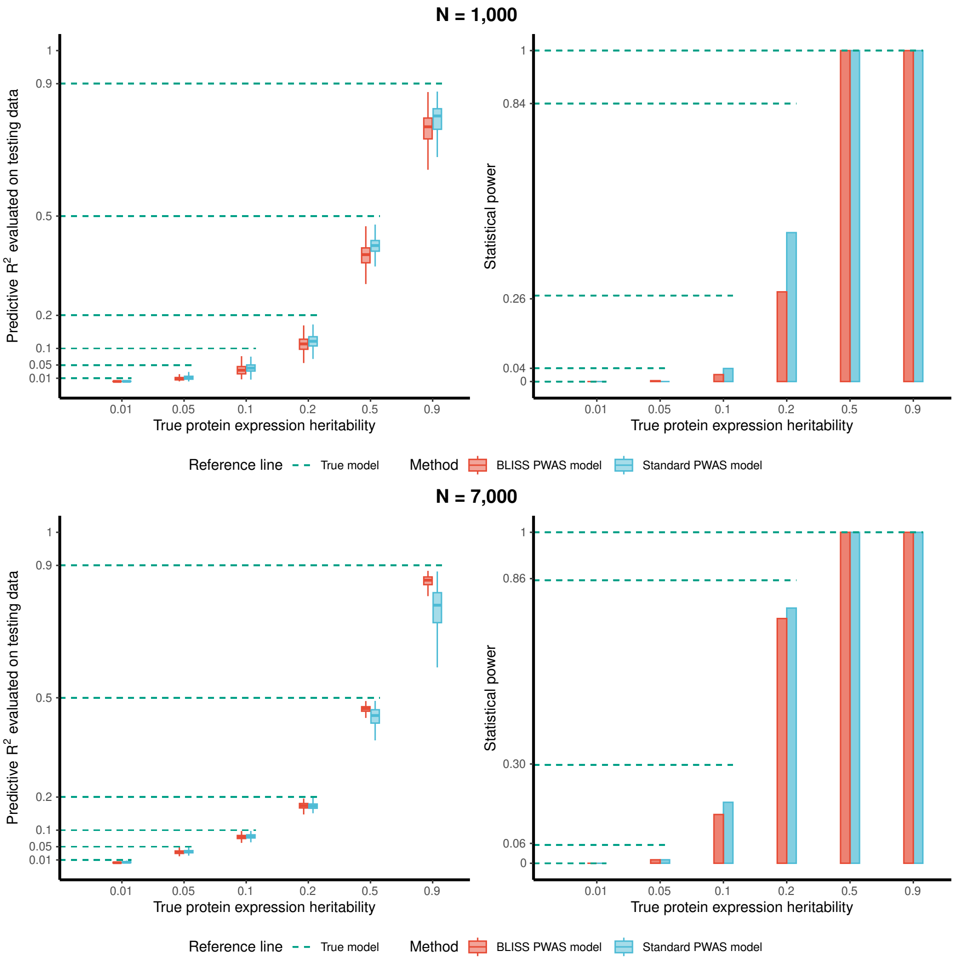


**Supplementary Figure 13: Performance comparison of BLISS and standard PWAS models with 5% causal SNPs.** Left panels show predictive $R^2$ under different preset heritability; right panels show statistical power. Results are presented for pQTL sample sizes of N = 1,000 (top) and N = 7,000 (bottom). BLISS PWAS models (red) were trained using the BLISS pipeline on summary-level data, while standard PWAS models (blue) were trained using Elastic Net on individual-level data. Horizontal dashed lines (green) indicate the theoretical heritability and power of the true underlying model. Box plots display the median, interquartile range (IQR), and whiskers extending to $1.5\times$IQR. This analysis complements the results presented in **Figs. 3A-3B** by examining model performance across different pQTL sample sizes.

#### 1.14 Supplementary Figure 14

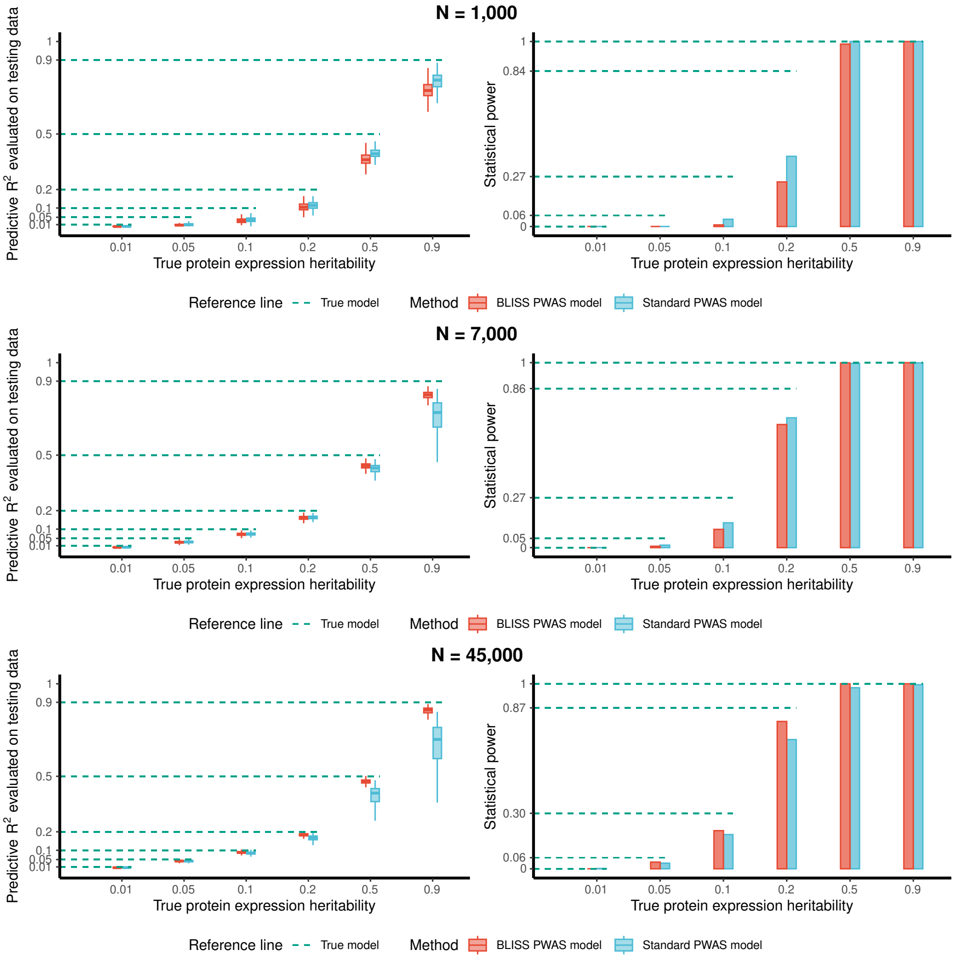


**Supplementary Figure 14: Performance comparison of BLISS and standard PWAS models with 10% causal SNPs.** Left panels show predictive $R^2$ under different preset heritability; right panels show statistical power. Results are presented for pQTL sample sizes of N = 1,000 (top), N = 7,000 (middle), and N = 45,000 (bottom). BLISS PWAS models (red) were trained using the BLISS pipeline on summary-level data, while standard PWAS models (blue) were trained using Elastic Net on individual-level data. Horizontal dashed lines (green) indicate the theoretical heritability and power of the true underlying model. Box plots display the median, interquartile range (IQR), and whiskers extending to $1.5\times$IQR. This analysis complements the results presented in **Figs. 3A-3B** by examining model performance with different casual SNPs percentage.

#### 1.15 Supplementary Figure 15

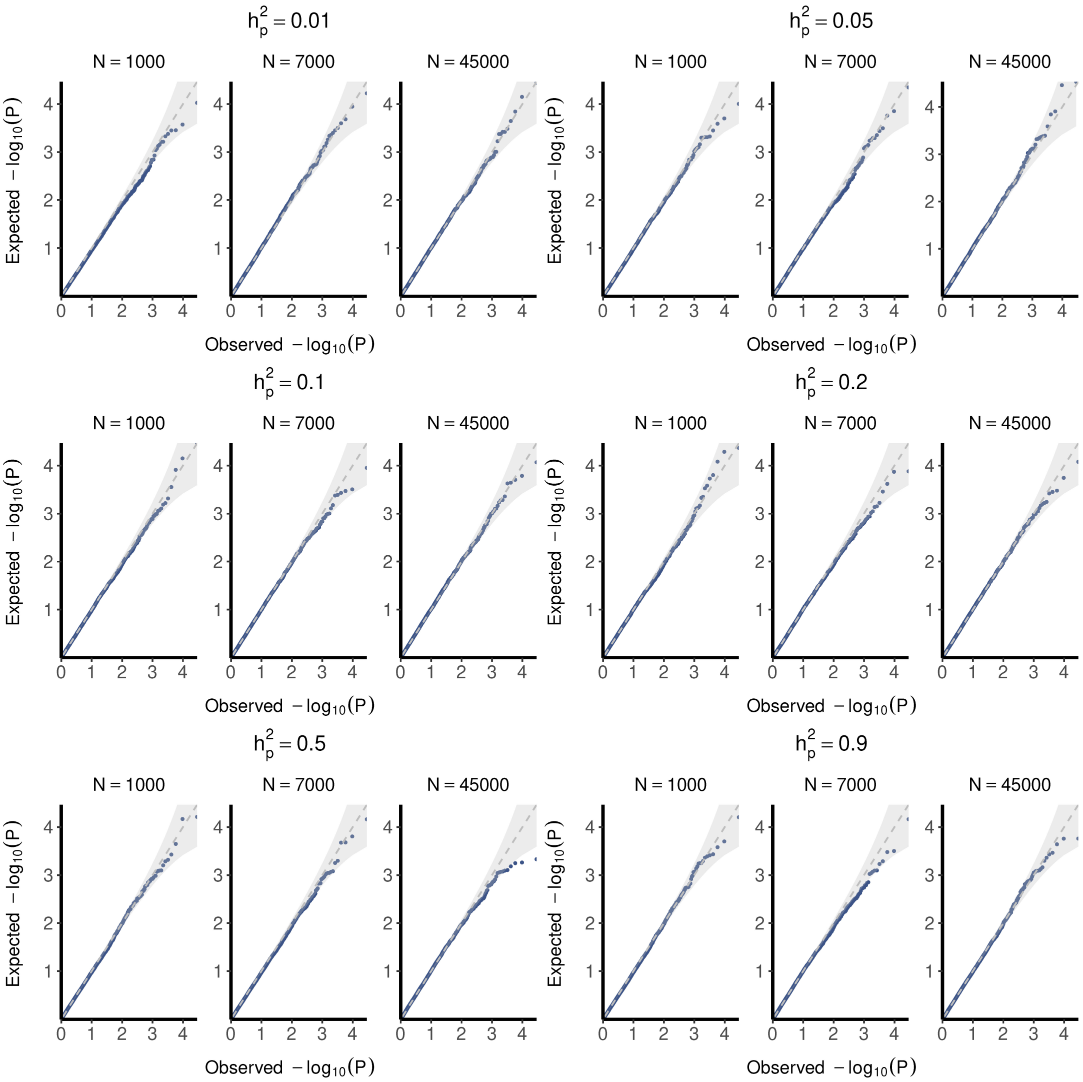


**Supplementary Figure 15: Quantile-quantile plots evaluating type I error control across heritability scenarios (with matching reference panel).** Quantile-quantile plots compare observed versus expected $-\log_{10}(P)$ values for BLISS models under the null hypothesis of no association. Results are stratified by protein expression heritability ($h_p^2 =$ 0.01, 0.05, 0.1, 0.2, 0.5, 0.9) and sample size (N = 1,000, 7,000, 45,000). Models were constructed using matching UKB-PPP European genetics data as the reference panel. Each point represents a protein, with gray shaded areas indicating 95% confidence intervals.

#### 1.16 Supplementary Figure 16

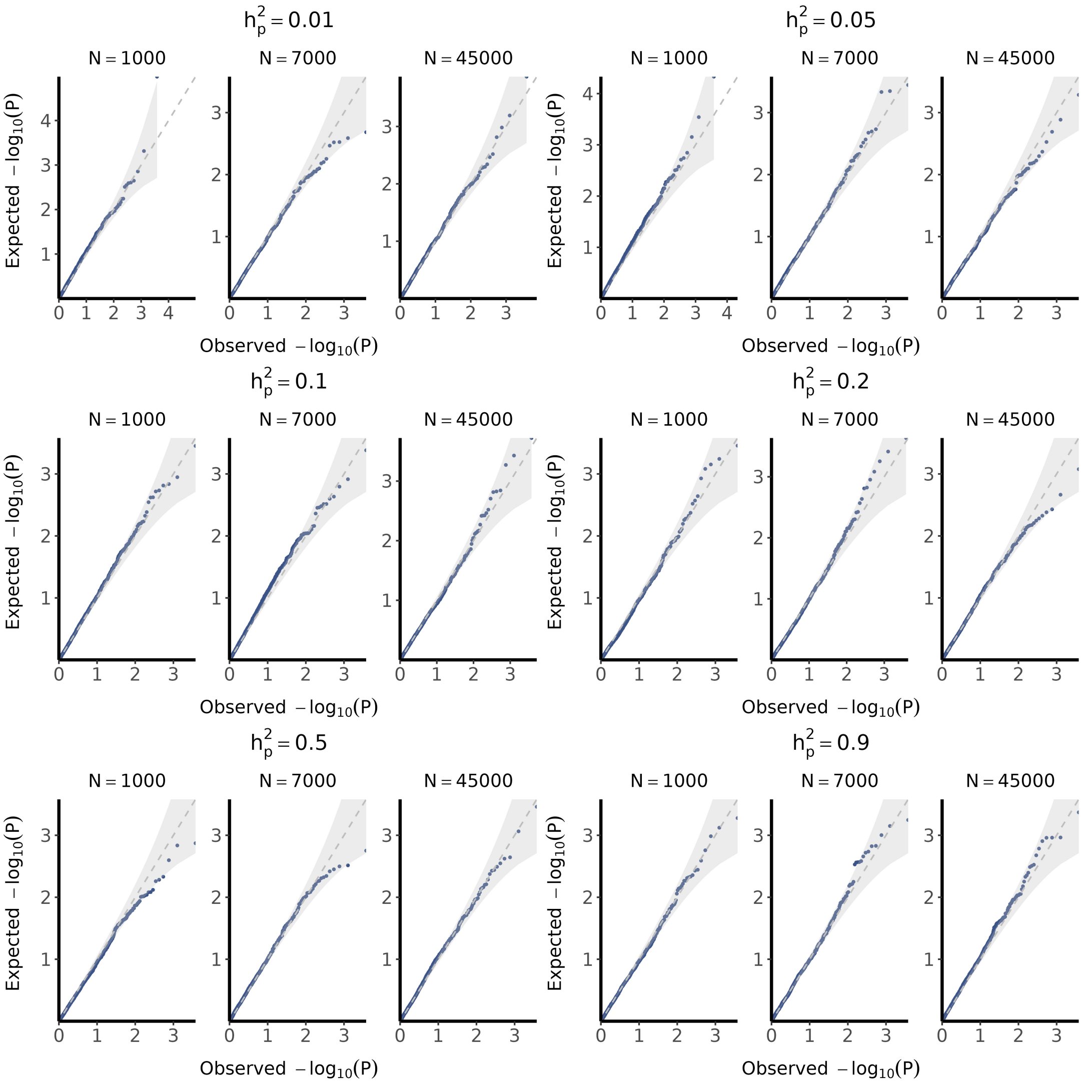


**Supplementary Figure 16: Quantile-quantile plots evaluating type I error control across heritability scenarios (with 1000 Genomes reference panel).** Quantile-quantile plots compare observed versus expected $-\log_{10}(P)$ values for BLISS models under the null hypothesis of no association. Results are stratified by protein expression heritability ($h_p^2 =$ 0.01, 0.05, 0.1, 0.2, 0.5, 0.9) and sample size (N = 1,000, 7,000, 45,000). Models were constructed using 1000 Genomes data (N = 503) as the reference panel. Each point represents a protein, with gray shaded areas indicating 95% confidence intervals.

#### 1.17 Supplementary Figure 17

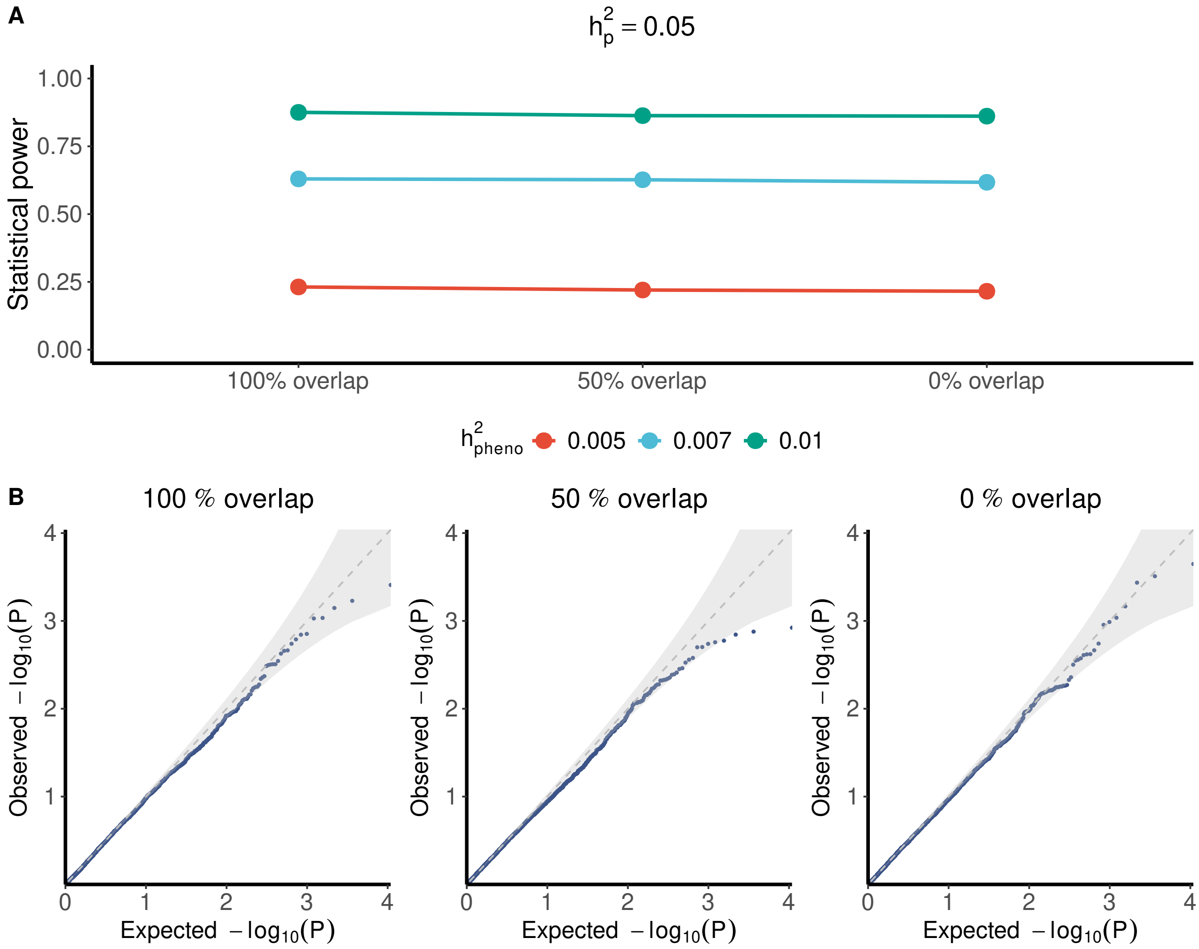


**Supplementary Figure 17: Impact of sample overlap between PWAS training and GWAS data on statistical performance.** (A) Compares the statistical power for detecting protein-phenotype associations remains stable across varying degrees of sample overlap (100%, 50%, and 0%; percentage of PWAS model training data included in phenotype GWAS summary data). Results are shown for three phenotypic heritability levels ($h^2_{pheno}$= 0.005, 0.007, 0.01) with protein expression heritability fixed at $h^2_p = 0.05$. (B) Quantile-quantile plots evaluating type I error control under different sample overlap scenarios. Each panel shows the relationship between observed and expected $-\log_{10}(P)$ values under the null hypothesis. Each point represents a protein, with gray shaded areas indicating 95% confidence intervals.

#### 1.18 Supplementary Figure 18

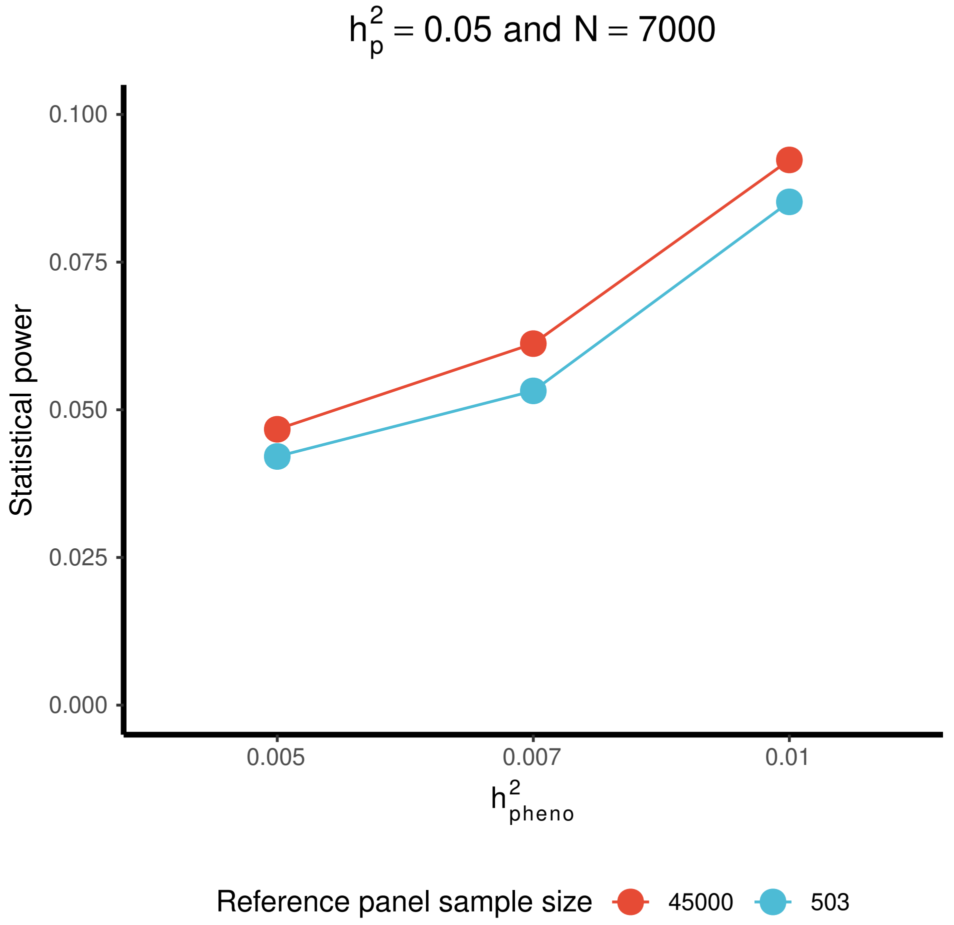


**Supplementary Figure 18: Statistical power comparison between PWAS models trained with different reference panel sample sizes.** Power to detect protein-phenotype associations across phenotypic heritability levels ($h^2_{pheno}$) for models trained with large (N = 45,000, red) versus small (N = 503, blue) reference panels. Protein expression heritability $h^2_p = 0.05$ and GWAS sample size N = 7,000.

#### 1.19 Supplementary Figure 19

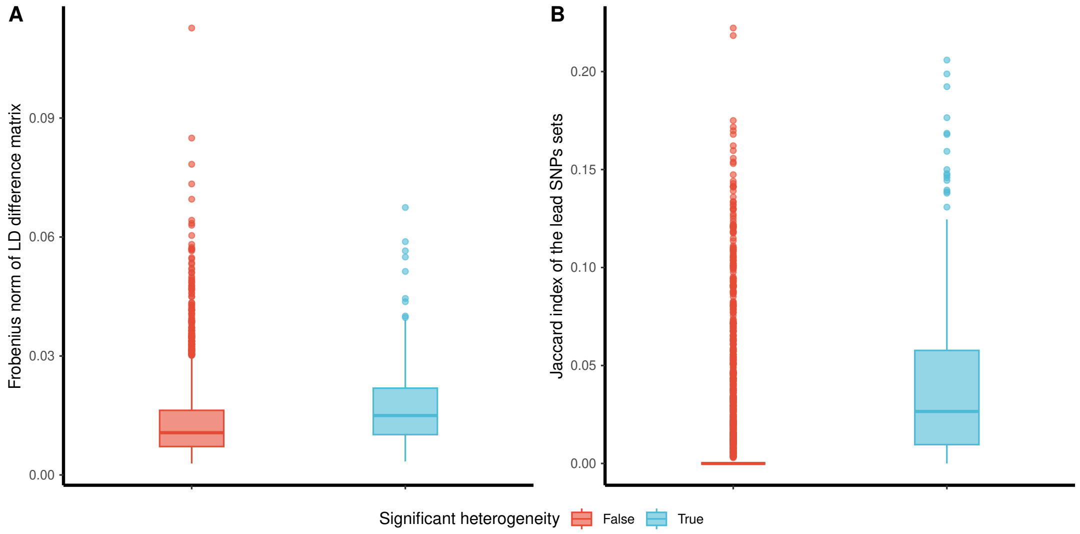


**Supplementary Figure 19: Comparison of proteins with and without significant cross-ancestry heterogeneity.** Proteins were stratified by the presence of significant heterogeneity between ancestry groups using Cochran's Q test (see **Table S2**).(A) compares the Linkage Disequilibrium (LD) structures between proteins with and without significant heterogeneity across UKB-PPP European and African, quantified by Frobenius norm of LD difference. (B) Compares the genetic data alignment between proteins with and without significant heterogeneity across UKB-PPP European and African, quantified by Jaccard index of lead SNPs ($p \leq 2.5 \times 10^{-6}$ in the corresponding pQTL summary-level data). Box plots show median, quartiles, and whiskers extending to $1.5\times$IQR (interquartile range).

#### 1.20 Supplementary Figure 20

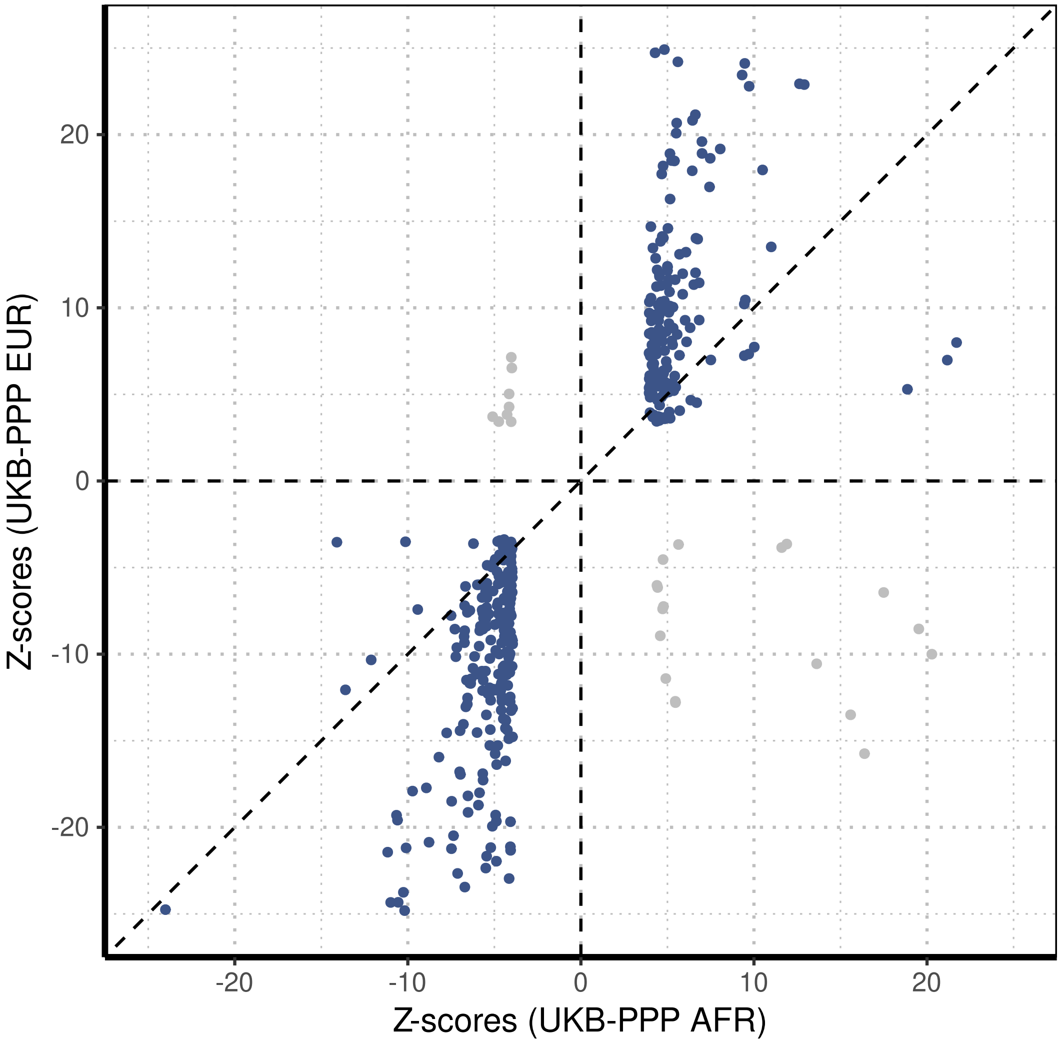


**Supplementary Figure 20: Comparison of genetic association $\mathbb{Z}$-scores between European and African ancestry populations.** Scatter plot comparing $Z$-scores from UKB-PPP European (UKB-PPP EUR) and African (UKB-PPP AFR) ancestry groups for associations achieving statistical significance (FDR < 0.05) in both populations. Blue points indicate concordant effect directions, gray points indicate discordant effects. Dashed lines represent $y = x$, $x = 0$, and $y = 0$.

#### 1.21 Supplementary Figure 21

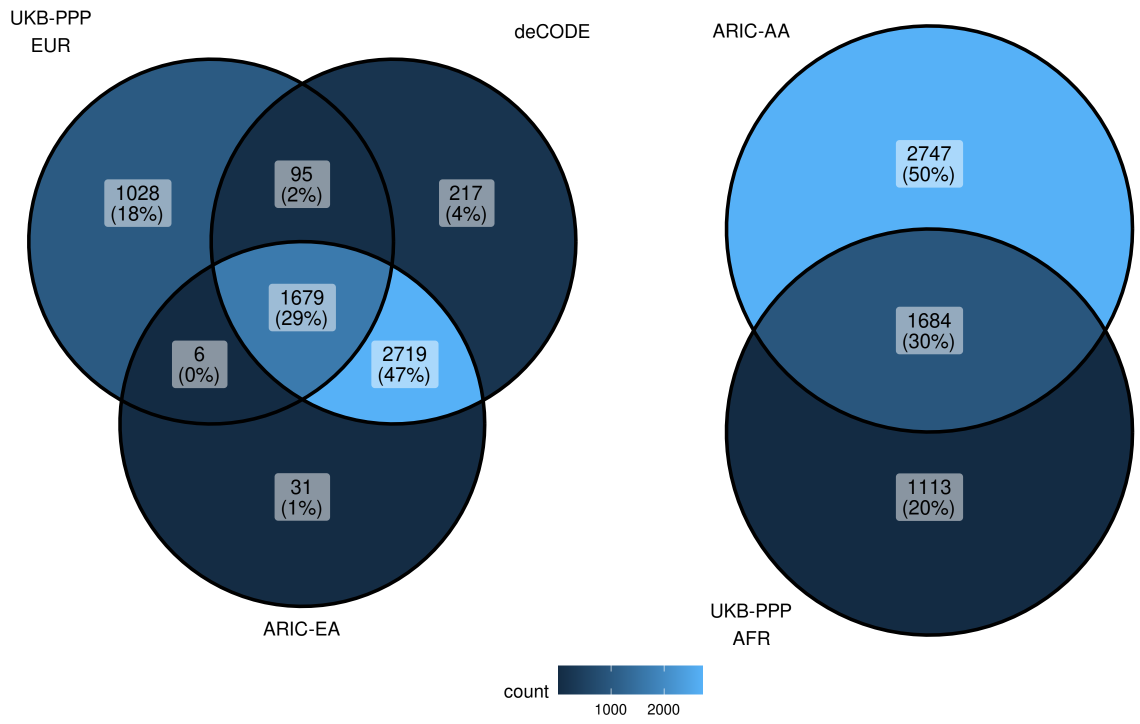


**Supplementary Figure 21: Overlap of protein-trait associations across studies and ancestry groups.** Venn diagrams show the number and percentage of shared significant protein-trait associations between studies. Left panel compares three studies in populations of European ancestry: UKB-PPP European (UKB-PPP EUR), deCODE, and ARIC European American (ARIC-EA). Right panel compares associations between UKB-PPP African (UBK-PPP AFR) and ARIC African American (ARIC-AA) populations. Numbers indicate association counts, with percentages showing the proportion of total associations within each overlap category. Color intensity reflects association density according to the scale bar.

#### 1.22 Supplementary Figure 22

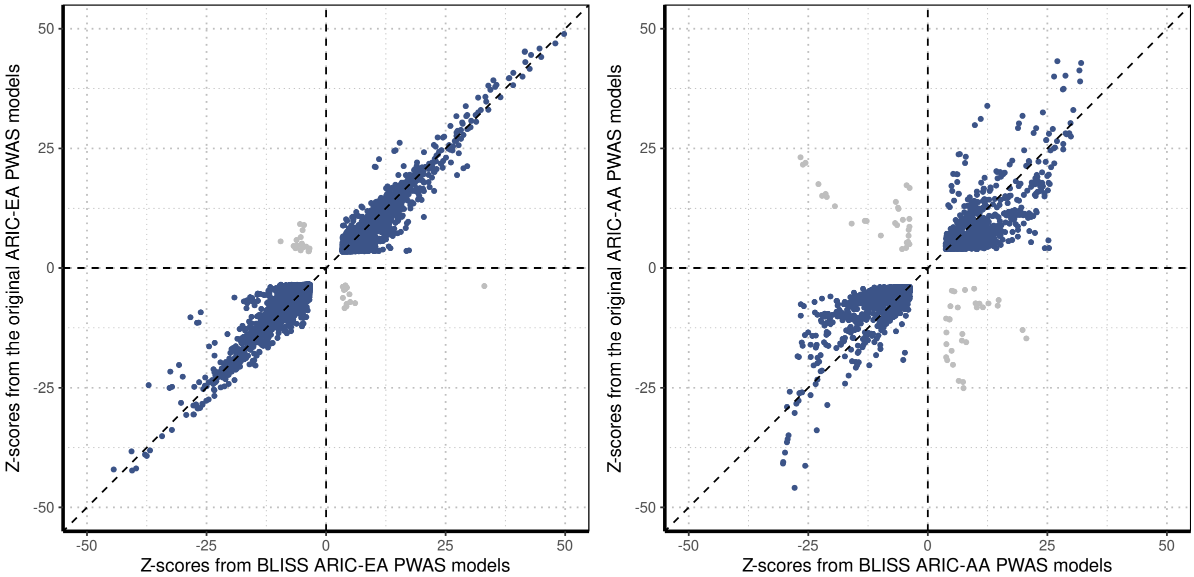


**Supplementary Figure 22: Comparison of $\mathbb{Z}$-scores between BLISS-trained and original PWAS models in the ARIC study.** Scatter plots compare $Z$-scores for significant protein-trait associations derived from BLISS-trained PWAS models (x-axis) versus original ARIC PWAS models (y-axis) stratified by ancestry. Left panel shows ARIC European American (ARIC-EA) comparisons, right panel shows ARIC African American (ARIC-AA) comparisons. Blue points indicate associations with concordant effect directions between methods and gray points indicate discordant effects. Dashed lines represent $y = x$, $x = 0$, and $y = 0$.

#### 1.23 Supplementary Figure 23

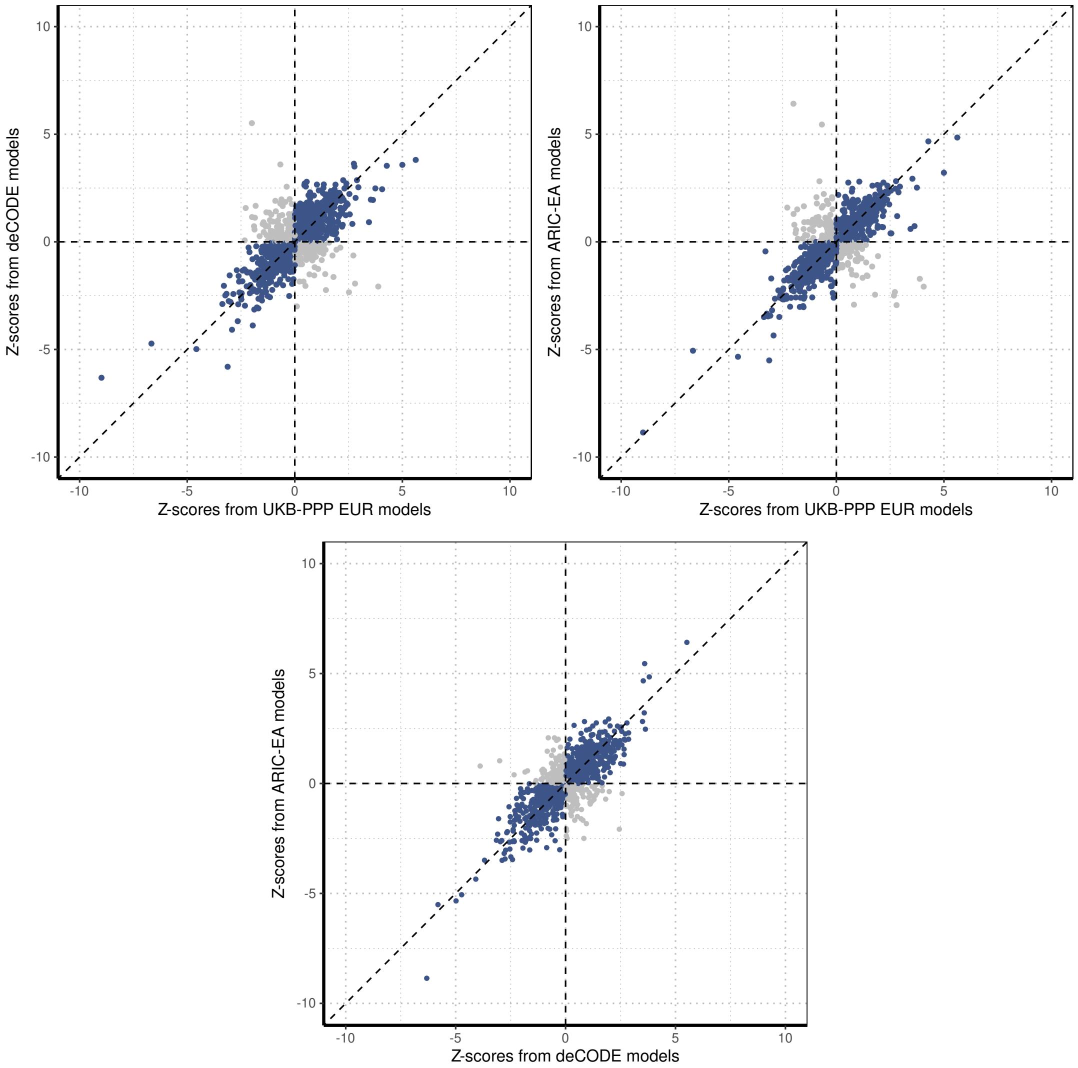


**Supplementary Figure 23: Comparison of $Z$-scores of associations between two proteomics platforms.** Scatter plots compare $Z$-scores among UKB-PPP European (UKB-PPP EUR; Olink), deCODE (SomaScan), and ARIC European American (ARIC-EA; SomaScan). Blue points indicate associations with concordant effect directions between methods and gray points indicate discordant effects. Dashed lines represent $y = x$, $x = 0$, and $y = 0$.

#### 1.24 Supplementary Figure 24

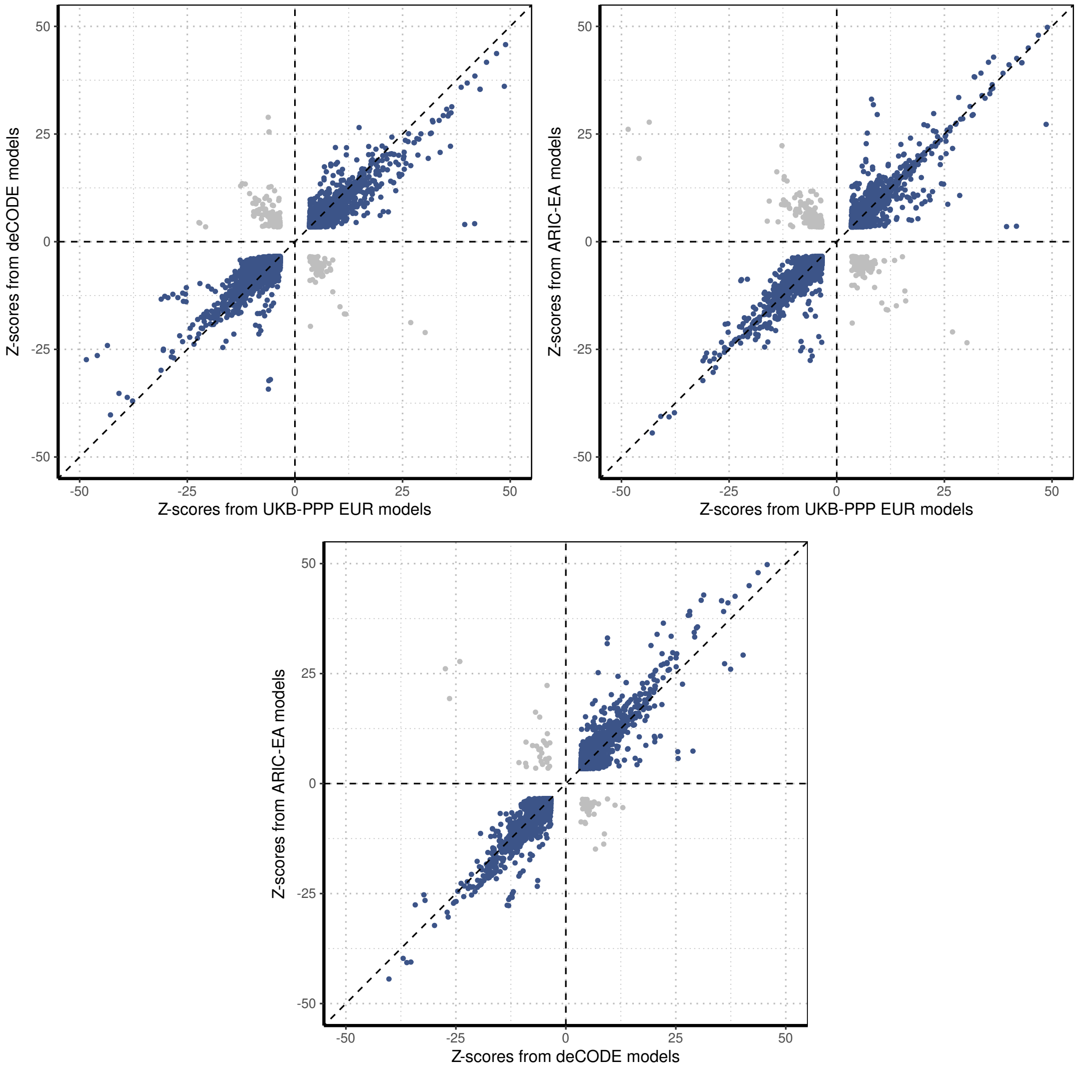


**Supplementary Figure 24: Comparison of Z-scores for shared associations across proteomics platforms.** Scatter plots compare $Z$-scores for protein-trait associations identified as significant in all three cohorts: UKB-PPP European (Olink platform), deCODE (SomaScan platform), and ARIC European American (SomaScan platform).Blue points indicate associations with concordant effect directions between methods and gray points indicate discordant effects. Dashed lines represent $y = x$, $x = 0$, and $y = 0$.

#### 1.25 Supplementary Figure 25

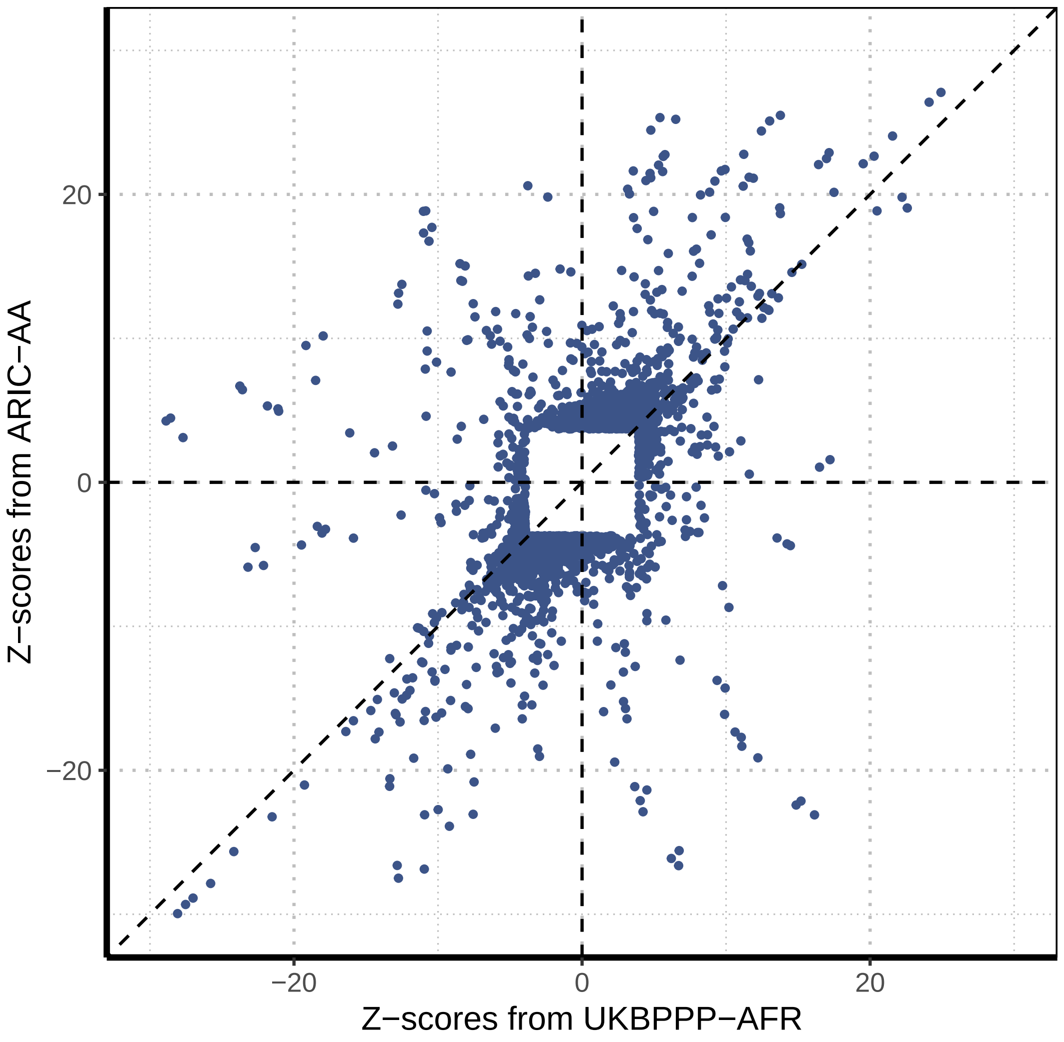


**Supplementary Figure 25: Significant associations identified by either UKB-PPP African or ARIC African American.** Compares the association studies' $Z$-scores of BLISS-trained model from UKB-PPP African (UKB-PPP AFR; x-axis) and from ARIC African American (ARIC-AA; y-axis). Associations identified significant in at least one study were included. Blue points indicate associations with concordant effect directions between methods and gray points indicate discordant effects. Dashed lines represent $y = x$, $x = 0$, and $y = 0$.

#### 1.26 Supplementary Figure 26

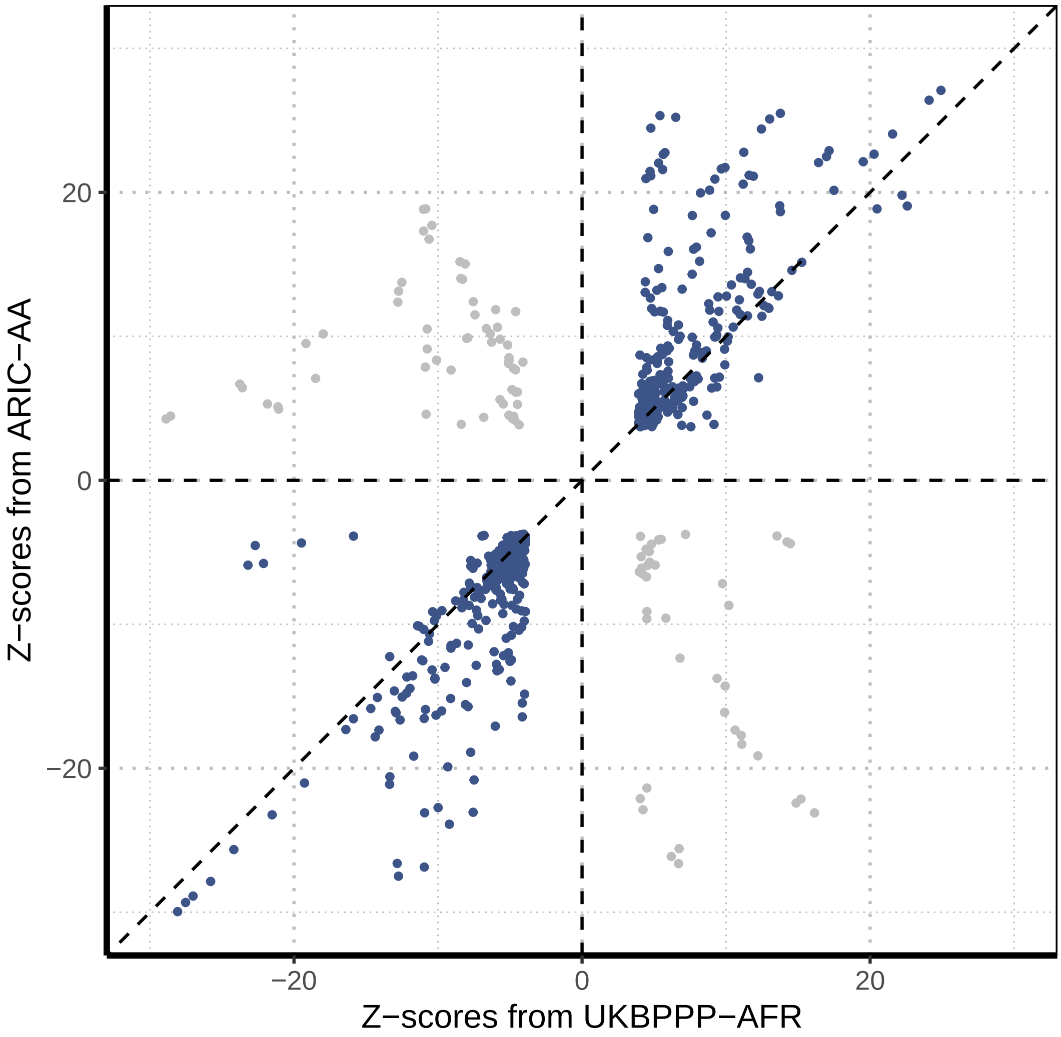


**Supplementary Figure 26: Significant associations identified by both UKB-PPP African and ARIC African American.** Compares the association studies' $Z$-scores of BLISS-trained model from UKB-PPP African (UKB-PPP AFR; x-axis) and from ARIC African American (ARIC-AA; y-axis). Associations identified significant in both studies were included. Blue points indicate associations with concordant effect directions between methods and gray points indicate discordant effects. Dashed lines represent $y = x$, $x = 0$, and $y = 0$.

#### 1.27 Supplementary Figure 27

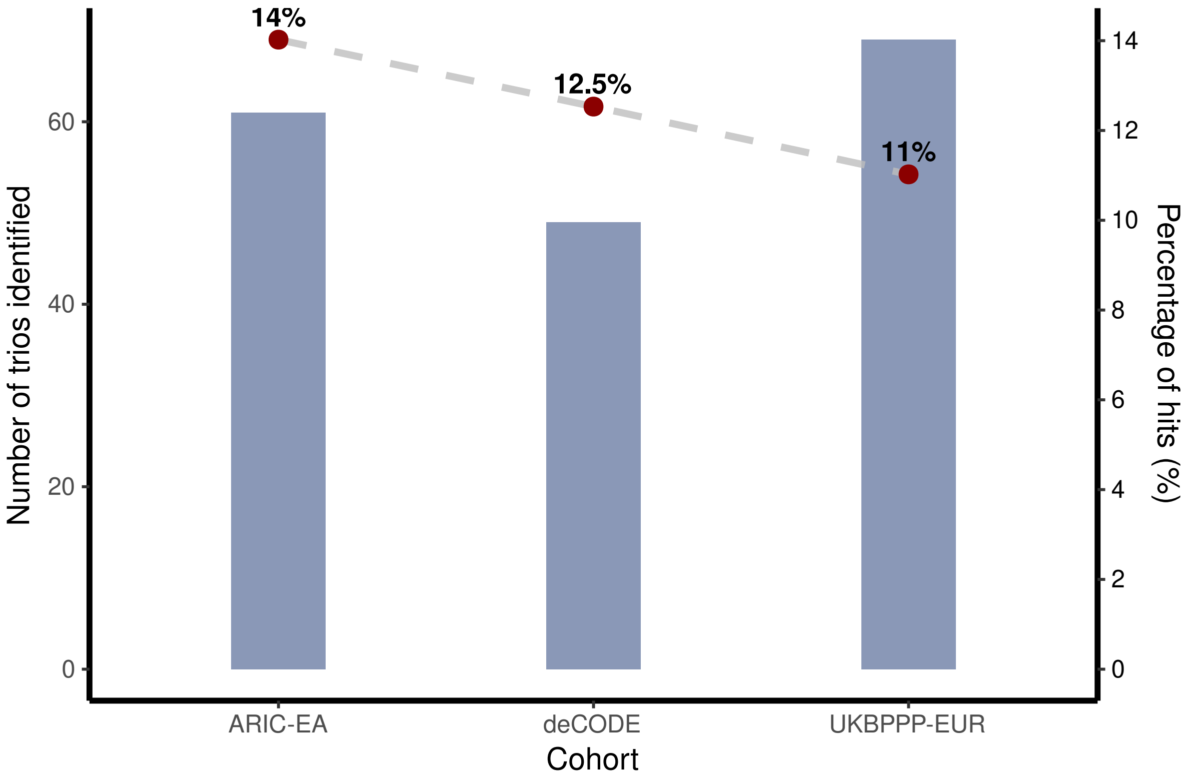


**Supplementary Figure 27: Performance comparison of drug target identification between Olink and SomaScan platform.** The blue bars (measured by the y-axis on the left side) stand for the number of trios found by ARIC European American (ARIC-EA), deCODE, and UKB-PPP European (UKB-PPP EUR). The red dots (measured by the y-axis on the right side) represent the percentage of successfully identified trios.

#### 1.28 Supplementary Figure 28

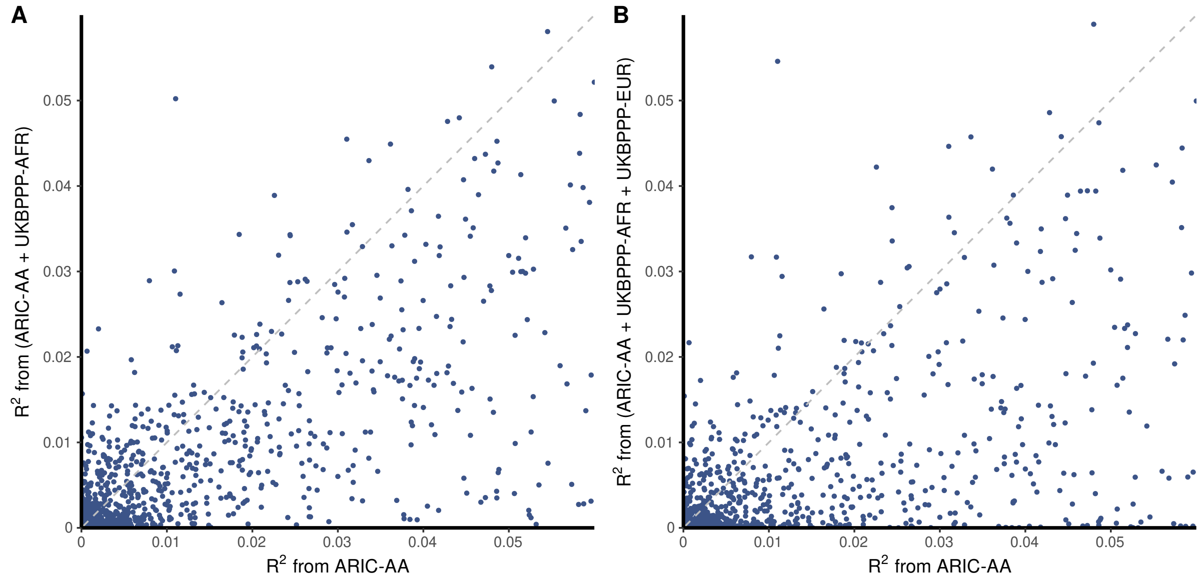


**Supplementary Figure 28: Comparison of BLISS-trained models under platform-mismatched scenarios.** Platform-mismatched scenarios were created by enhancing BLISS-trained ARIC African American (ARIC-AA; SomaScan) models with UKB-PPP African (UKB-PPP AFR; Olink) and UKB-PPP European (UKB-PPP EUR; Olink). (A) Compares the predictive performance of BLISS-trained ARIC-AA models (x-axis) and super learner-enhanced ARIC-AA models (with UKB-PPP AFR; y-axis). (B) Compares the predictive performance between BLISS-trained ARIC-AA models (x-axis) and super learner-enhanced ARIC-AA models (with UKB-PPP AFR and UKB-PPP EUR; y-axis). Each point represents a protein; dashed line represents $y = x$.

### 2 Comparative advancement of PWAS models derived by BLISS

The existing PWAS models, while foundational, were constrained by the available pQTL data and computational methods of their time. For example, PWAS models derived from the INTERVAL study [8] focused on European ancestry. These models were built by standard PWAS pipeline using N = 3,301 pQTL data measured by SomaScan platform, yielding 1,389 validated PWAS models with $R^2 \geq 0.01$ [12]. Another important resource comprises PWAS models derived from the ARIC study [10], which were built by applying standard PWAS pipeline to African American (N = 1,871) and European American (N = 7,213) pQTL data measured by SomaScan platform.

In comparison, the PWAS models generated using BLISS represent a substantial advance over these prior models in terms of ancestral diversity, scale, and proteomic platform coverage.

First, we provide PWAS models for European, African, and Asian ancestries, leveraging substantially larger and more diverse pQTL datasets. For European ancestry, we build PWAS models using three large pQTL datasets: UK Biobank Pharma Proteomics Project (UKB-PPP, Olink platform, N = 49,341 White British individuals), deCODE (SomaScan platform, N = 35,892 Icelanders), and ARIC (SomaScan platform, N = 7,213 European Americans). For African ancestry, we leverage both ARIC (SomaScan platform, N = 1,871 African Americans) and UKB-PPP (Olink platform, N = 1,181 African ancestry individuals). Furthermore, we introduce PWAS models for Asian ancestry using UKB-PPP data (Olink platform, N = 923 Asian ancestry individuals), an ancestry for which large-scale PWAS model resources were previously lacking.

Second, we provide PWAS models for both the Olink (antibody-based) and SomaScan (aptamer-based) platforms, thus expanding proteomic platform coverage. This expansion provides PWAS models for 1,024 proteins exclusively measured by the Olink platform, which were absent from SomaScan-based models built using deCODE and ARIC datasets. Furthermore, these new resources also enable systematic cross-platform validation and investigation of potential proteoform-specific effects for the 1,679 proteins common across all three European ancestry datasets (UKB-PPP EUR, deCODE, and ARIC-EA).

In summary, the BLISS resource provides a more comprehensive and powerful tool for multi-ancestry PWAS by substantially increasing the scale, ancestral representation, and proteomic breadth of available models.

### 3 Impact of PWAS model uncertainty on type I error rate and power

We examine how estimation uncertainty in PWAS models affects Type 1 error rate and power for PWAS framework. Following the derivation from [9], we show that under the valid instrument assumptions, estimation uncertainty in PWAS models primarily impacts statistical power while Type I error rates remain well-controlled.

#### 3.1 Model Framework

We denote the genotypes for $p$ $cis$-SNPs for the protein of interest as $\mathbf{G} \in \mathcal{R}^p$, with true LD matrix $\mathbf{\Sigma_{GG}} = \text{Cov}(\mathbf{G})$. Denote $X \in \mathcal{R}$ as the true expression level of the protein (which is unobserved in Stage II), and $Y \in \mathcal{R}$ as the trait of interest. We then consider a typical two-stage framework in PWAS:

1. **Stage I (protein expression prediction model):** The protein expression $X$ is genetically determined by $cis$-SNPs:

$$X = \mathbf{G}^T \boldsymbol{\gamma} + \xi, \tag{1}$$

   where $\boldsymbol{\gamma} \in \mathcal{R}^p$ is the vector of true genetic effects of SNPs on protein expression $X$, and $\xi$ is a non-genetic/environmental component with $\text{E}(\xi) = 0$ and $\text{var}(\xi) = \sigma_\xi^2$.

2. **Stage II (trait model):** The trait $Y$ is influenced by the protein expression $X$:

$$Y = \beta \cdot X + \varepsilon, \tag{2}$$

   where $\beta$ is the true causal effect of protein $X$ on trait $Y$, and $\varepsilon$ is an error term, potentially correlated with $\xi$, with $\text{E}(\varepsilon) = 0$ and $\text{var}(\varepsilon) = \sigma_\varepsilon^2$. Let $\text{cov}(\xi, \varepsilon) = \sigma_{\xi\varepsilon}$, which captures linkage effects or pleiotropy where genetic variants influencing $X$ (via $\xi$ being correlated with $\mathbf{G}^T\boldsymbol{\gamma}$ residuals if model is imperfect) also influence $Y$ through pathways not mediated by $X$. Notably, other covariates such as age, gender, population structure estimated from top principal components of the genotype matrix can be included in the above model, which has been ignored for derivation simplicity.

Substituting equation (1) into equation (2), we obtain the reduced form model for $Y$:

$$Y = \mathbf{G}^T(\beta \cdot \boldsymbol{\gamma}) + \beta \cdot \xi + \varepsilon. \tag{3}$$

The variance of the composite error term in this reduced model is $\sigma_t^2 = \text{var}(\beta \cdot \xi + \varepsilon) = \beta^2\sigma_\xi^2 + \sigma_\varepsilon^2 + 2\beta\sigma_{\xi\varepsilon}$. The PWAS approach first estimates $\boldsymbol{\gamma}$ using a pQTL dataset (sample size $n_1$) to get $\hat{\boldsymbol{\gamma}}$. Then, it tests the association between the genetically predicted protein expression $\hat{X} = \mathbf{G}^T\hat{\gamma}$ and $Y$ using a GWAS dataset (sample size $n_2$), effectively estimating $\beta$.

#### 3.2 Asymptotic Variance of estimated $\beta$

Suppose the pQTL dataset (Stage I) provides an estimate $\hat{\gamma}$ such that $\sqrt{n_1}(\hat{\gamma} - \gamma) \xrightarrow{d} N(\mathbf{0}, \boldsymbol{\Theta})$, where $\boldsymbol{\Theta}$ is the asymptotic covariance matrix of the scaled estimation errors of $\hat{\gamma}$, quantifying the uncertainty in the PWAS model. Here $\boldsymbol{\Theta}$ equals zeros when $\gamma$ is known without estimation error.

Next, we discuss two settings: (i) the setting where we have access to individual-level outcome GWAS data, and (ii) the setting where only summary-level GWAS data and a corresponding LD reference panel are available.

For the first setting, when $\hat{\beta}$ is estimated using individual-level GWAS data (Stage II) by regressing $Y$ on $\mathbf{G}^T\hat{\gamma}$, its asymptotic distribution is given by (following Proposition 1 of [9], adapted for no invalid IVs):

$$\sqrt{n_2}(\hat{\beta} - \beta) \xrightarrow{d} N(0, v), \tag{4}$$

where the asymptotic variance $v$ is:

$$v = \frac{\sigma_t^2}{\Psi} + \frac{n_2}{n_1} \cdot \frac{\beta^2 \cdot \phi}{\Psi^2}. \tag{5}$$

Here, $\Psi = \boldsymbol{\gamma}^T\boldsymbol{\Sigma}_{\mathbf{GG}}\boldsymbol{\gamma}$ represents the variance of the true genetically determined protein expression, and $\phi = \boldsymbol{\gamma}^T\boldsymbol{\Sigma}_{\mathbf{GG}}\boldsymbol{\Theta}\boldsymbol{\Sigma}_{\mathbf{GG}}\boldsymbol{\gamma}$ is the component of the variance attributed to the estimation uncertainty in $\hat{\boldsymbol{\gamma}}$.

For the second setting, where $\beta$ is estimated using summary-level GWAS data (Stage II) and an external LD reference panel of size $n_0$ is used to estimate the LD matrix $\boldsymbol{\Sigma}_{\mathbf{GG}}$ (denoted $\hat{\boldsymbol{\Sigma}}_{\mathbf{GG}}$), the estimator $\tilde{\beta}$ has an asymptotic distribution (following Theorem 2 of [9]):

$$\sqrt{n_2}(\tilde{\beta} - \beta) \xrightarrow{d} N(0, v_c), \tag{6}$$

where the asymptotic variance $v_c$ is:

$$v_c \approx v + \frac{n_2}{n_0}\beta^2\kappa. \tag{7}$$

Here, $v$ is defined in equation (5), and $\kappa$ is a term that captures the additional variance arising from the sampling error in the external LD reference panel. Notably, this additional variance component due to using an external LD panel is also proportional to $\beta^2$.

##### 3.3 Implications

The primary goal of PWAS is to test the null hypothesis $H_0 : \beta = 0$ (i.e., no association between the genetically predicted protein $X$ and trait $Y$). We next examine the structure of the asymptotic variances $v$ (equation 5) and $v_c$ (equation 7) under the null hypothesis ($H_0 : \beta = 0$) and note the following facts:

1. The term $\beta^2\sigma_\xi^2$ and $\beta\sigma_{\xi\varepsilon}$ in $\sigma_t^2$ become zero. So $\sigma_t^2 = \beta^2\sigma_\xi^2 + \sigma_\varepsilon^2 + 2\beta\sigma_{\xi\varepsilon}$ simplifies to $\sigma_t^2 = \sigma_\varepsilon^2$.
2. The second term in $v$, $\frac{n_2}{n_1} \cdot \frac{\beta^2 \cdot \phi}{\Psi^2}$, becomes zero because it is multiplied by $\beta^2$.
3. Similarly, the additional variance term in $v_c$ due to the use of LD reference panel, $\frac{n_2}{n_0}\beta^2\kappa$, also becomes zero.

Therefore, under the null hypothesis $H_0 : \beta = 0$, both asymptotic variances simplify to:

$$v_{\text{null}} = \frac{\sigma_\varepsilon^2}{\Psi} = \frac{\text{var}(\varepsilon)}{\boldsymbol{\gamma}^T \boldsymbol{\Sigma}_{\mathbf{GG}} \boldsymbol{\gamma}}. \tag{8}$$

This simplified variance under the null, $v_{\text{null}}$, does not depend on $\Theta$ (the uncertainty of $\hat{\gamma}$) nor on $\kappa$ (the uncertainty from the LD reference panel) beyond their presence in estimating $\Psi$ with $\hat{\Psi} = \hat{\boldsymbol{\gamma}}^T \hat{\boldsymbol{\Sigma}}_{\mathbf{GG}} \hat{\boldsymbol{\gamma}}$. Therefore, a standard Wald test statistic, $T = \hat{\beta}/\text{se}(\hat{\beta})$, where the standard error is based on an estimate of $v_{\text{null}}$, remains valid. This demonstrates that uncertainty in $\hat{\gamma}$ and $\hat{\boldsymbol{\Sigma}}_{\mathbf{GG}}$ does not inflate the Type I error rate of the PWAS association test.

Notably, the situation for the statistical power is different. When $\beta \neq 0$, uncertainty in estimating $\gamma$ (from pQTL data) and $\boldsymbol{\Sigma}_{\mathbf{GG}}$ (from a reference panel) may inflate the variance of $\hat{\beta}$ or $\tilde{\beta}$ through the additional terms in the asymptotic variance. This inflation may reduce the precision of PWAS models and consequently reduces statistical power compared to an idealized true PWAS model where $\gamma$ is known.

#### 4 Enhanced discovery via multi-ancestry meta-analysis

To quantify the power gained from a multi-ancestry approach, we performed a fixed-effects meta-analysis of PWAS results from the MVP dataset. We focused on 1,195,594 protein-phenotype pairs common to both European and African ancestry cohorts, spanning 875 proteins and 1,367 phenotypes. Our analysis demonstrates that a multi-ancestry framework enhances discovery, and that this enhancement is substantially amplified by incorporating our super learner-enhanced African models.

First, the multi-ancestry meta-analysis combining European BLISS results with super learner-enhanced African PWAS results identified 7,991 significant protein-trait associations ($P < 1.0 \times 10^{-4}$) involving 757 proteins across 894 traits at a $0.05$ FDR level (Table S3). Notably, 676 (8.5%) of these associations were not detected (FDR

$\geq 0.05$) by European-specific BLISS model alone, underscoring the increased discovery power provided by multi-ancestry meta-analysis.

Second, we investigate the value of the super learner-enhanced African models by comparing two meta-analyses: (i) A baseline meta-analysis combining European BLISS results with African standard PWAS results; and (ii) An enhanced meta-analysis combining European BLISS results with super learner-enhanced African PWAS results. The baseline meta-analysis identified 5,346 significant associations (FDR $< 0.05$). In contrast, the enhanced meta-analysis yielded 7,991 significant associations at FDR $< 0.05$, a 1.5-fold increase in discovery (Table S3). This demonstrates that the improved imputation accuracy of the super learner-enhanced models directly translates to a substantial statistical power gain in multi-ancestry meta-analysis.

In summary, these results suggest that a multi-ancestry meta-analysis of PWAS results, particularly when augmented with super learner-enhanced PWAS models, substantially increases discovery power.

### 5 Cross-platform comparison in protein–phenotype associations for African ancestry

We conducted a parallel cross-platform comparison for African-ancestry models, the Olink-based UKB-PPP AFR models and SomaScan-based ARIC-AA models. This analysis revealed comparable patterns of platform-specific discovery and cross-platform concordance as those observed in European ancestry analysis.

First, as observed in the European cohorts, proteins unique to each platform contributed substantially to the complementary discoveries. At a 5% FDR level, SomaScan-based ARIC-AA models identified 6,941 significant protein-phenotype associations, 6,221 of which were not detected by the Olink-based UKB-PPP AFR models. This set included 4,467 associations involving 657 proteins exclusively measured by SomaScan. Conversely, Olink-based UKB-PPP AFR models identified 2,063 significant protein-trait associations across 444 phenotypes (Fig. 4A). Among these, 1,343 significant associations were not identified by the SomaScan-based ARIC-AA models, including 986 associations involving 203 proteins uniquely measured by Olink.

For the 444 proteins common to both the UKB-PPP AFR and ARIC-AA models, we found that the Olink-based UKB-PPP AFR and SomaScan-based ARIC-AA models identified 2,530 ($P < 2.1 \times 10^{-4}$) and 1,094 ($P < 8.9 \times 10^{-5}$) significant protein-phenotype associations, respectively, at a 5% FDR level. UKB-PPP AFR models revealed 363 significant associations not detected by ARIC-AA. Conversely, ARIC-AA models identified 1,799 associations absent in UKB-PPP AFR. Similar to European-based findings, we observed strong and positive correlation in their association $Z$-scores. Among protein-phenotype pairs significant in either model, the cross-platform $Z$-score correlations were robust ($\rho = 0.57$, Fig. S25). When examining the overlap in significant findings, we found that of the 2,893 unique protein-phenotype associations identified by either model at an FDR $< 0.05$, 731 (25%) were significant in both. Notably, 631 of the 731 (86%) shared associations demonstrated consistent directions of effect (Fig. S26).

### 6 Platform consistency is critical in super learner-based PWAS model building

While our main analysis demonstrates that super learner can improve the predicitive accuracy of African PWAS models by integrating data from larger, platform-matched European PWAS models (Figs. 2B-2C and Fig. S10), it remains unclear whether super learner framework can benefit from integrating PWAS models from distinct platforms. To address this, we performed two experiments to evaluate cross-platform model integration through super learner.

First, we evaluated whether combining African PWAS models from different platforms was beneficial. We applied our super learner framework to integrate SomaScan-based ARIC-AA models with Olink-based UKB-PPP AFR models. This cross-platform ancestry-matched integration decreased the model performance, resulting in a 29% median decrease in predictive $R^2$ compared to using the single-platform UKB-PPP AFR models alone ($P < 2.2 \times 10^{-16}$, Wilcoxon signed-rank test; Fig. S28A).

We next investigated whether integrating large Olink-based UKB-PPP EUR models could overcome this negative effect. We applied the super learner framework to integrate Olink-based UKB-PPP EUR models plus SomaScan-based ARIC-AA models with Olink-based UKB-PPP AFR models. This complex integration failed to provide any benefit beyond within-platform (Olink-only) cross-ancestry super learner approach (Fig. S28B).

Collectively, these findings demonstrate that platform consistency is important for super learner to successfully leverage cross-ancestry PWAS models. Directly combining PWAS models from different platforms introduces heterogeneity that diminishes, rather than improves, predictive accuracy.

### 7 A practical guide to interpreting PWAS findings and integrating complementary evidence

#### 7.1 PWAS: context, strengths, and inherent limitations

PWAS identifies the association between the genetically determined component of protein expression and a phenotype. This approach is biologically motivated, aiming to connect proteins to traits through the genetic evidence.

Compared to conventional regional (or gene-based) genetic association tests, PWAS offers distinct advantages. While a mathematical linkage exists, as both can be viewed as assessing the aggregated genetic signals within a $cis$-region, PWAS directly tests the role of a specific protein's genetically regulated expression by using pQTL data to weight genetic variants.

Despite these strengths, the causal interpretation of PWAS findings faces inherent challenges. First, a PWAS signal may arise if its predictive variants (i.e., SNPs with non-zero weights in protein prediction models) are in linkage disequilibrium (LD) with the true causal variant for a neighboring gene or regulatory elements with direct effects on the phenotype [11, 3]. Second, the current coverage of the proteome is incomplete, meaning the genetically predicted level of a measured protein may be correlated with that of an unmeasured, yet truly causal, protein. Third, the predictive variants in PWAS models may influence the phenotype through pathways independent of the modeled protein, violating a key assumption of no horizontal pleiotropy. Therefore, a statistically significant PWAS result represents a robust starting point for hypothesis generation but should not, in isolation, be interpreted as proof of causality for tested proteins.

#### 7.2 Integrating PWAS with complementary analytical approaches

To construct a more compelling case for causality, it is essential to triangulate findings with evidence from other approaches, including statistical colocalization, fine-mapping, Mendelian Randomization (MR), and methods specifically designed for more robust causal inference in the context of TWAS/PWAS.

Statistical colocalization methods aim to determine the posterior probability that a GWAS trait signal and a molecular QTL signal (such as a pQTL) within the same genomic region are driven by the same causal variants. This provides a distinct line of evidence compared to PWAS. As elucidated by [3], PWAS identifies biomarkers whose genetically predicted component associates with phenotypes, whereas colocalization focuses on identifying specific shared causal variants. A strong colocalization result (e.g., PPH4 $> 0.8$) for a protein identified by PWAS significantly bolsters the evidence that the modeled protein mediates the GWAS signal through a shared genetic variant. However, the absence of strong colocalization does not necessarily invalidate a PWAS finding. Such discrepancies can arise if, for instance, the protein's regulation is polygenic, involving multiple small-effect $cis$-pQTLs that collectively contribute to the PWAS model but no single variant meets the stringent threshold. Our empirical drug validation analyses [6] support this, showing that a notable fraction (21 out of 69, 33.3%) of PWAS-identified drug-protein-phenotype trios lacked strong colocalization support (PPH4 $< 0.8$).

Fine-mapping techniques serve to identify a credible set of variants that are most likely to be causal for a given GWAS or QTL signal within a specific locus. Applying fine-mapping to both the pQTL signal and the GWAS trait signal can help ascertain whether their respective credible sets overlap or are in close proximity. This can also refine the set of specific variants driving the PWAS prediction model and allow an assessment of their plausibility as causal variants for the trait. Examples of fine-mapping methods include SuSiE, which has been applied in TWAS contexts [4]. However, fine-mapping has its own limitations. It assumes that causal variants are among those genotyped or well-imputed and requires accurate LD estimates from appropriate reference panels.

MR utilizes genetic variants as instrumental variables to infer the causal effect of an exposure (e.g., protein expression level) on an outcome (phenotype of interest). PWAS itself can be considered a form of $cis$-MR [5]. More formally, MR can be applied using specific, often lead or fine-mapped, pQTLs as instruments for the protein of interest. However, MR establishes causality under strong, and often untestable, assumptions: instrument relevance (robust association with protein expression levels), exclusion restriction (instruments affect the trait solely through the protein, precluding horizontal pleiotropy), and independence (instruments are independent of confounders of the protein-trait association). Violations of these assumptions, particularly horizontal pleiotropy, represent a significant challenge, and results can be highly sensitive to instrument selection. Various sensitivity analyses exist but also possess their own limitations.

Additionally, an evolving suite of methods aims to provide more robust causal inference within TWAS/PWAS contexts. These approaches often focus on dissecting horizontal pleiotropy or improving the identification of the most likely causal gene in regions of complex LD. Examples include methods that explicitly model and attempt to correct for pleiotropy (e.g., by adapting Egger regression principles or leveraging multi-SNP conditional analyses) or integrate information from multiple tissues or biological contexts to disambiguate signals [1, 2]. The 2ScML method [9] exemplifies an approach designed to handle invalid instruments in TWAS. As with other specialized techniques, these "robust" methods also rely on their own specific assumptions, which may not always be satisfied and are hard to verify, and their performance can vary depending on the nature of the confounding present.

In summary, no single statistical method provides irrefutable proof of causality in observational genetic data. PWAS, TWAS, colocalization, fine-mapping, and MR operate under distinct assumptions, and their findings can be influenced by numerous factors, including LD structure, allele frequencies, sample sizes, and the true underlying biological complexity. Researchers should maintain an awareness of the assumptions underpinning each method employed and critically assess their likely validity within the specific application. Thus, transparency in reporting the methods used, the assumptions made, and any limitations in the interpretation of results is critically important. By adopting this comprehensive and integrative framework, the PWAS models developed by BLISS can serve as a powerful resource for advancing our understanding of disease biology and identifying novel therapeutic strategies.

#### 7.3 A suggested workflow for prioritizing PWAS discoveries

We recommend a staged approach to move beyond initial PWAS discoveries. First, perform a rigorous proteome-wide scan using BLISS to identify an initial list of significant protein-phenotype associations, with appropriate correction for multiple testing (e.g., FDR or Bonferroni). Second, for these candidates, gather complementary evidence by performing colocalization, TWAS, statistical fine-mapping, and, where applicable, MR analyses using carefully selected instruments. Results from alternative "robust" TWAS/PWAS methods may also be included as sensitivity analyses. Third, synthesize all available evidence. High-confidence candidates will exhibit convergent support across multiple methods. Consideration should also be given to the novelty of the findings and their alignment with existing biological hypotheses, pathways, or clinical understanding. Discrepancies (e.g., a strong PWAS signal with weak colocalization) should be interpreted in light of each method's assumptions and limitations, such as the possibility of polygenic regulation. Fourth, the refined list of candidates should be formally experimentally validated to establish causality. Such studies might involve cellular or animal models to investigate the impact of altering protein expression or activity on disease-relevant phenotypes, as illustrated by recent work in pancreatic cancer [12].
